# Quantitative Imaging and Characterization of Ganglioside Diversity in Mouse and Human Brain Tissues

**DOI:** 10.64898/2026.05.20.726404

**Authors:** Shadrack M. Mutuku, Nicolas Tomasiello, Caine Smith, Greg Sutherland, Reza Ranjbar Choubeh, Maria José Q. Mantas, Lihang Yao, Nathan G. Hatcher, Nico Verbeeck, Shane R. Ellis, Kim Ekroos

**Affiliations:** Molecular Horizons and School of Science, Faculty of Health, Science and Medicine, University of Wollongong, Wollongong, NSW, Australia; New South Wales Brain Tissue Research Centre, Charles Perkins Centre, School of Medical Sciences Faculty of Medicine and Health, The University of Sydney, NSW, Australia; Aspect Analytics NV, Genk, 3600, Belgium; Merck & Co., Inc., 770 Sumneytown Pk, West Point, PA, 19486, USA; Lipidomics Consulting Ltd., Esbo, Finland

**Keywords:** gangliosides, mass spectrometry imaging, spatial lipidomics, brain imaging

## Abstract

Our understanding of how ganglioside species are spatially and quantitatively distributed within regions in mammalian remains limited. Studies are typically translated from rodents, assuming that gangliosides species contents within brain regions reflect the human condition. Herein, we provide a rich spatial ganglioside brain atlas describing the content, compositional differences, and concentrations of 48 ganglioside species across different regions in mouse and humans with no clinically known neurodegeneration. Our quantitative mass spectrometry imaging (MSI) approach allows non-rigid co-registration of mass spectrometry and microscopy images to allow precise spatial alignment and extraction of normalised ganglioside concentration data to reference mouse brain tissue anatomy and neuropathological annotated tissues for flexible downstream lipidomics analysis. Considerable differences and similarities are observed permitting region-specific ganglioside maps across twelve brain regions in wild type mouse. Gangliosides in mouse brain tissue generally preferred Cer 36:1;O2 configurations whereas human gangliosides in corresponding anatomically-annotated regions tended to favour Cer 38:1;O2 and longer backbones. Notably, this observation is more pronounced in gray matter compared to white matter. Collectively, this study defines the precise and quantitative ganglioside atlases across mice and human brains to guide and accelerate the discovery of biomarkers and therapeutic targets for brain diseases.

## Introduction

Gangliosides are highly complex amphiphilic glycosphingolipids found in plasma membrane domains in cells ^1, 2^ and are particularly abundant in the brain^3^. They consist of one or more sialic acid (SA) residues attached to an oligosaccharide structure conjugated to a ceramide (Cer) backbone. The predominant SA in humans is N-acetylneuraminic acid (Neu5Ac) ^4, 5^, whereas N-glycolylneuraminic acid (Neu5Gc) is common in other mammals, such as mice ^6^. Neu5Gc is not expressed in humans due to the absence of the cytidine monophosphate-N-acetylneuraminic acid hydroxylase, whose encoded protein adds a hydroxyl group to Neu5Ac. Ganglioside biosynthesis takes place in the Golgi apparatus ^7^ via a series of anabolic reactions starting from lactosylceramide (LacCer) and involves membrane-bound sialyltransferases and glucosyltransferases^8, 9^. Structural diversity arises from a variety of sources including the number and position of SA resides, the nature of the glycan, and length and degree of unsaturation in the acyl and amide chains of the ceramide backbone^10, 11^.

Gangliosides constitute at least 80% of all glycans in the brain and have important functions in brain maturation and neurological tissue development^3^, and are mainly found in neurons ^12^ and within the gray matter^13^. Enzymatic-driven changes to the levels of gangliosides have been linked to neurodegenerative conditions such as Alzheimer’s disease (AD)^14^ and Parkinson’s disease (PD)^15^. Deficiency in glucocerebrosidase (GBA)^16^ and scavenger receptor class B member 2 (SCARB2)^17^ genes impact lysosomal capacity to catabolize glucosylceramide leading to altered levels of the gangliosides and wider systematic changes that affect other glycosphingolipid metabolic processes that have been detected in cerebrospinal fluid and blood^13^. In PD-GBA patients (individuals carrying mutant GBA1), significant elevation of total gangliosides in the middle temporal gyrus, cingulate gyrus and striatum brain regions has been reported^18^. Hadaczek and colleagues reported significant decreases in GM1a, alongside its metabolic precursor GD1 levels in the occipital cortex and substantia nigra (SN) of PD patients^19^. This can be rationalized as denervation within the SN leads to a depleted neuronal mass and hence low ganglioside content. On the other hand, Gegg et al. reported that fatty acyl C18:0 GM2 and GM3 containing species were slightly elevated in the putamen and cerebellum of PD-GBA1 and sporadic PD patients compared to controls^20^. It could be considered that GM2 and GM3 accumulate at the expense of GM1 due to being upstream substrates of GM1. In the context of function, the decrease in GM1 would promote neurotoxicity^21^, as GM1 protects alpha-synuclein (ASYN) to maintain its non-aggregating alpha-helical and probable non-toxic conformation^22^ and promotes autophagy and autophagy-dependent ASYN clearance^23^.

Ganglioside determination has traditionally been conducted by 1D and 2D thin-layer chromatography (TLC), providing class-level contents^24, 25^. To resolve higher structural specificity, electrospray ionisation mass spectrometry (ESI-MS) has been applied, often coupled with liquid chromatography tandem mass spectrometry (LC-MS/MS)^26, 27^. Shotgun ESI has also been applied to quantitatively study ganglioside content differences of brain and optical nerve tissues in a murine model of GM3 and GM2 synthase genetic deficiencies^28^, as well as identifying the ganglioside signature of rat hippocampal synaptic junctions and membrane fractions^29^. Critically, given the divergent and complex nature of ganglioside structures, and their relationship to function, augmentation of LC-MS/shotgun lipidomics to other bioanalytical techniques to attain unambiguous identification is increasingly needed. Ion mobility separation (IMS) separates isobaric molecules based on collision-cross section area. For example, flow injection analysis combined with structures for lossless ion manipulation (SLIM) with high resolution IMS ^30^ has demonstrated separation of isomeric GD1a/b d36:1 isomers while field asymmetric IMS (FAIMS) has been used to detect up to 117 unique ganglioside species including, precursors containing five and six sialic acid moieties^31^. In both cases, SLIM and FAIMS have the added benefit of much reduced analytical times (2 – 6 min) compared to LC-MS workflows. Four dimensional reversed-phase LC-MS coupled to trapped ion mobility spectrometry TIMS and parallel accumulation serial elution fragmentation has detected 159 ganglioside entities in human serum and porcine brain extract ^32^. In general, these MS-based methods have enabled the characterization of ganglioside in a variety of complex samples prepared from tissue homogenates.

The above-mentioned bulk analyses do not provide any information on spatial localization, which is a prerequisite to understanding their precise physiological functions in disparate neurological anatomies. Recent studies report quantitative approaches utilizing laser capture microdissection (LCM)^26^ and other methods ^33, 34^. Although the adoption of LCM gives spatial information, it can still invariably dilute highly localized enrichments, as sampled areas typically range from several tens of micrometers up to a few millimeters in diameter. Mass spectrometry imaging (MSI) is a versatile label-free technique that measures the distribution of many biomolecules within tissue sections with single cell resolution. The most popular MSI applied to gangliosides has been matrix-assisted laser desorption/ionization (MALDI) ^35, 36, 37^, as well as desorption electrospray ionization (DESI) ^38, 39^. MALDI-MSI has been applied to study pathological changes during neurodegeneration in post-mortem human brain tissue^40^. In AD, increased GM3 was recently found to co-localize with amyloid beta plaques^40^. IMS has also been coupled with MSI to resolve isomeric species such as GD1a and GD1b^36^ and GM1 isomers^37^.

A general limitation of MSI is the inability to provide accurate spatial quantification of detected species throughout different tissue regions, largely due to analyte- and region-dependent ionisation efficiencies ^41, 42^ and ion suppression effects ^43^. To overcome this, quantitative MSI (Q-MSI) can be performed by applying an internal standard (often an isotopically labelled form of the analyte of interest) deposited over the sample at a known concentration and then performing pixel-by-pixel normalization^44^. However, these approaches have largely been targeted to a single or only on a handful of analyte(s) of interest^45,46^. Here, we have further adapted our previous broad Q-MSI approach^47^ to spatially quantify gangliosides in mammalian brain tissues. By employing this novel method, we present the first quantitative spatial maps of ganglioside species in human and rodent brains using MSI. We show the quantitative ganglioside landscape in twelve defined mouse brain regions, compared to human brain frontal cortex and midbrain tissues, as well as providing absolute concentrations, i.e. per pmol/mm^2^. The established atlases identify region specific variances across the mouse brain, differences between human and mouse, and dissimilarities in white and gray brain matters. These ganglioside atlases are critical to understanding the complex ganglioside metabolism in brain and offering new opportunities in guiding and accelerating neurodegenerative biomarker and therapeutic target discoveries.

## Results

### Ganglioside Q-MSI workflow

From the combined mouse and human sample data set, a ganglioside target list was created based on previous literature and using LIPID MAPS® Structure Database (LMSD)^48^ and GlyCosmos Glycolipids database (https://glycosmos.org/glycolipids). This initial list was first cross-checked against the MSI data using a 3-ppm mass tolerance to filter only those observed in the data. The final list of 48 unique gangliosides species across 18 classes reported herein was then created by applying a spatial filter followed by a manual curation step that kept only clear ion images that displayed consistent and structured distributions throughout at least one tissue type (mouse brain, human frontal cortex or human midbrain (see methods for further details). In parallel, we conducted orthogonal LC-MS/MS analyses on mouse total brain to further confirm the identification and presence of selected targets. From this we could confirm 45 species covering 16 classes, whereas three species belonging to three classes were annotated as tentative (see methods and *Supplementary File 1*)

Q-MSI was performed by depositing internal standards onto the tissue at known concentration (Figure 1) and then normalizing signals for endogenous gangliosides against the corresponding IS. As only GM3, GM1 and GD1 standards were commercially available at the time, precise quantification of GM3, GM1 and GD1 species was achieved, whereas for other ganglioside classes only estimations could be achieved by reference to the most suitable internal standard (see methods for further details). Given gangliosides primarily ionize on the sialic acid via deprotonation, the ionization efficiency is assumed to be similar.

**Figure 1.**
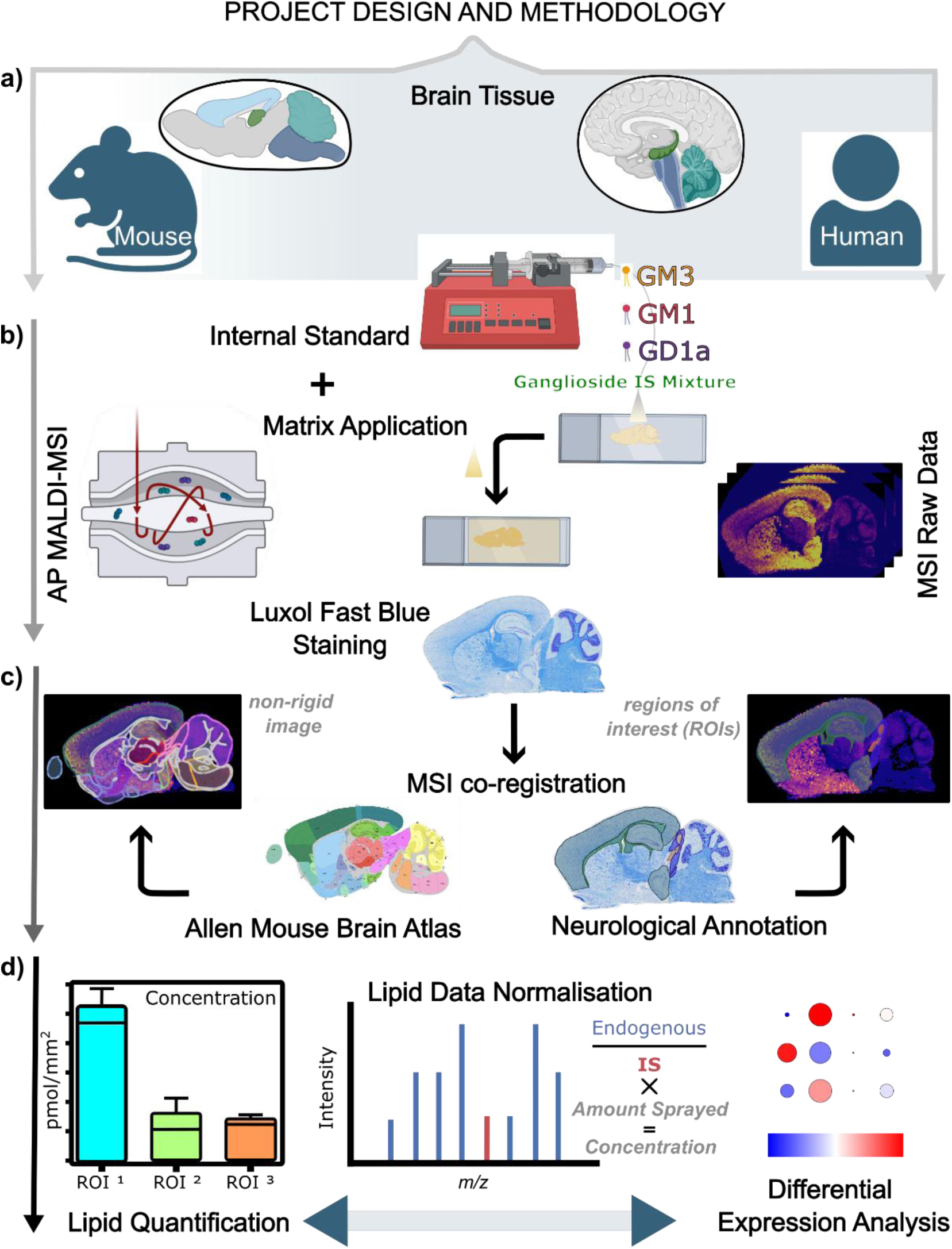
Quantitative Ganglioside MSI project design. **A)** Frozen tissue sections were collected from human frontal cortex and midbrain tissues with no neurological conditions, and wild type mice brain. **B)** High mass resolution ganglioside Q-MSI data was acquired by atmospheric pressure (AP) matrix-assisted laser/desorption ionisation on an Orbitrap Fusion instrument. **C).** MSI data underwent peak extraction, standard-based normalization with in-source fragmentation correction, and spatial filtering. Using post-MALDI Luxol fast blue microscopy, spatial co-registration was performed on MSI data by a non-rigid method to match morphological tissue features to the Allan Mouse Brain Atlas, or via manual annotation of regions of interest (ROIs). **D).** Extracted normalized intensities from pixels in each ROI were used to produce ganglioside amounts expressed as concentration per unit area (pmol/mm^2^) and subsequent differential expression analysis to quantify ganglioside content differences across tissue sections.

MALDI analysis of sialylated gangliosides is complicated by in-source fragmentation (ISF) leading to loss of sialic acid^49^. This is most significant for GM1 and GD1 species as they represent the most abundant gangliosides in brain and ISF of GD1 leads to artificial increases in GM1. While for GD1 this is accounted for by the IS normalization — as the standard also undergoes the same ISF — ISF of GD1 will lead to an over representation of GM1 concentration. Therefore, we developed a method to correct GM1 concentrations. This was achieved by calculating the extent of GD1 to GM1 ISF and then applying correction factors to remove the increased GM1 concentration arising from GD1 fragmentation (see details in methods). The effects of this on GM1 quantification are shown in *Supplementary Information Figure 1*. Notably, prior to ISF correction GM1 appears at higher concentration than GD1, however after correction GD1 is ∼ 20% higher than GM1, which is consistent with previous quantified amounts in brain recorded using LC-MS/MS ^18^ and shotgun lipidomics^28^, respectively.

*Figure 2A* shows a representative averaged mass spectrum from a mouse brain tissue within the *m/z* range where most gangliosides are found. Peaks assigned to different selected gangliosides are annotated. This shows high signal-to-noise values for many ganglioside species, with the most abundant signals arising from the 36:1:O2 and 38:1:O2 species of GM, GD and GT classes. These species agree with other studies such as Lee et al^34^. *Figure 2B-D* shows extracted peaks assigned to GM3, GM1, and GD1 species, respectively, with the corresponding IS peaks shown in red. The concentration of all curated gangliosides across all twelve mouse regions and in human frontal cortex and human midbrain specimens are provided in *Supplementary File 2*.

**Figure 2.**
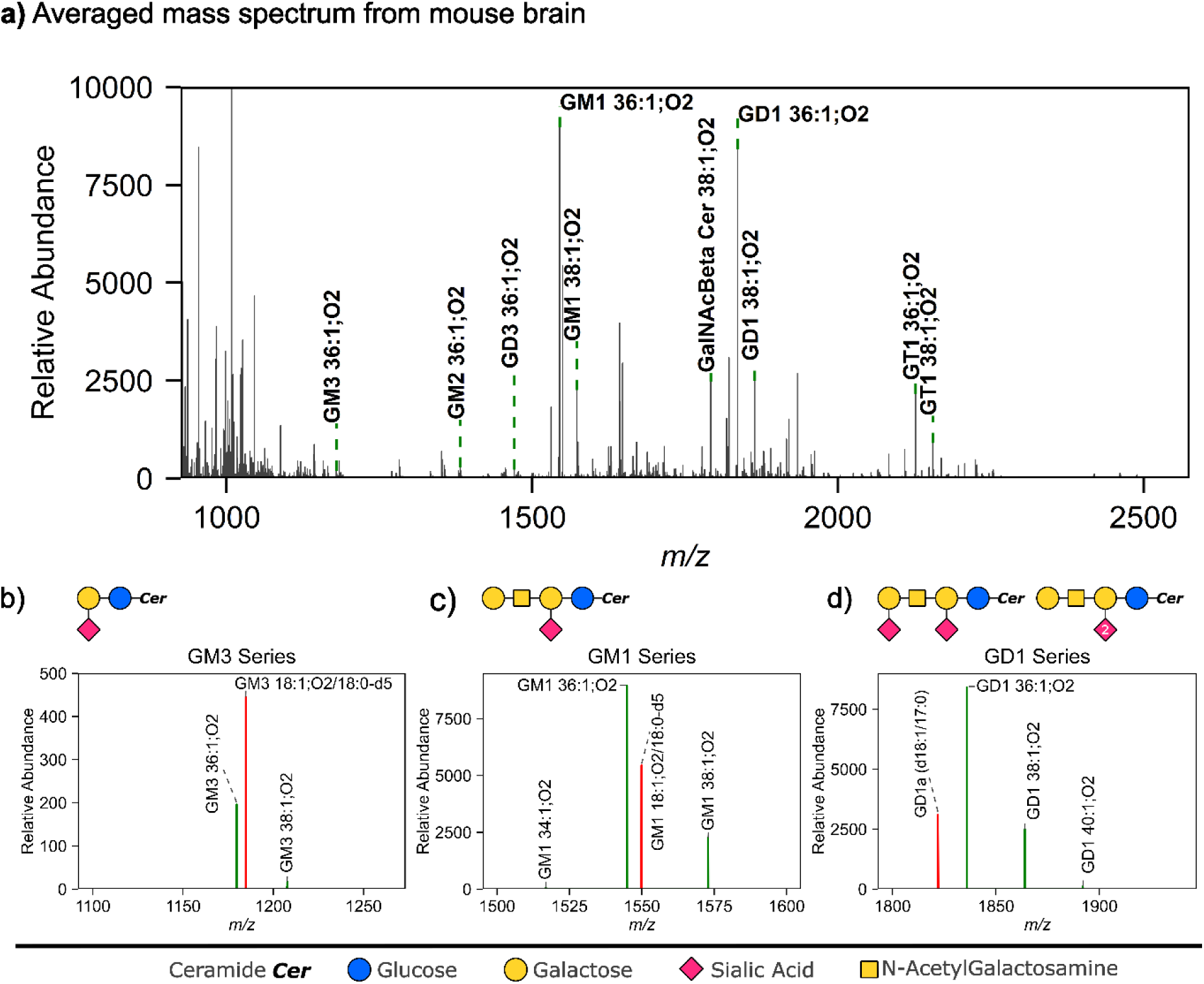
Principle of ganglioside Q-MSI. Top panel; A) Average mass spectra from a representative mouse brain sagittal tissue section across the *m/z* range where gangliosides are observed. Bottom panel shows extracted peaks assigned to B) GM3, C) GM1 and D) GD1 ganglioside species. Endogenous ganglioside peaks are shown in green while reference internal standard peaks for normalization are shown in red. Key for the ganglioside molecular structure is shown at the bottom; ceramide backbone is abbreviated as Cer; blue circle, glucose; yellow circle, galactose; pink diamond, SA (Neu5Ac) and yellow square, N-Acetylgalactosamine.

### Ganglioside Atlas of Mouse Brain

Using the Q-MSI workflow we first investigated the diversity and localisation of gangliosides in different mouse brain regions. We first compared 12 distinct regions of mouse brain anatomically defined by the Allen Mouse Brain Atlas (AMBA, *Figure 3A*). Here, the tissue sections were taken closest to sagittal level 10 corresponding to a lateral depth of 2.15 mm. MSI data was co-registered to the luxol fast blue (LFB) stained tissues which were then co-registered to the AMBA to enable extraction of region-specific mass spectra (see methods). *Figure 3B* shows total concentrations of 18 ganglioside classes across these 12 brain regions. Consistent with previous works^30^ the most abundant classes were GD1, GT1 and GM1. The hippocampal region generally exhibited the highest ganglioside concentrations across most classes including GM1, GD1, GT1 and GalNAcbeta with mean concentrations of 5.46±0.51, 21.54±2.30, 7.90±0.74, and 5.93±0.62 pmol/mm^2^, respectively. The cerebellar nuclei showed particularly low concentrations for GD2, GM2 and GT3 with concentrations of 0.008±0.008, 0.142±0.033 and 0.001±0.001 pmol/mm^2^ (*Figure 3B)*. A 2D-TLC analysis of seven-month-old mouse samples, also showed GD1a, GT1b and GM1, as the most abundant classes along with detection of O-acetylated analogues of these molecules ^25^. We do note that our data likely underrepresents classes like GT1 which undergo ISF but are not corrected here due to a lack of GT1 standard. Future work will incorporate additional standards now available to allow ISF correction of GT and GQ species.

**Figure 3.**
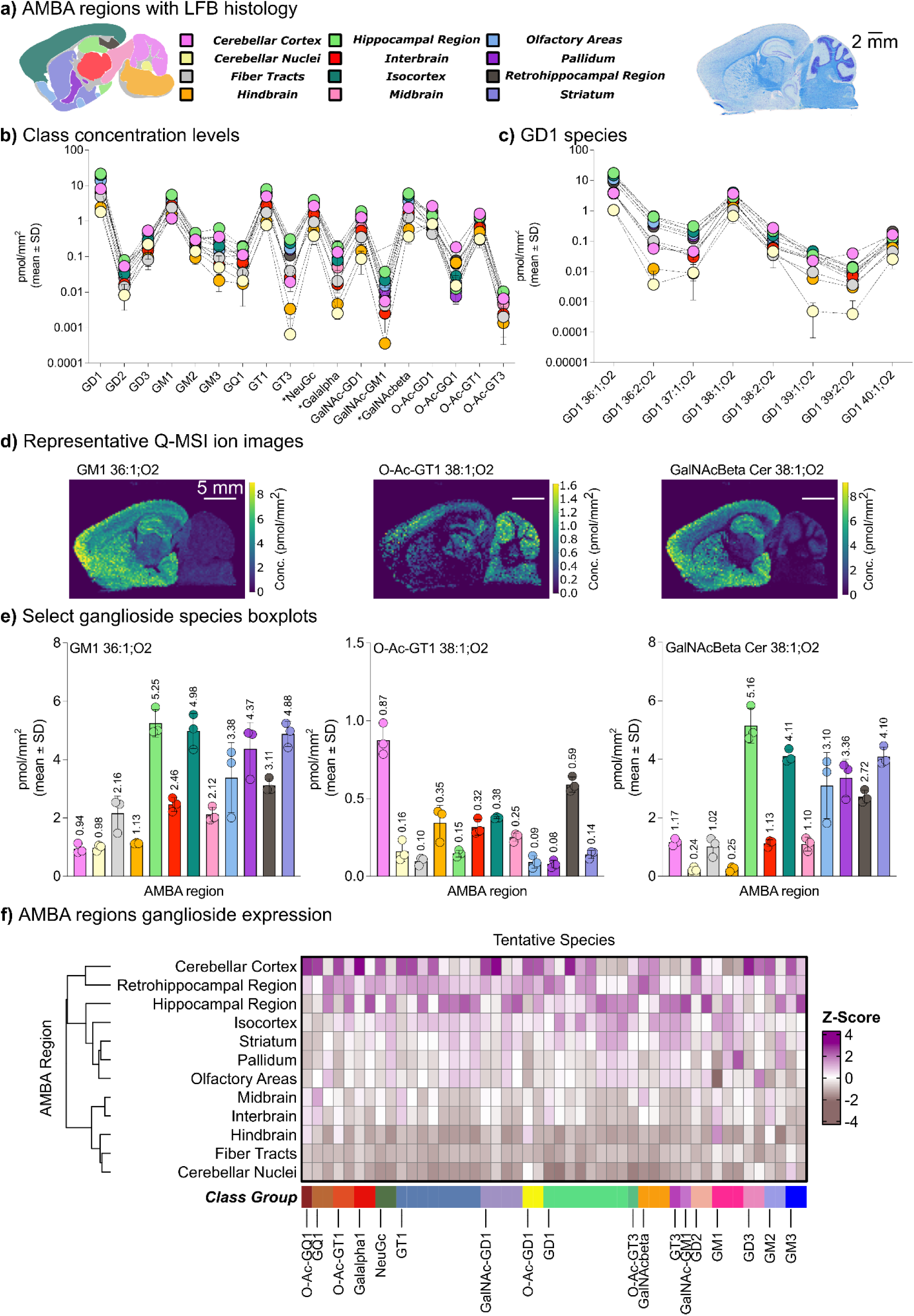
Comparison of ganglioside differences in AMBA regions. **A**) Allen Mouse Brain Atlas (AMBA) regions color coded for distinct anatomical regions in the mouse sagittal section. **B)** Average concentration of different ganglioside classes per AMBA region. **C)** Average concentrations of GD1 species per AMBA region. Circle colors correspond to the denoted AMBA region according to **A**. Results represented as mean pmol/mm^2^ ± SD (n=3). **D)** Quantitative ion images of GM1 36:1;O2, O-Ac-GT1 38:1;O2 and GalNAcbeta Cer 38:1;O2. Note that concentrations reported in ion images are prior to ISF correction of GM1. The corresponding concentrations are shown in **E)**. Results represented as mean pmol/mm^2^ ± SD. **F)** Clustering heatmap analysis of AMBA ROIs. The dendrogram on the left depicts levels of ROI groupings. Columns indicate clusters of 48 individual ganglioside species stratified into 19 classes. The Z-score scale bar indicates standardized normalized intensity values where each tile is the average, with magenta corresponding to high ganglioside enrichment and brown to low ganglioside enrichment. Lipid classes marked with * are shortened where NeuGc stands for NeuGcalpha2-3Galbeta1-4GlcNAcbeta1-3(GlcNAcbeta1-6)Galbeta1-4GlcNAcbeta1-3Galbeta1-4Glcbeta-Cer, Galalpha for Galalpha1-3Galalpha1-3Galbeta1-3GalNAcbeta1-4(NeuAcalpha2-8NeuAcalpha2-3)Galbeta1-4Glcbeta-Cer, and GalNAcbeta for GalNAcbeta1-4(NeuGcalpha2-3)Galbeta1-3GalNAcbeta1-4Galbeta1-4Glcbeta-Cer.

The lowest concentrations in the hippocampal region classes were observed for GD2and O-Ac-GT3, with O-Ac-GT3 observed at one-thousand-fold lower concentration than GD1. We found the lowest concentrations of gangliosides in cerebellar nuclei and hindbrain. 1.81±0.53 pmol/mm^2^ GD1 was determined in cerebellar nuclei which is about 12-fold less than in hippocampal region. Even bigger differences were observed for GT3, with hippocampal region containing about 0.31±0.01 pmol/mm^2^, and hindbrain and cerebellar nuclei containing only 0.0034±0.0015 and 0.00065±0.00113 pmol/mm^2^ respectively. Similar patterns were observed for other classes, such as GM3 and Galalpha (Galalpha1-3Galalpha1-3Galbeta1-3GalNAcbeta1-4(NeuAcalpha2-8NeuAcalpha2-3)Galbeta1-4Glcbeta-Cer). In contrast, the GM1 concentrations were much more similar across the regions, and for example, only 4.5-fold differences in GM1 concentrations between hippocampal region and cerebellar cortex. This shows that each region has its unique ganglioside content that can span wide concentration ranges, reaching up to a few thousand-fold between highest and lowest classes.

That each brain region consists of its own complex ganglioside profile species becomes more evident when analysing the individual species within each ganglioside class. *Figure 3C* shows the concentrations of different GD1 species within the 12 AMBA regions. GD1 36:1;O2 is most abundant with concentrations ranging from 1.06±0.23 to 17.39±2.10 pmol/mm^2^ in cerebellar nuclei and hippocampal region while GD1 36:2;O2 varies from 0.004±0.006 to 0.656±0.015 pmol/mm^2^ in cerebellar nuclei and isocortex. Another MALDI-MSI study of washing formalin-fixed fresh-frozen rat brain also reported shows a relative abundance of ∼3:1 for GM1 36:1;O2 and GM1 38:1;O2 in the striatum and cerebellar cortex^50^, consistent with our data. Interestingly, the odd-chain species GD1 39:1;O2 and GD2 39:2;O2, were dramatically reduced in cerebellar nuclei compared to the other regions, again demonstrating that each region is composed of unique ganglioside content.

*Figure 3D* shows the tissue distribution of selected gangliosides in IS normalized ion images where per pixel intensities are represented in pmol/mm^2^ (note these concentrations are prior to ISF correction which was performed for the regional analysis but not at the pixel level for MSI visualization). We observed that GM1 36:1;O2 is high in the hippocampal region, isocortex, pallidum and striatum whereas O-Ac-GT1 38:1;O2 is more localized to cerebellar cortex and retrohippocampal region. GalNAcBeta Cer 38:1;O2 displays a similar distribution to GM1 36:1;O2 except for the cerebellum’s lobule granular layers. The concentration breakdown of these three species across the 12 AMBA regions shown in *Figure 3E* illustrates how the species are quantitatively distributed. We further performed hierarchical clustering analysis on the complete ganglioside data set to identify unique signatures of ganglioside expression across AMBA regions (*Figure 3F*). Expectedly, the cerebellar cortex showed the highest enrichment levels. This was followed by the cluster of hippocampal and retrohippocampal regions, and two sub clusters of isocortex, striatum and pallidum, and olfactory areas. The regions with overall lowest ganglioside concentrations were two sub clusters of midbrain, interbrain and hindbrain, fiber tracts, and cerebellar nuclei, as these are largely comprised of white matter (WM) rather than the gray matter (GrM) that is rich in neurons. We note that for these regions, signal to noise of the lower abundance gangliosides means relative differences within these regions are estimates only and should be interpreted with caution.

### Evaluation of Ganglioside Quantities and Composition in Mouse and Human Brain Tissues

Having established the quantitative spatial map of ganglioside in mouse brain we next sought to compare the mouse AMBA-referenced ganglioside composition maps to that present in human midbrain and frontal cortex tissue as these brain areas lie at the root of numerous brain disorders, including AD ^51^ and PD ^52^. Given the sampling variability amongst the human tissues, and the dramatic differences observed between white (WM) matter and gray matter (GrM) regions in mouse tissues, we compared corresponding WM and GrM regions between the mouse and human frontal cortex, and WM regions in the midbrain (noting that only two of the midbrain sections had clearly defined GrM regions). This was achieved by histological annotation of LFB-stained tissue sections whereby only regions that could confidently be assigned were used for subsequent analysis. *Figure 4A* shows optical images of representative tissues for mouse, human cortex and human midbrain, while the full set of Q-MSI *m/*z images are provided in the *Supplementary Information*.

**Figure 4.**
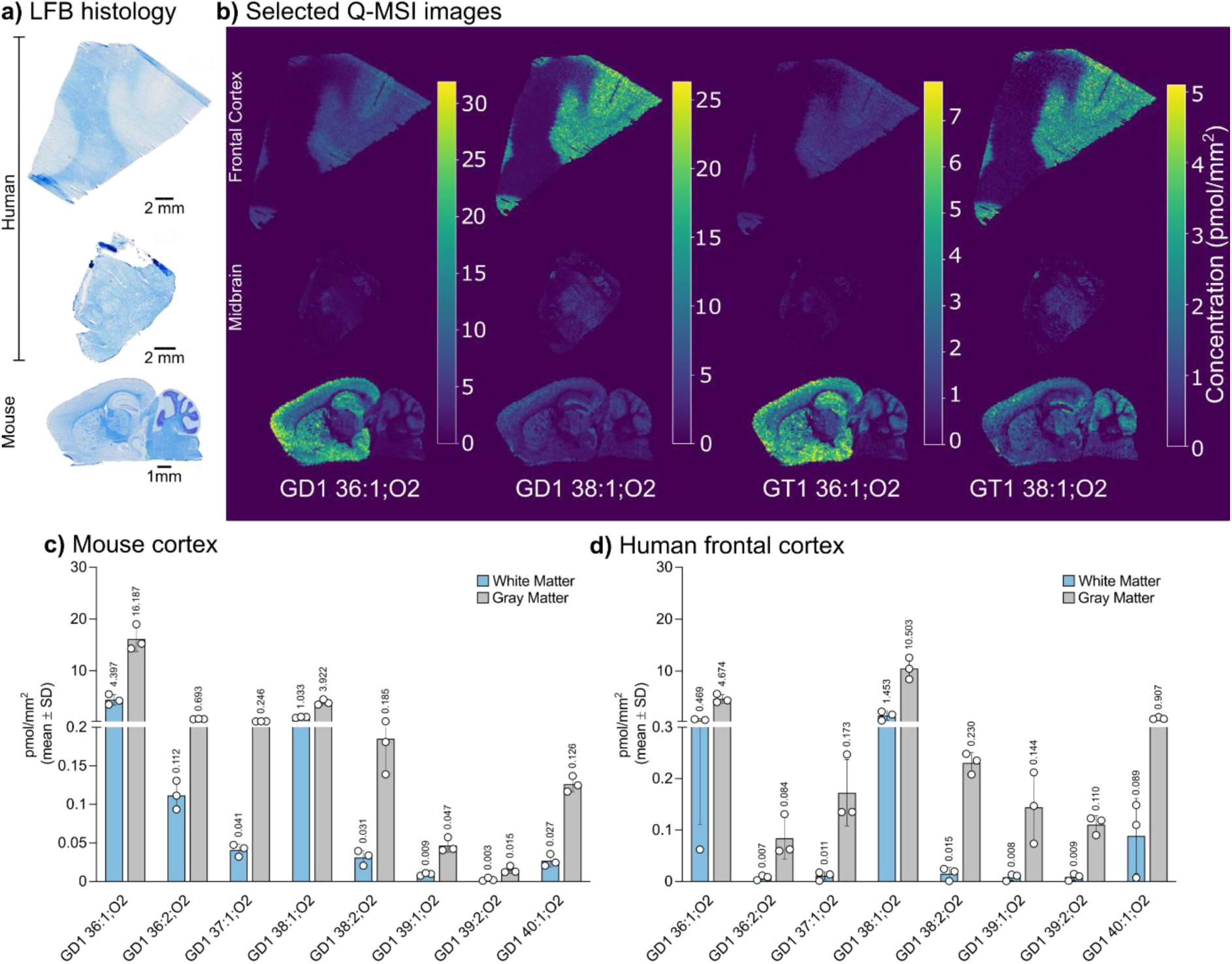
Ganglioside quantitative profiles between human and mouse brain tissues. **A)** Representative LFB-stained tissues of human frontal cortex, human midbrain and sagittal mouse brain acquired after MSI. **B)** Quantitative ganglioside ion images showing differences in localization and concentration of selected ganglioside species, GD1 36:1;O2, GD1 38:1;O2, GT1 36:1;O2 and GT1 38:1;O2. Bar plots showing comparison of GD1 species amounts between gray matter (GrM) and white matter (WM) regions are shown in **C)** for mouse cortex and **D)** for human frontal cortex (n=3 per sample).

. For the main ganglioside species belonging to GD1 and GT1 classes, concentration levels were between 5 – 30 pmol/mm^2^ dependent on mammalian origin and brain area. The quantitative images of selected gangliosides across representative human and mouse tissues provided in *Figure 4B* demonstrate a clear correlation between distinct tissue types. Similar to mouse, the human GrM regions exhibited higher GD1 and GT1 concentrations with significant lower concentrations in the WM. This is particularly apparent in human frontal cortex tissues that have larger, layered GrM areas while the midbrain samples were predominantly composed of WM. Guided by LFB histology, we could precisely define the GrM and WM borders in both mouse and human tissues (*Figure 4C*) and assess the compositional differences between mouse and human. The differences between tissue types are most striking when comparing the quantitative GD1 profiles across WM and GrM regions of mouse cortex and human frontal cortex, respectively *(Figure 4C* and *4D*). To contextualize the translational significance of our findings, we investigated the holistic ganglioside compositional differences between mouse cortex and human frontal cortex.

Of the 48 gangliosides measured across the GrM of mouse and human frontal cortex, we identified 17 individual species to be significantly (p-value <0.05) more abundant in mouse cortex, while 14 species were higher in human frontal cortex (*Figure 5A*). This demonstrates unique ganglioside composition quantities between mouse cortex GrM and human frontal cortex GrM. This was further explored in *Figure 5B,* showing the correlations between measured ganglioside concentrations in human and mouse GrM cortex. Species above the dotted line are relatively more concentrated in human tissue, and those below the line found at higher concentration in mouse. Here, GD1 38:1;O2 (10.5±2.1 pmol/mm^2^) is the most concentrated in human followed by GalNacBeta Cer 40:1;O2 (2.6±0.5 pmol/mm^2^) and GM1 38:1;O2 (1.9±0.7 pmol/mm^2^). In contrast, GD1 36:1;O2 (16.1±2.5 pmol/mm^2^) followed by GM1 36:1;O2 (5.1±0.6 pmol/mm^2^) and GalNacBeta Cer 38:1;O2 (4.6±0.6 pmol/mm^2^) are most abundant in mouse. On the other hand, when comparing low concentration species we find that GD1 40:1;O2 (0.9±0.2 pmol/mm^2^) and GM2 38:1O2 (0.6±0.1 pmol/mm^2^) are associated with human frontal cortex whilst O-Ac-GT1 36:1;O2 (1.0±0.2 pmol/mm^2^), GT1 38:4;O2 (0.6±0.1 pmol/mm^2^) and GD1 36:2;O2 (0.7±0.1 pmol/mm^2^) are elevated in mouse cortex (*Figure 5B, inset*). Irrespective of concentration and ganglioside class, a common differential factor between mouse and human is, as observed above, the higher 38:1;O2 to 36:1;O2 ratio in human^50, 53^. However, there are some exceptions like GalNacBeta Cer 38:1;O2 being more abundant in mice brain. The Q-MSI images highlighted in *Figure 5C* serve as an example of concentration differences exhibited by the unique ganglioside localizations and quantities between human frontal cortex and mouse brain tissues. Comparing short to long chain ceramide backbones of all summed gangliosides reveals that d36, d38 and d40 are the most predominant (*Figure 5D*). In addition to the observations in cortex GrM, d34, d36 and d37 ganglioside species in mouse cortex relative WM are increased 6.5-, 8.0-, and 3.2-fold, respectively relative to human cortex WM. Conversely, the comparative d40 and d42 species levels in human cortex relative to mouse cortex GM are 0.3- and 0.1-fold lower, respectively. Additional comparisons between human and mouse regions are provided in *Supplementary Information Figure 2*. Collectively, these data reinforce the inter-species differences in the brain glycosphingolipidome.

**Figure 5.**
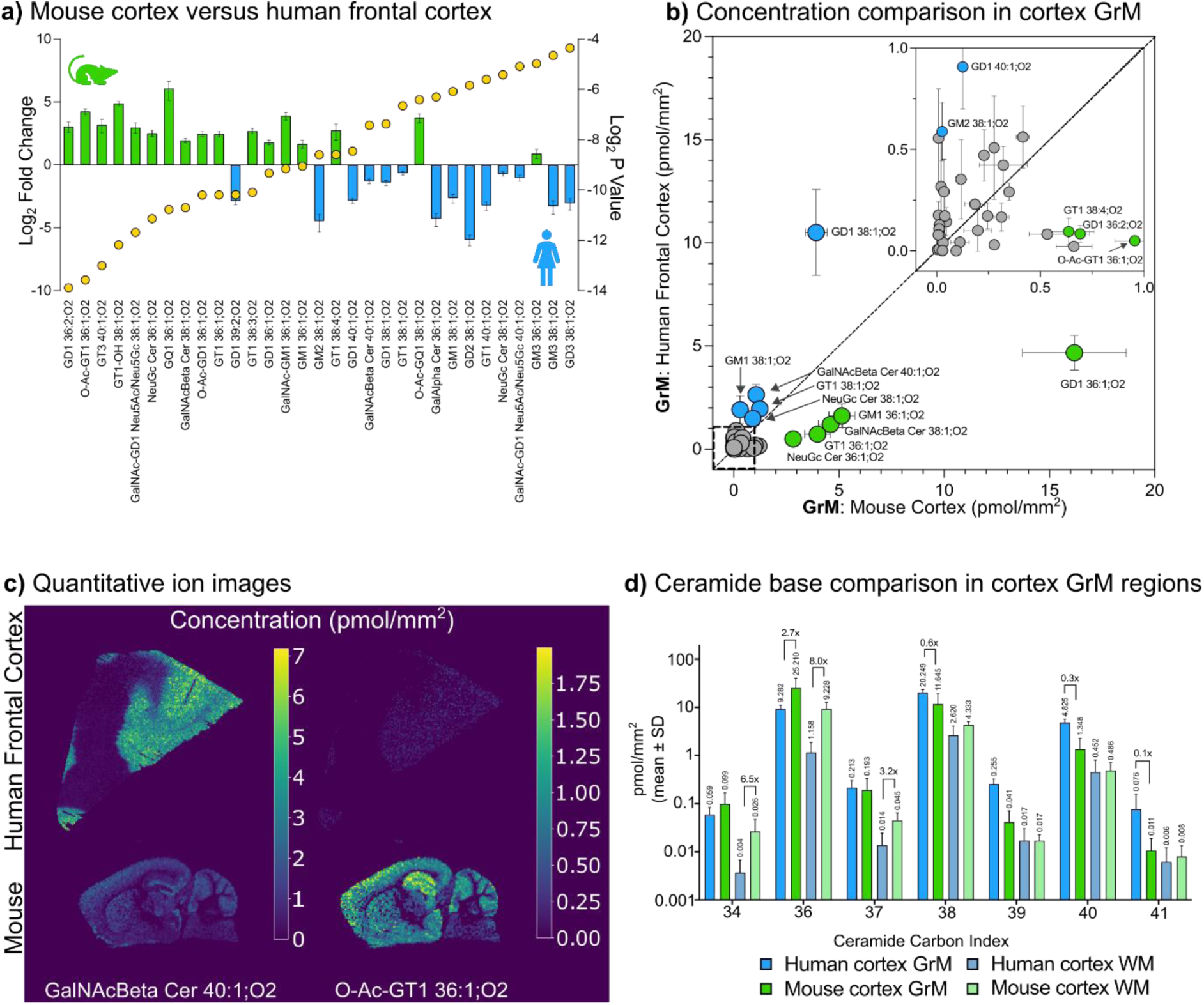
Ganglioside quantitative profiles between human and mouse brain tissues. **A)** Fold change bar plots of ganglioside concentration in gray matter (GrM) ROIs. Top bars show species measured in higher amounts in mouse cortex relative to human frontal cortex, while bottom bars show species in measured in higher amounts in human frontal cortex relative to mouse cortex, as indicated by log_2_ fold change, left y-axis. **B)** Correlation scatter plot of ganglioside concentrations between GrM of human frontal cortex and mouse cortex. Blue and green species correspond to those shown in A. Grey species are those with no significant difference **C).** Representative ion images for differentially concentrated ganglioside species that show that GalNAcBeta Cer 40:1;O2, left, is more pronounced in human frontal cortex while O-Ac-GT1 36:1;O2, right, is higher in mouse brains. **D)** Comparison of ceramide backbone carbon index for GrM and white matter (WM) regions of human frontal cortex and mouse cortex. Differences are marked by vertical annotation bars.

The identities of representative abundant even chain species, and the unusual, odd chain species, were further supported with tissue MS/MS (*Supplementary Information, Figure 3A, 4A*). The key diagnostic product ions are consistent with previous reports of CID/HCD analysis in porcine brain^32^. Moreover, at the lower mass range, our data (*Supplementary Information Figures 3B, 4B*) reflects CID studies of ceramide by ESI-MS reported by Hsu and Turk^54^. Thus at MS/MS level, we were able to deduce that GM1 36:1;O2 *m/z* 1544.87, GD1 36:1;O2 *m/z* 1835.95, and GT1 36:1;O2 *m/z* 2127.06 were predominantly composed of Cer d18:1/18:0, while GM1 38:1;O2, *m/z* 1572.90 GD1 38:1;O2 *m/z* 1863.99 and GT1 38:1;O2 *m/z* 2155.09, were predominately composed of Cer d20:1/18:0. We also report the fragmentation profile of an odd-chain species GD1 37:1;O2 (*Supplementary Information Figure 5*). These findings are in good alignment with previous studies. For example, Huang et al showed by LC-MS/MS a higher ratio of 38:1;O2 to 36:1;O2 in the main gangliosides in human brain^53^, while this ratio was opposite in mouse brain. Noteworthy, as they measured whole brain mouse tissue with a different genetic background, collectively their results and ours suggests that gangliosides with 36:1;O2 backbones are predominant to mice in general. In line with our results, Huang et al identified 38:1;O2 to be the main ganglioside ceramide backbones in human dorsolateral prefrontal cortex^53^. Rosenberg and Stern already showed that during brain maturation there is a shift from 18:1- to 20:1-sphingosine that seems necessary for healthy brain development^55^. Identifying 20:1-sphingosine based gangliosides can therefore be expected, however why this is predominantly found in humans and less in mouse cortex is unclear. In addition, our findings further suggest that mouse cortex is richer in O-acetylated species that have been implicated to play key roles in neuronal development by altering the sialic acid properties^6^. We did not detect any O-Ac-GD3, which is likely due to using wild type mice and type mice and healthy humans rather than under O-Ac-GD3 stimulated conditions such as neuronal regeneration^56^ and apoptosis ^57^.

We finally performed a comprehensive comparison of all gangliosides and across all analyzed WM and GrM tissue regions (note that for midbrain tissues only two sections had annotated GrM regions). The complete concentration tables are provided in the *Supplementary Information File 2*. Clustering heatmap analysis demonstrates distinct ganglioside compositional signatures for each sample type measured (*Figure 6A*). The highest ganglioside expression is as expected in GrM, however, and as shown above, the human frontal cortex has a different ganglioside makeup than that found in mouse cortex. In addition, the content of mouse midbrain GrM is different to mouse cortex GrM, showing more pronounced concentrations of GQ1 and O-Ac-GQ1, and several less abundant species (*Figure 6A*). Overall, homing in on the region-specific differences of human frontal cortex and midbrain tissues, the GrM still overwhelmingly appears to have higher total amounts of ganglioside for most classes compared to WM (*Figure 6B, 6C*). Diving deeper at the individual species amounts of the human gray matter, we see GQ1 36:1;O2 as the only ganglioside that potentially occurs in larger amounts in the midbrain compared to the frontal cortex (*Figure 6D*). Although less gangliosides are expressed in WM regions, similar trends are observed for GrM. Collectively, the results show that ganglioside contents are dependent on type of mammalian species and selective region. All Q-MSI images across mosue and human tissues are provided in *Supplementary Information Figure 6*.

**Figure 6.**
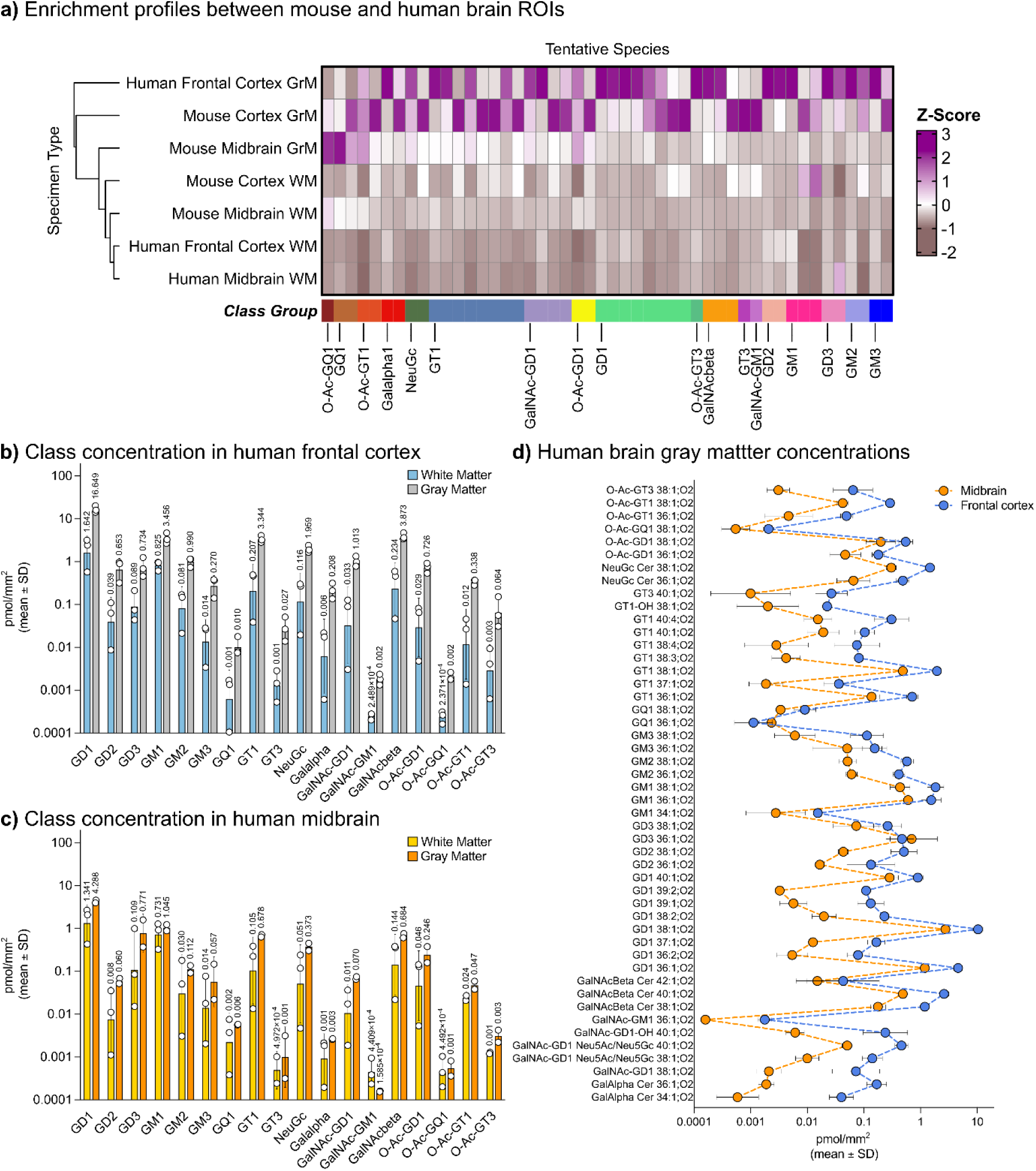
Ganglioside class profiles between human and mouse brain tissues. **A)** Hierarchical clustering heatmap of regions from gray matter (GrM) across human frontal cortex and mouse brain tissues, and white matter human frontal cortex, human midbrain and mouse tissues (n=3 for each). The similarities of the regions are shown by the dendrogram on the left, specimen type. Columns indicate clusters of 48 different ganglioside species grouped in 18 classes annotated by distinct colors. The Z-score scale bar indicates standardized normalized intensity values where each tile is the average. High ganglioside enrichment denoted in magenta and low ganglioside enrichment in brown. Data is the mean of 3 observations except for human midbrain GrM which was only confidently annotated in two of the samples. **B)** Comparison of total concentration across 18 different ganglioside classes between human frontal cortex WM, blue bars, and GrM, grey bars. **C)** Comparison of total concentration across 18 different ganglioside classes between human midbrain WM, yellow bars, and GrM, orange bars. Results represented as mean pmol/mm^2^ ± SD. **D)** Individual ganglioside species concentrations between GrM regions of human midbrain, (orange) and human frontal cortex GrM regions (navy dots). Results represented as mean mol% ± SD. Note that for human midbrain only two of three samples had regions that were confidently annotated as gray matter, all other regions represent n=3.

## Discussion

Gangliosides serve pivotal roles in mammalian neuronal development ranging from signal transduction, endocytosis, membrane trafficking, and synaptic plasticity ^58^. They are specifically abundant in neurons and glia where they can undergo dramatic changes during cell development and can account for 10–12% of the total brain lipid content ^59^. Since discovered back in 1942 by Ernst Klenk ^60^, studies have underpinned their involvement in multiple neurological disorders ^15, 17, 61, 62, 63^. Around 200 species have been identified ^63^, yet our understanding of how the ganglioside species are quantitatively distributed within brain regions has remained limited. This restriction has hindered comprehensive comparisons between different mammalians, challenging identification, and translation of preclinical models that mirror human physiology. Here, we have addressed these challenges by establishing an analytical platform for the accurate spatial quantification of ganglioside species with the creation of adult wild-type mouse and healthy human brain region-defined reference libraries. In contrast to previous studies that were based on quantitative analysis of bulk extracts, or MSI in the absence of internal standards for quantification, our approach provides the quantitative spatial ganglioside fingerprint of individual brain regions. Applying this method, we accurately fingerprinted the quantitative ganglioside landscape, covering in total 48 individual species across 18 classes, from 12 anatomically AMBA defined regions of eight-month-old wild-type mice and two regions of neuropathologically normal aged humans.

To minimize false identifications, we thoroughly iterated our targeted ganglioside list to match literature and databases, including applying stringent MSI analysis thresholds. We conducted orthogonal LC-MS/MS analyses on total mouse brains to further verify the identifications. In most cases we could verify the identities, however we could not confirm the GalNAcbeta1-4(NeuGcalpha2-3)Galbeta1-3GalNAcbeta1-4Galbeta1-4Glcbeta-Cer and NeuGcalpha2-3Galbeta1-4GlcNAcbeta1-3(GlcNAcbeta1-6)Galbeta1-4GlcNAcbeta1-3Galbeta1-4Glcbeta-Cer classes. Thus, species belonging to these classes remain tentative, and future work is warranted to confirm these species by synthetic standards. Collectively, this not only sets the foundation to comprehensively study the spatial organization of gangliosides but also unlocks the future studies of uncharted species.

Several studies, irrespective if by TLC^25^ or MS^28^, find GD1 and GT1 the most abundant ganglioside classes in the mouse brain. However, how the gangliosides are distributed across brain-regions has remained limited due to analytical hurdles. With our Q-MSI approach, we observe good concordance in the region defined ganglioside identities deciphered by Lee and colleagues^34^ analyzing isolated tissue regions by LC-MS/MS, with 36:1;O2 and 38:1;O2 being the main chain compositions. In contrast, our study is the first to determine the quantities expressed in pmol/mm^2^, of individual gangliosides within precisely defined regions. This is achieved by spraying defined amounts of internal standards onto the tissue prior to the MS analysis. Since most previous studies report concentrations in extracts (*e.g.* pmol/mg protein or pmol/g wet weight) it is challenging to compare such results to ours. However, if we generalize that a 12 µm thick slice covering a 1mm^2^ area to weigh 0.012 mg (1mg/mm^3^), the concentration of GM1 in the hippocampal region would approximately correspond to 450 pmol/mg. Intriguingly, this closely aligns to the amounts previously obtained in hippocampus^64^ (300 pmol/mg) using sialic acid quantification on isolated tissues. This indicates that our Q-MSI approach offers the advantages of reliable quantitative comparison of individual gangliosides across brain regions and sample types, in addition to minimizing potential sampling errors as no physical dissection of individual brain regions is needed. Thus, this work reports holistic brain ganglioside atlas unlocking new avenues to understand the highly complex ganglioside biology down to the individual species and with their precise localizations. We note that the current MSI data is acquired at 100 µm and was practical choice given the size of the human sections and the use of a slow but high mass resolving power Orbitrap mass spectrometer. This means that individual pixels and thus annotated ROIs contain a mixture of cell and/tissue types. Moreover, given the diversity of the individual human samples post-MSI LFB staining did not allow confident assignment of WM and GrM regions throughout the whole human tissue and thus only regions that were confidently annotated were used. Future work will also aim to apply higher speed and higher resolution MSI technologies to understand ganglioside changes throughout the brain with single cell resolution and cell type specificity throughout the whole tissue.

Given the important role of gangliosides in neurodegenerative diseases, it is essential to understand how well the mouse data translates to humans. The distinct differences observed between equivalent regions of human and mouse tissue are in-line with previous reports demonstrating a preference for 38:1:O2 gangliosides in humans and 36:1:O2 in mouse. Our data provides a preference for 20:1;O2 sphingosine bases in humans whereas in mouse 18:1;O2 dominates. As palmitate and stearate are both abundant in mouse^65^ and human^66^ brain, future studies are warranted to elucidate this selectivity including biological role. In humans, the preferences are not only for longer long-chain bases, but also very-long chained fatty acids above C20. In contrast, O-acetylated gangliosides are more common to mouse, with O-Ac-GD1 36:1;O2 and O-Ac-GT1 36:1;O2 being five- and ten-fold more concentrated in the cortex gray matter. The differences extend beyond these given examples, not only highlighting configurational and species distinctions but also inequalities between regions. Importantly, our work defines the expected regional ganglioside landscapes in normal healthy mammalians, providing an invaluable resource for future preclinical model optimizations closer to mimicking humans and studying neurodegenerative diseases.

Building on our previous work^47^ we extended the coverage of our Q-MSI platform to include complex gangliosides. The developed platform is versatile to which additional internal standards can be added to further extend class specific quantification and broader correction for in-source fragmentation. The current work demonstrates brain region specific ganglioside fingerprints across mammalian species, creating critical local libraries to understanding the complex ganglioside metabolism and interconnected network in brain. Future work will include expanding regions of interest, determining the spatial ganglioside defects in neurodegenerative diseases, and defining the precise ganglioside changes during aging^67^. In summary, this work offers new opportunities in studying ganglioside biology, guiding and accelerating neurodegenerative biomarker discovery and therapeutic targets, and overall healthy aging.

## Acknowledgments

This work was supported by the Michael J Fox Foundation (grant numbers MJFF-022753 and MJFF-019154) and the Australian Research Council (LE250100150). Tissue samples were obtained from the Queen Square Brain Bank, and we gratefully acknowledge the QSBB staff for their support in tissue collection and provision. The Queen Square Brain Bank is supported by the Reta Lila Weston Institute of Neurological Studies, UCL Queen Square Institute of Neurology. The authors acknowledge the facilities, the technical and scientific assistance of the Imaging Facility, Faculty of Science, Medicine and Health, University of Wollongong.

## Data Availability Statement

Mass spectrometry imaging data (.imzML) from both human and mouse tissues can be downloaded from Zenodo (doi: 10.5281/zenodo.20302554) and will also be provided by the corresponding authors upon request.

## Materials and Methods

### Chemicals and Reagents

LC-MS grade water, gradient grade ethanol and analytical grade ammonium sulfate were purchased from Merck Life Science (Truganina, VIC, Australia). 2,5-dihydroxyacetophenone (DHA, 98.0% purity) was purchased from AK Scientific Inc (Union City, CA, USA). Gangliosides standards C18:0 GM3-d5 (synthetic) [860073W-100UG], C18:0 GM1-d5 (synthetic) [860076W-100UG] and C17:0 GD1a (d18:1/17:0) [860075W-100UG] were provided by AVANTI Polar Lipids (Alabaster, Alabama, USA). The exact concentration of the IS mixture is provided in Table 1.

**Table 1.** Ganglioside composition of the internal standards.

| Internal standard | Concentration<br>(mg/mL) | Adduct | *m/z* | Picomoles/mm<sup>2</sup> |
| --- | --- | --- | --- | --- |
| C18:0 GM3-d5 (synthetic) | 0.105 | [M-H]<sup>-</sup> | 1184.7686 | 0.41 |
| C18:0 GM1-d5 (synthetic) | 0.094 | [M-H]<sup>-</sup> | 1549.9008 | 1.86 |
| C17:0 GD1a (d18:1/17:0) | 0.100 | [M-H]<sup>-</sup> | 1821.9491 | 2.53 |

### Human Tissue Samples

The project was conducted under the approval of the University of Wollongong Human Ethics Committee protocol number 2022/367. Human midbrain and prefrontal cortex sections were provided by the Queen Square Brain Bank for Neurological Disorders, University College London, UK and shipped frozen under dry ice from the UK to the University of Wollongong. The neuropathological details of clinical donors for the human brain tissues are provided in *Supplementary Information Table 1*. Sections samples had a thickness of 10 micron (µm) and were mounted on microscope glass slides. Tissue sections were kept in −80 °C freezer until analyses. Slides were removed from storage and quickly transferred into hygroscopic desiccator box and vacuum dried for 30 minutes.

### Mouse Brain Samples

Brain specimens were collected from eight-month-old wild type C57BL/6N (Jackson Laboratory, ME, USA) and flash frozen as previously described. Brain tissues were shipped by Merck $ Co., Inc, Rahway, NJ, USA to the University of Wollongong, Australia (UOW) in dry ice (−80°C). Frozen brain specimens were sectioned on the sagittal plane at a depth of 12 µm inside a cryostat (CM1950, Leica Biosystems, Germany) maintained at −20°C and mounted on normal plain microscope glass slides (Rowe Scientific Pty Ltd, Australia). Animal studies were conducted under the approval of the Merck & Co., Inc., Rahway, NJ, USA Institutional Animal Care and Use Committee and endorsed by the Animal Ethics Committee at UOW.

### Mass Spectrometry Sample preparation

Samples were first coated with internal standard lipid mixture of ganglioside. The ganglioside IS (GIS) mixture used for spraying was systematically optimized so that IS signals reflected the intensities observed for endogenous species of the same class. Individual stock solutions in methanol of 7.2 µL C18:0 GM3-d5 (synthetic), 48 µL C18:0 GM1-d5 (synthetic) and 72 µL C17:0 GD1a (d18:1/17:0) standards, total volume 127.2 µL, of were diluted to a total volume of 2,000 µL in methanol. The GIS mixture was deposited following a previous method ^47^. An off-line 2.5 mL Leur lock gas-tight syringe was connected to the TM-sprayer (HTX, Technologies, USA) for application of the ganglioside internal standard (GIS) mixture. Briefly, the sprayer parameters were temperature, 50 °C; number of layers, 16; flow rate, 60 µL/min; velocity, 1,200 mm/min; track spacing, 2 mm; nebulizer gas flow rate, 10 psi, 2L/min and drying time in between passes, 30 sec. The amount of internal standard deposited per unit area was calculated using the following formula:

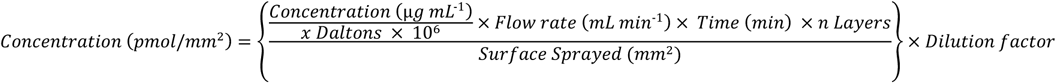

After the GIS was applied the microscope slide sample was briefly returned to the desiccator. Brain tissue sections were then coated with 2, 5-DHA matrix using a TM-Sprayer (HTX Technologies, USA). Matrix application protocol was as follows; 10 mg/ml of 2, 5-DHA in 70% ethanol (aqueous) solution with 50 mM ammonium sulphate, velocity 1200 mm/min, 2 mm track line spacing, 20 layers at a flow rate of 30 µL/min at 80°C and a drying time of 30 sec in between passes (layers).

### Mass Spectrometry Imaging

MALDI-MSI data was acquired using an Atmospheric Pressure (AP)-MALDI UHR source (MassTech, Maryland, USA) coupled to an Orbitrap Fusion Tribrid mass spectrometer (Thermo Fisher Scientific, San Jose, CA, USA). The AP-MALDI source was equipped with a 355 nm (λ) Nd:YAG laser operating at a laser frequency of 500 Hz and attenuated at 7% laser power. The pixel size was set at a lateral distance of 100 µm (x, y). Data was acquired in negative mode at the mass-to-charge ratio (*m/z*) scan range of 500 – 2,500 and mass resolving power of 240,000 FWHM @*m/z* 200 and ion injection time of 500 ms giving a total elapsed scan time of 1.055 sec. The corresponding AP-MALDI MS imaging settings based on these scan settings were an MS speed of 1.37 spectra/sec (i.e., 1.3 spectra per pixel, chosen due to the raster mode acquisition and to ensure each pixel comprises at least one full scan event) using a stage velocity of 0.07291 mm/sec.

### Luxol Fast Blue Staining

Histological staining of neuronal tissue structures was adopted from the modified Klüver-Barrera protocol ^68^. After AP-MALDI-MSI, human and mouse tissue sections were rinsed with methanol for 30 sec to remove matrix followed by washing in 95% ethanol (aq.) and 70% ethanol (aq.) and re-dehydrated in Milli-Q H_2_O (water) for 2 min each. Tissue sections were fixed in 10% neutral buffered formalin for 10 min and air-dried for at least 30 min. Samples were then placed 0.3% acid-alcohol solution for 3 min and then rinsed by immersion in 95% ethanol. Samples were incubated in 0.1% luxol fast blue (LFB) solution in glass Coplin jars heated at 58°C inside an oven overnight. Excess LFB was then rinsed off with 95% ethanol for 1 min and water. Samples were then carefully differentiated in 0.05% lithium carbonate solution for 20 sec followed by two changes of 70% ethanol for up to 20 sec until gray matter was pale while the white matter retained a blue hue. Samples were then rinsed twice with water and remained there after differentiation. Tissue sections were subsequently counter stained by the Nissl method. Samples were placed in warmed freshly-filtered 0.1% cresyl violet solution and placed in an oven set at 50°C for 20 min. Samples were quickly rinsed in water and differentiated in 95% ethanol for 2 – 5 min (and checked microscopically for best result). Samples were then dehydrated by two changes of ethanol for 2 min each and cleared in two changes of xylene. Tissue sections were then cover slipped with DPex mounting medium. Bright-field optical scanned images of LFB-stained samples were digitally captured by a 10X objective lens using a Confocal SP8 FALCON microscope (Leica Microsystems, Germany).

### Mass Spectrometry Imaging Data Analysis

Mass recalibration of MSI data (Thermo .raw) was performed using Recal Offline where necessary using theoretical ion peaks of the three mix ganglioside IS mixture (Table 1) and [PI 38:4-H]^−^ @ *m/z* 885.54985, [SHexCer 42:2;O2-H]^−^ @ *m/z* 888.624007, GM1 36:1;O2-H]^−^ @ *m/z* 1544.86938 and GD1 36:1;O2-H]^−^ @ *m/z* 1863.99609. ProteoWizard MSConvert GUI was used to process .raw files into mzML output ^69^, and then processed into .imzML format using the corresponding .xml position files in LipostarMSI (Molecualr Horizon Srl, Perugia, Italy) ^70^. MSI data (.imzML) for n=3 for either human midbrain, human frontal cortex or mouse tissue samples were imported into LipostarMSI for initial data assessment. Peak detection parameters was perform using Savitzky-Golay smoothing; window size 7 points, 2 degrees and at 1 iterations. Peak picking was set to 0.00 SNR, 0.00 amu noise window size, 0.00 minimum absolute intensity and peaks under 0.50% of the base peak were discarded. Isotopic clustering was performed with an abundance deviation at 30% and *m/z* image correlation threshold at 0.50%.

### MSI and histology data processing

MSI and accompanying LFB-stained histology data were imported into the Weave™ platform (Aspect Analytics NV, Genk, Belgium). The MSI data (.imzML) was imported and converted to an internal, cloud-native format (.zarr). Intensities of the species of interest (target ganglioside list and internal standards) were extracted through binning of the spectra around the corresponding m/z value. A window of ± 3 ppm around the target m/z value was applied, and the mean value for each pixel in this window was taken as the ion intensity for said pixel. Images were normalized by conducting a pixel-wise division of the original m/z image by its matching internal standard m/z image. Winsorizing was employed to eliminate hotspots from the normalized images, replacing intensities above the .995 quantile with the value of that quantile. Missing values resulting from zero division in the normalization process were ignored for downstream analysis. After normalization to the reference image, the normalized pixel intensities are multiplied by their matching internal standard concentration to achieve the final concentration images.

MSI and the LFB histological images were co-registered using a landmark-based, non-rigid registration algorithm in the Weave platform, mapping both datasets to a shared coordinate system. Anatomical regions of interest (ROIs) were acquired in two ways, (i) through direct annotation of the LFB histology mouse and human tissues and (ii) through integration with the Allen Mouse Brain Atlas (AMBA) for only the mouse tissues. For (i), pathologists were given access to the portal to perform annotation of the LFB-stained brain tissues, where they marked gray matter and white matter regions in either mouse cortex, mouse midbrain sagittal sections, (human) frontal cortex or (human) midbrain tissues. Through the shared coordinate system, these ROIs were transferred to the MSI data.

For (ii), the closest matching Nissl histology section in the AMBA (sagittal section 113) was downloaded alongside its anatomical annotations through the AMBA API. The AMBA histology was imported into the Weave platform, and co-registered with the LFB histology using the same landmark-based, non-rigid registration approach described above. In this way, LFB microscopy, MSI, the AMBA histology and its accompanying atlas annotations were all mapped to the same coordinate system, similar to previous work ^71^. This allows the atlas annotations to be transferred to the MSI data, and used in the same manner as the manually annotated pathologist ROIs.

The AMBA provides highly detailed anatomical annotations of the mouse brain, with very fine spatial regions. Given that the MSI data is acquired in a different animal, and that the tissue is deformed in extraction, freezing and sectioning, even detailed non-rigid registration cannot guarantee proper alignment of small brain structures. For this reason, we only use sufficiently large anatomical regions in the AMBA, resulting in 12 AMBA-defined ROIs; cerebellar cortex, cerebellar nuclei, fiber tracts, hindbrain, hippocampal region, interbrain, isocortex, midbrain, olfactory areas, pallidum, retrohippocampal region and striatum. The co-registered ROIs were used to extract ion intensities from the pixels within these ROIs. The mean concentrations for each anatomical region were used for downstream analysis.

### Ganglioside Species Identification List

Lipid species belonging to the ganglioside class were identified using LipostarMSI and manually curated using a merged MSI data set of mouse and human brain tissues. The combined feature list was subjected to an *m/z* tolerance of 0.000 amu ± 3.0 ppm and searched against the Lipid Maps “bulk” Structure Database (LMSD) for deprotonated ions [M-H]^−^. The list was then approved for even chain species. The resulting ID compounds were then exported as .csv files. Additionally, we used the GlyCosmos database and other peer-reviewed literature sources ^28, 34^ to compile a list of known brain gangliosides and other possible theoretical matches not identified through LMSD. Here, all mass (*m/z*) features in the LipostarMSI super sample lipid compound list were cross-referenced to the expanded GlyCosmos list to detect other present ganglioside masses. Ganglioside species without a direct IS were paired with the most closely glycan assembly on the LacCer backbone. From these potential matches, a spatial filtering was performed, followed by a manual check of image quality to remove off-tissue signals that happened to match to possible gangliosides and ensure each ion image showed district structure throughout all replicate of at least one sample type. Spatial filtering was performed per peak. For each tissue, two coherence metrics were computed on the annotated-region ion image and normalized by the corresponding full-tissue value: (i) a neighborhood intensity score, taking the mean intensity of the 8-connected neighbors of each non-zero pixel, and (ii) connectivity, the mean fraction of those neighbors that were also non-zero. A peak was retained if its normalized neighborhood intensity score or normalized connectivity reached ≥ 0.9 in at least 5 samples. The final manually assembled list included an IS normalization reference peak for each target endogenous *m/z* species.

### Correction In-source Fragmentation

Loss of sialic acid (Neu5Ac) residues is commonly observed during MALDI analysis of gangliosides with the resulting fragments being identical to true endogenous lipids. For example, GD1 will lose a Neu5Ac ([M-H-H_2_O]^−^ = *m/z* 290.0881; *M =* 291.0954 Da) and form the corresponding GM1. As such, the GM1 abundance is over-represented and the GD abundance under-represented. To correct for this, we used the GD1 IS to determine the extent of fragmentation across a given tissue and thus determine how much of the GM1 signal was artificially enhanced. To do this we first calculated the fragmentation ratio (i.e signal of the GM1 fragment, *m/z* 1530.8537 formed from the GD1 IS, *m/z* 1821.9491 as a fraction of the intact GD1 IS). From this ratio the additional signal added to a given endogenous GM1x:y:z produced from the corresponding GD1x:y:z (corrections are applied to same x:y:z pair, e.g. GD136:1;O2 and GM1 36:1;O2). This additional signal could then be calculated in pmol/mm^2^ following isotope correction and this concentration removed from the endogenous GM1x:y:z ion and added to the GD1x:y:z. The fragmentation corrected concentrations were then aggregated per ROI.

### Tandem Mass Spectrometry Ganglioside Validation

Tandem mass spectrometry (MS/MS) was carried out for select main ganglioside species. On the Orbitrap Fusion, to enhance product ion signals, the laser repetition, lateral resolution and stage velocity were adjusted 2,000 Hz, 0.01 mm and 0.12 mm/sec, respectively. At MS2 level, FTMS scans were conducted with an injection time of 2,500 msec for each precursor for collision induced dissociation (CID) and high energy CID (HCD) activation with an isolation window, 2.0 Da, normalized AGC target, 250%. The MS2 activation time was 10 msec with normalized (%) CID energy at 33.5 or HCD at 34.0. On the timsTOF Flex MALDI-2 (Bruker Daltonics, GmBH, Bremen, Germany), the instrument was calibrated in negative polarity using the Agilent Tune Mix in enhanced quadratic method prior to MALDI-MS/MS on-tissue data acquisition for odd-chain ganglioside species. The laser parameters were 5,000 shots at 10,000 Hz, 60% laser power with a beam scan size of 150 μm (x, y). The collision energy was set at 80 V.

### Identification of gangliosides by LC-MS

Prior to analyses, rodent brains were homogenized using HPLC water at 4 ℃ at around 0.1 g/ml. 50 mL aliquots were transferred to 96-well plates mixed with 10 µg/ml of d_3_GM1C_18_, d_9_GM2C_16_, d_3_GM3C^18^, d_7_GD1bC_18_, d_7_GD2C_18_, d_7_GD3C_18_, d_7_GT3C_18_ each (AVANTI Polar Lipids (Alabaster, Alabama, USA). 1 ml of CHCl_3_, 500 ml of MeOH, 250 ml of 0.015 M NaCL were added and the samples were vortexed for 10 minutes, followed by centrifugation for 10 min at 3300 RCF. The upper phase was collected and 200 ml of 0.015M NaCl added to the lower phase, followed by mixing for 10 minutes and centrifugation for 10 min at 3300RCF. The upper phase was collected and combined with the first upper phase.

The gangliosides were further purified by SPE (INERTSEP 96WP PHARMA, 60MG, NC3455612). 1 mL of MeOH/0.015M NaCl (1/1) was used for conditioning. The combined upper phases were loaded on to the SPE and by flow through gravity collected. The collected flow through was reloaded on to the SPE plate. The SPE was washed using 1 mL water. 750 mL of MeOH was added, followed with 750 mL of MeOH/CHCl_3_ (1/1), and the flow through was collected, dried down, and reconstituted 100 mL MeOH prior to LC-MS analysis.

Selected gangliosides species were monitored using multiple reaction monitoring scanning acquisition using a SCIEX 4500B QTRAP mass spectrometer operating in negative electrospray ionization mode (Sciex LLC, Framingham, MA) coupled to a Waters Acquity UPLC instrument (Waters Corp., Milford, MA). Separation was achieved using a Waters XBridge BEH Phenyl 2.1×50 mm, 3.5 μm, 130Å pore size, (Cat# 186003322) using a thirty-minute gradient running from low to high organic mobile phase buffer (mobile phase A: 0.028% (v/v%) ammonium hydroxide in water; mobile phase B: 0.028% (v/v%) ammonium hydroxide in MeOH according to Table 1. Internal standards were used as guidance for the identification of the endogenous species. The verified ganglioside species are shown in *Supplementary File 1*.

**Table 2.** LC gradient used for LC-MS/MS verification of gangliosides.

| Time<br>(min) | Flow Rate (mL/min) | A% | B% |
| --- | --- | --- | --- |
| 0 | 0.6 | 95 | 5 |
| 5 | 0.6 | 95 | 5 |
| 6 | 0.3 | 75 | 25 |
| 24 | 0.3 | 0 | 100 |
| 25 | 0.3 | 0 | 100 |
| 28 | 0.6 | 95 | 5 |
| 30 | 0.6 | 95 | 5 |

