## Supplementary Information for "Quantitative Imaging and Characterization of Ganglioside Diversity in Mouse and Human Brain Tissues"

**Table of Contents**

Supporting Tables ... 3

Supplementary Table 1. Neuropathological details of clinical donors for human brain tissues ... 3

Supplementary Table 2. *m/z* values used for mass recalibration (all detected as $[M-H]^-$ ions). ... 4

Supporting Figures ... 5

Supplementary Information Figure 1: Corrected GM1 concentrations. ... 5

Supplementary Information Figure 2: Human and Mouse White Matter and Gray Matter Comparisons ... 6

Supplementary Information Figure 3A. MS/MS spectra of selected gangliosides detected in mouse brain. ... 7

Supplementary Information Figure 3B. Identification of acyl chain length in gangliosides ... 8

Supplementary Information Figure 4A. MS/MS spectra of selected gangliosides detected in human frontal cortex. ... 9

Supplementary Information Figure 4B. Identification of acyl chain length in gangliosides. ... 10

Supplementary Information Figure 5. MS/MS spectrum of selected odd-chain ganglioside, supporting their identification. ... 11

Supplementary Figures 6. Images of manually curated list of identified gangliosides *m/z* ion composite brain tissue images. ... 12

**Supporting Tables**

**Supplementary Table 1. Neuropathological details of clinical donors for human brain tissues**

| *Patient ID* | *Gender* | *Disease* | *Age at Death* | *Sample area* | *Mutation status* | *Genetics* | *A-Syn load* | *Braak Tau* | *Braak LB/LB score* | *LB disease type* | *Thal* | *CERAD* | *ABC* | *Other Path* |
|---|---|---|---|---|---|---|---|---|---|---|---|---|---|---|
| *26* | F | Control | 79 | Frontal cortex and Midbrain | Unknown | none stated | no a syn pathology seen | I | 0 | none | not stated | sparse | not stated | Pathological ageing , Mild hyaline arteriolosclerosis |
| *34* | F | Control | 92 | Frontal cortex and Midbrain | Unknown | none stated | no a syn pathology seen | III | 0 | none | 0 | sparse | not stated | Pathological ageing |
| *42* | M | Control | 71 | Frontal cortex and Midbrain | Unknown | none stated | no a syn pathology seen | 0 | 0 | none | 0 | 0 | A0B0C0 | No significant abnormalities |

*a syn – alpha (α)-synuclein*
*A – Amyloid (Thal Phase); score for amyloid-β plaques*
*B – Braak stage (Tau); assessment for spread of neurofibrillary tangles*
*C – CERAD score for neuritic plaques*
*LB – Lewy body*
*ABC – Combination of scores for standardized assessment / classification of Alzheimer's Disease neuropathological changes*
*CERAD – Consortium to Establish a Registry for Alzheimer's Disease*

**Supplementary Table 2.** ***m/z*** **values used for mass recalibration (all detected as [M-H]$^-$ ions).**

| | Lipid / Internal Standard | Molecular Formula | *m/z* |
|---|---|---|---|
| **1.** | C18:0 GM1-d5 (synthetic) | $C_{73}H_{126}D_5N_3O_{31}$ | 1549.90076 |
| **2.** | C17:0 GD1a (d18:1/17:0) | $C_{83}H_{146}N_4O_{39}$ | 1821.94914 |
| **3.** | C18:0 GM3-d5 (synthetic) | $C_{59}H_{103}D_5N_2O_{21}$ | 1184.76856 |
| **4.** | SHexCer 42;2;O2 | $C_{48}H_{91}NO_{11}S$ | 888.624007 |
| **5.** | GM1 36:1;O2 | $C_{73}H_{131}N_3O_{31}$ | 1544.86938 |
| **6.** | GD1 38:1;O2 | $C_{86}H_{152}N_4O_{39}$ | 1863.99609 |
| **7.** | PI 38:4 | $C_{47}H_{83}O_{13}P$ | 885.54985 |

**Supporting Figures**

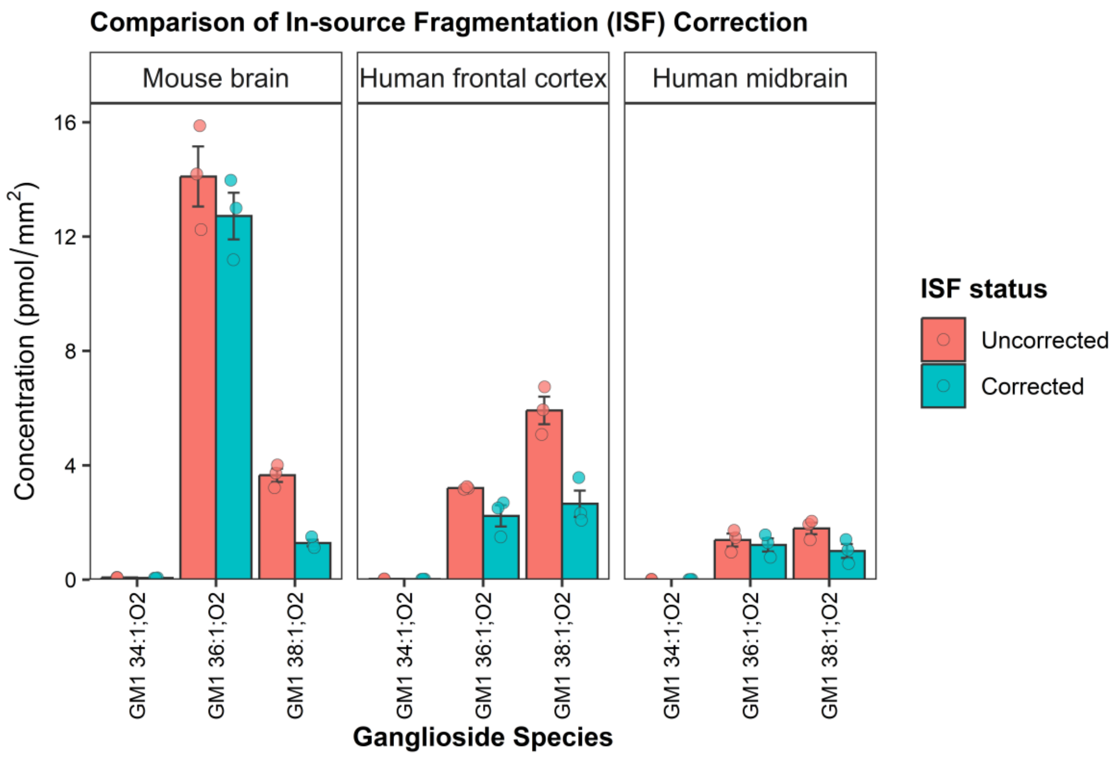


**Supplementary Information Figure 1: Corrected GM1 concentrations.**

Comparison of GM1 concentrations obtained by Q-MSI before (red bars) and after (blue bars) correction of GD1 in source fragmentation (ISF). Each bar shows the Q-MSI derived concentration for a particular GM1 lipid across mouse (left side), human frontal cortex (middle) and human midbrain (right) whole tissues, with each biological replicate represented by a circle. Error bars represents standard error of means. ISF correction utilised the extent of C17:0 GD1a (d18:1/17:0) *m/z* 1821.9491 breaking down to its corresponding GM1 analogue at *m/z* 1530.8537. This calculated fragmentation ratio was then used to estimate how much signal of a given GM1 species was the result of the corresponding GD1 species. Further details are provided in the Experimental Methods.

a) Mouse Cortex Fold Change Gray Matter vs White Matter

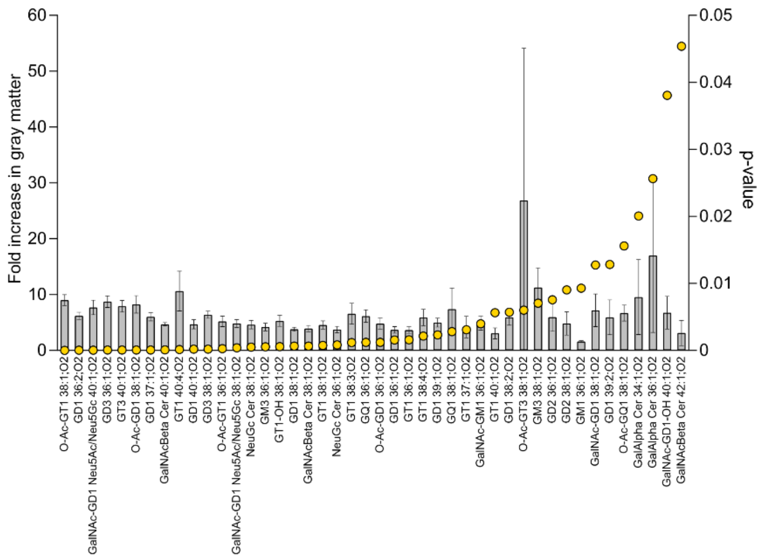


b) Human Cortex Fold Change Gray Matter vs White Matter

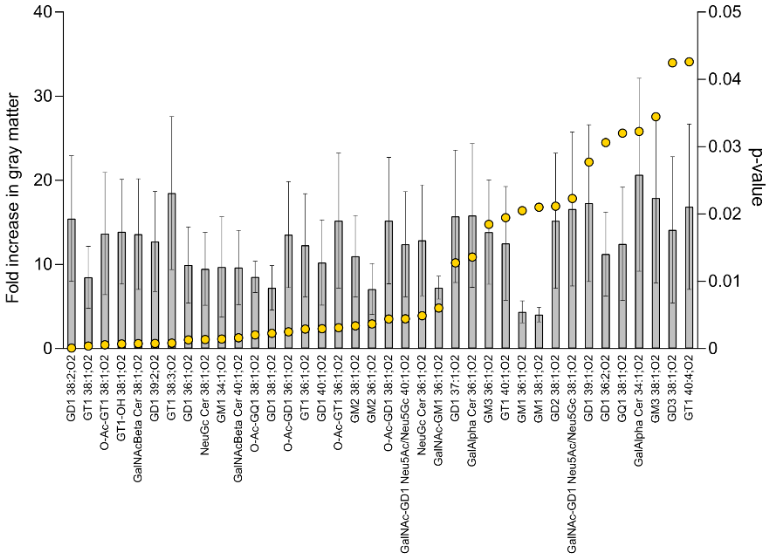


**Supplementary Information Figure 2: Human and Mouse White Matter and Gray Matter Comparisons**

Fold change bar plots of ganglioside concentration showing significant fold change increase in annotated gray matter regions compared to white matter comparisons for (a) mouse cortex and (b) human frontal cortex tissues. Data points depicted as yellow dots represent the significance level indicated by $log_2$ p-value, right y-axis

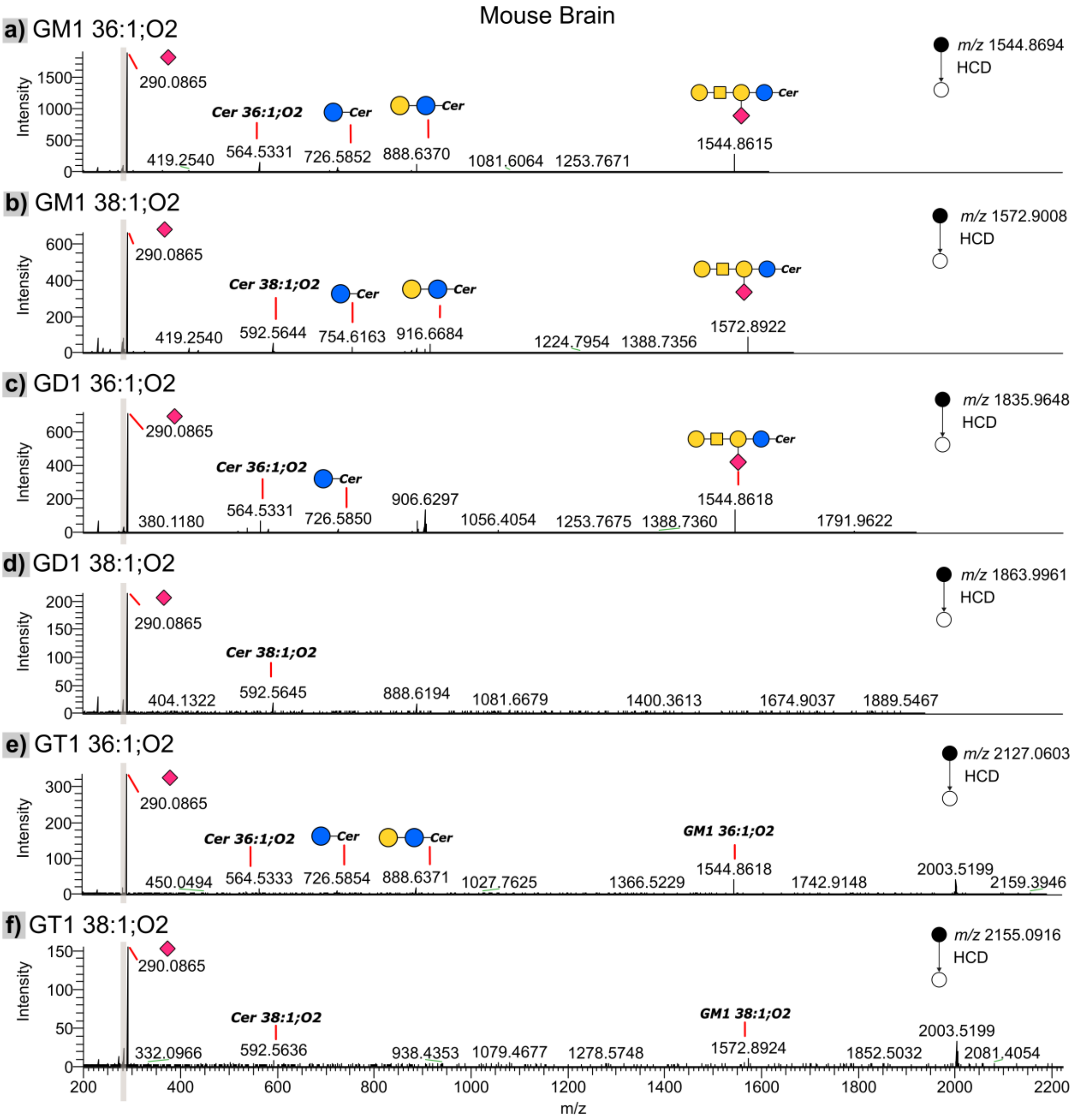


**Supplementary Information Figure 3A. MS/MS spectra of selected gangliosides detected in mouse brain.**

On-tissue MS/MS spectra acquired from $[M-H]^-$ ions of (a) GM1 36:1;O2, (b) GM1 38:1;O2, (c) GD1 36:1;O2, (d) GD1 38:1;O2, (e) GT1 36:1;O2 and (f) GT1 38:1;O2 detected from mouse brain. MS/MS was performed using an injection time of 2500 ms; normalized AGC target of 250%, higher collision dissociation (HCD) with a normalised collision energy of 34% (normalised), mass resolution of 240,000 and the instrument operated in high mass mode with FTMS detection. Characteristic fragment ion peaks are depicted with glycan notation where possible. The Hex2Cer 36:1;O2 *m/z* 888.6370, HexCer 36:1;O2 *m/z* 726.58, Cer 36:1;O2 *m/z* 564.53 and Neu5Ac *m/z* 290.08 (sialic acid-$H_2O$) product ions are observed from MS/MS of (a) GM1 36:1;O2, (c) GD1 36:1;O2 and (e) GT1 36:1;O2. The Hex2Cer 38:1;O2 *m/z* 916.66, HexCer 38:1;O2 *m/z* 754.61, Cer 38:1;O2 *m/z* 592.56 and Neu5Ac *m/z* 290.08 (sialic acid-$H_2O$) product ions are observed from MS/MS of (b) GM1 38:1;O2, (d) GD1 38:1;O2 and (f) GT1 38:1;O2. The gray box indicates low mass range ~ *m/z* 274 – 284.

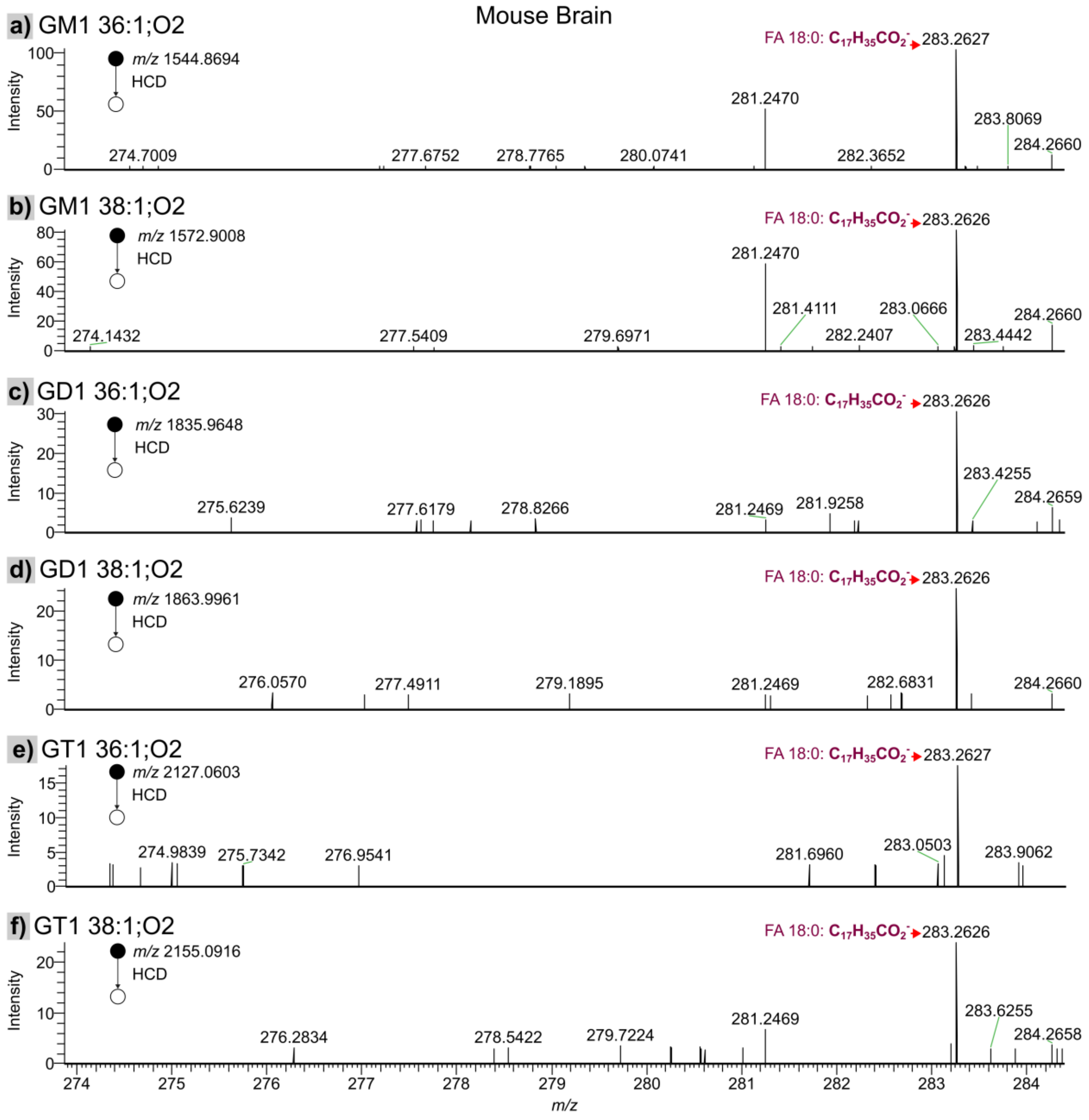


**Supplementary Information Figure 3B. Identification of acyl chain length in gangliosides**

Zoomed in mass range from *m/z* 274-284 for the MS/MS spectra shown in *Supplementary Information Figure 2A* revealing diagnostic fragments allowing the assignment of acyl chain length. Data is acquired from the $[M-H]^-$ ions of (a) GM1 36:1;O2, (b) GM1 38:1;O2, (c) GD1 36:1;O2, (d) GD1 38:1;O2, (e) GT1 36:1;O2 and (f) GT1 38:1;O2 detected from mouse brain. The diagnostic fragment ion at *m/z* 283.26 allows assignment of the fatty acid as an 18:0 chain and thus long chain base (LCB) as either d18:1 or d20:1 for 36:1;O2 and 38:1;O2 sum composition species, respectively. The fragmentation mechanism is outlined in detail in reference according to elaborate CID characterization studies of ceramide standards by Hsu and Turk 2002 (reference 54 in main text). Briefly, the low abundant *N*-acylsphingosine, (N-(octadecanoyl)-sphing-4-enine at *m/z* 564 or *N*-octadecanoyl)-eicosasphing-4-enine at *m/z* 592) is formed from a proposed fragmentation cascade (data not shown, full scheme found in reference 54). Initial C-C bond cleavage at position 2-3 of the sphingoid chain to produce an *N*-stearoylaminoethanol anion by loss of the LCB as an aldehyde. Further loss of $H_2$ from this common *N*-acyl-aminoethanol produces an *N*-stearoylaminoethylen-1-ol which may rearrange to a carboxyethenolamine anion that results into a stearic carboxylate ion, $C_{17}H_{35}CO_2^-$, *m/z* 283.26, by elimination of an arizine group.

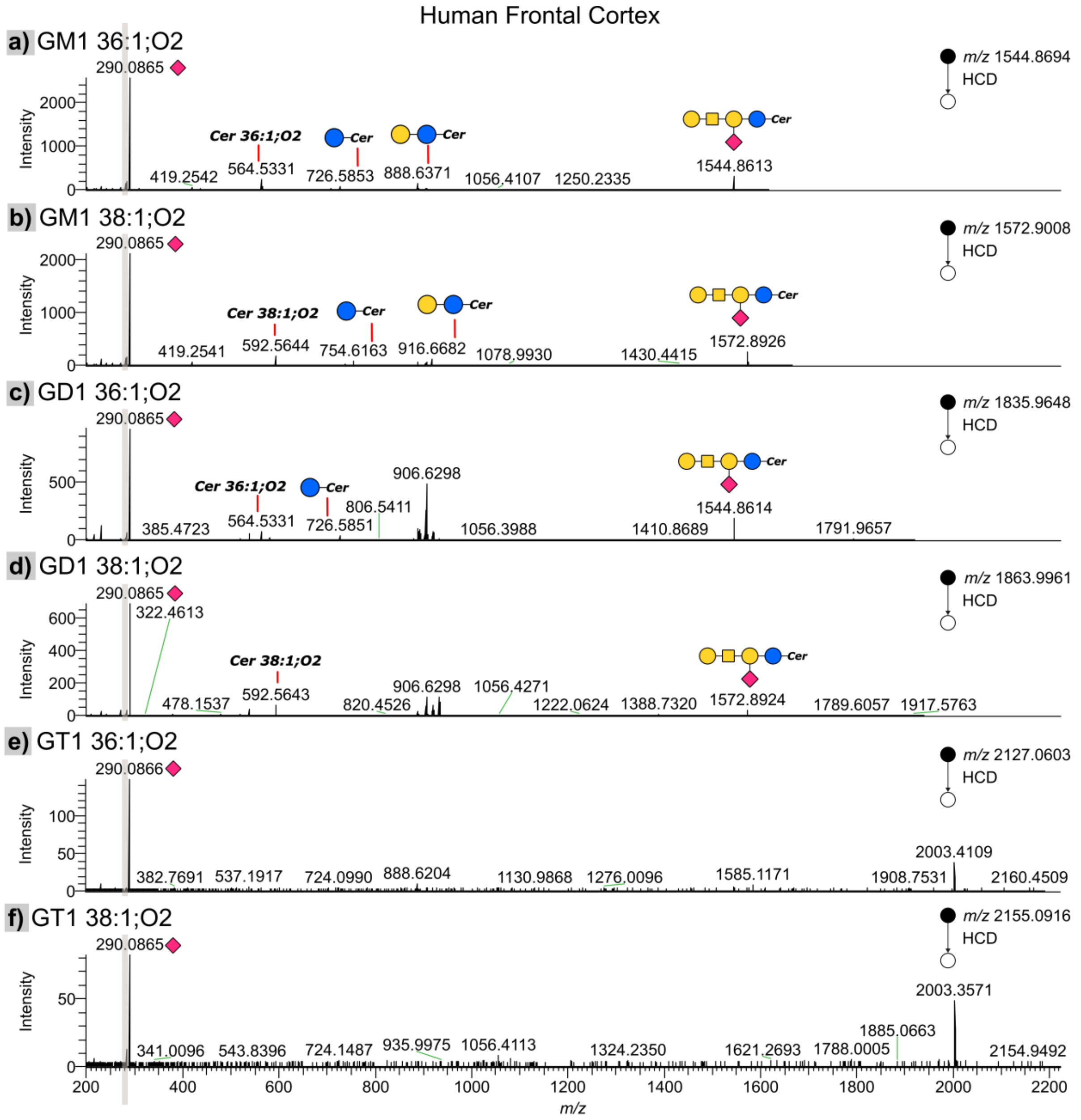


**Supplementary Information Figure 4A. MS/MS spectra of selected gangliosides detected in human frontal cortex.**

On-tissue MS/MS spectra acquired from $[M-H]^-$ ions of (a) GM1 36:1;O2, (b) GM1 38:1;O2, (c) GD1 36:1;O2, (d) GD1 38:1;O2, (e) GT1 36:1;O2 and (f) GT1 38:1;O2 from on-tissue MALDI-MS analysis in human frontal cortex tissue. The settings for MS/MS are as described in *Supplementary Information Figure 2A*. The Hex2Cer 36:1;O2 *m/z* 888.6370, HexCer 36:1;O2 *m/z* 726.58, Cer 36:1;O2 *m/z* 564.53 and Neu5Ac *m/z* 290.08 (sialic acid-$H_2O$) product ions are observed from MS/MS of (a) GM1 36:1;O2, while HexCer 36:1;O2 and Cer 36:1;O2 are observed for (c) GD1 36:1;O2. The Hex2Cer 38:1;O2 *m/z* 916.66 38:1;O2, HexCer 38:1;O2 *m/z* 754.61 and Cer 38:1;O2 *m/z* 592.56 product ions are observed from MS/MS of b) GM1 38:1;O2. The Cer 38:1;O2 *m/z* 592.56 and GM1 38:1;O2 fragments can be seen for (d) GD1 38:1;O2. Both GD1 36:1;O2 and GD1 38:1;O2 yield the corresponding equivalent GM1 fragments through $[M-Neu5Ac]^-$. The less abundant (e) GT1 36:1;O2 and (f) GT1 38:1;O2 predominantly generate sialic acid-related fragments at *m/z* 290.09.

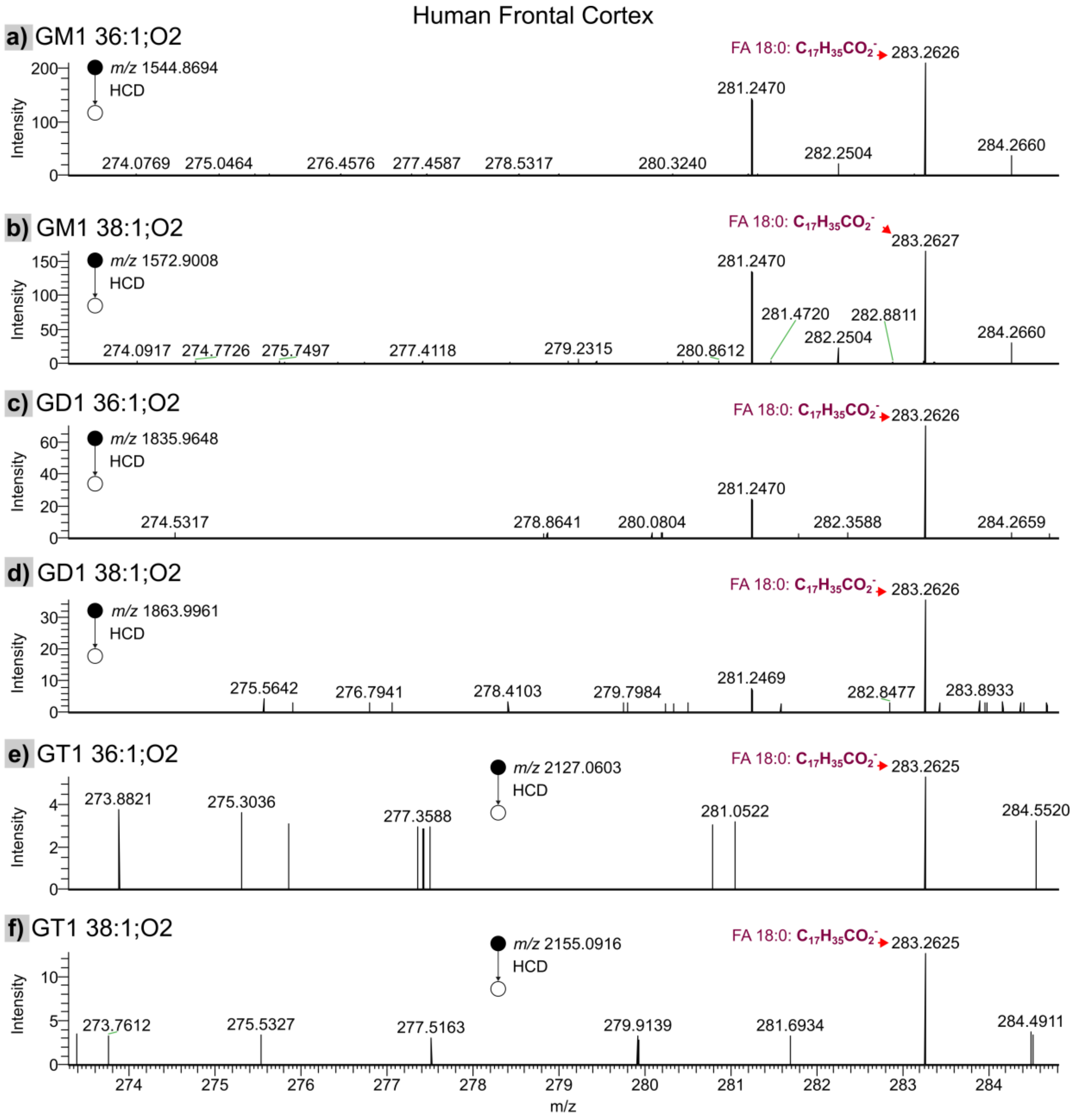


**Supplementary Information Figure 4B. Identification of acyl chain length in gangliosides.**

Zoomed in mass range from *m/z* 274 - 284 for the MS/MS spectra shown in *Supplementary Information Figure 3A* revealing diagnostic fragments allowing the assignment of acyl chain length. Data is acquired from the $[M-H]^-$ ions of (a) GM1 36:1;O2, (b) GM1 38:1;O2, (c) GD1 36:1;O2, (d) GD1 38:1;O2, (e) GT1 36:1;O2 and (f) GT1 38:1;O2. The lower mass range end shows the fatty acyl product ion FA 18:0 across three separate mass channels of the respective ganglioside precursor masses. The assignment of the LCB base as d18:1 for GM1 36:1;O2, GD1 36:1;O2 and GT1 36:1;O2, and as d20:1 for GM1 38:1;O2, GD1 38:1;O2 and GT1 38:1;O2 is as previously rationalized in *Supplementary Information Figure 2B*. That is, the identification of the uniform diagnostic fragment ion at *m/z* 283.26 is consistent with the mouse brain MS/MS data for a common 18:0 fatty acid for both 36:1;O2 and 38:1;O2 species.

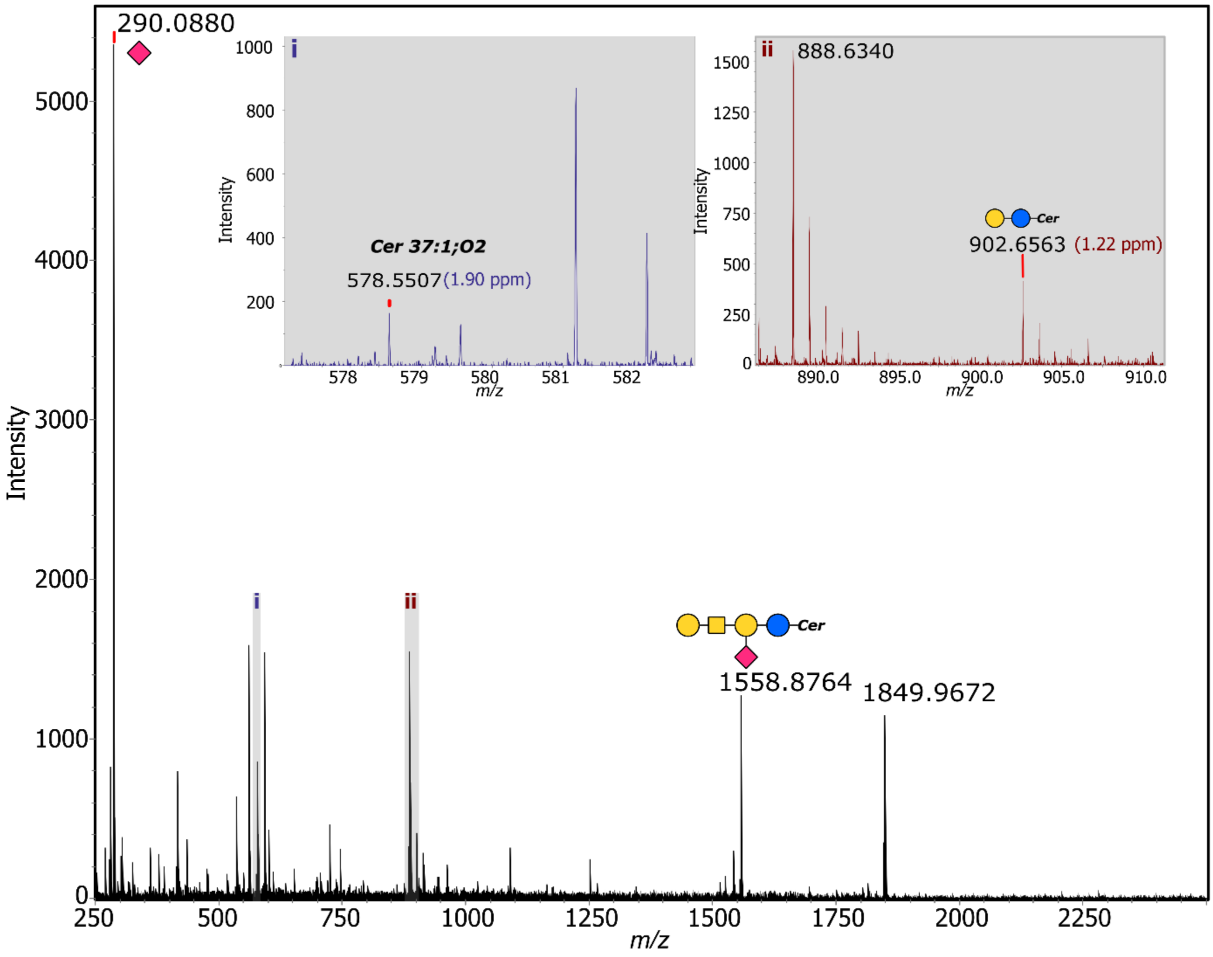


**Supplementary Information Figure 5. MS/MS spectrum of selected odd-chain ganglioside, supporting their identification.**

MS/MS spectrum of *m/z* 1849.96 assigned as the [M-H]$^-$ ion of GD1 37:1;O2 on human frontal cortex tissue. The corresponding GM1 37:1;O2 fragment is observed at the expected *m/z* 1558.87. The highlighted gray areas with inset boxes show confirmatory product ions peaks at corresponding to Cer 37:1;O2 at *m/z* 578.55 and Hex2Cer 37:1;O2 at *m/z* 902.65. The Neu5Ac fragment peak is shown at *m/z* 290.08. Data was acquired using a timsTOF Flex.

**Supplementary Figures 6. Images of manually curated list of identified gangliosides *m/z* ion composite brain tissue images.**

Top row, human frontal cortex (FC); middle row: human midbrain (MB) and bottom row; sagittal mouse brains. The IS normalised Q-MSI images are displayed at 99.9 quantile. Scale bar on the right denotes concentration as pmol/mm$^2$.

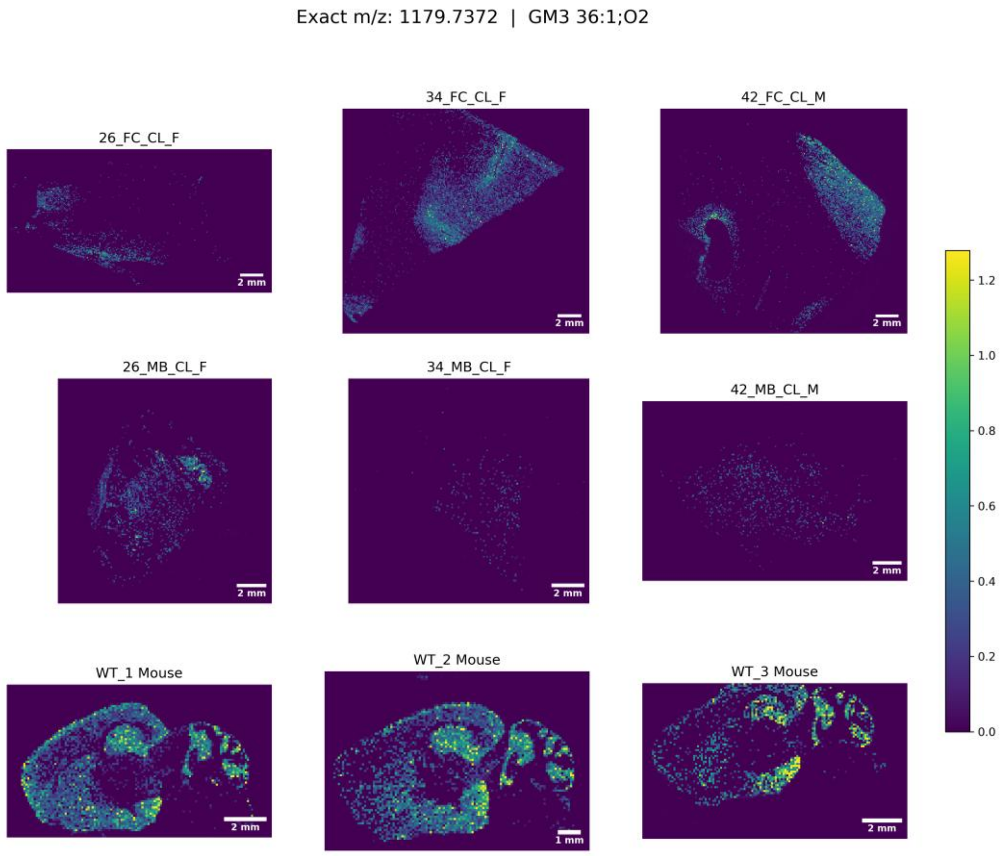

Exact m/z: 1207.7685 | GM3 38:1;O2

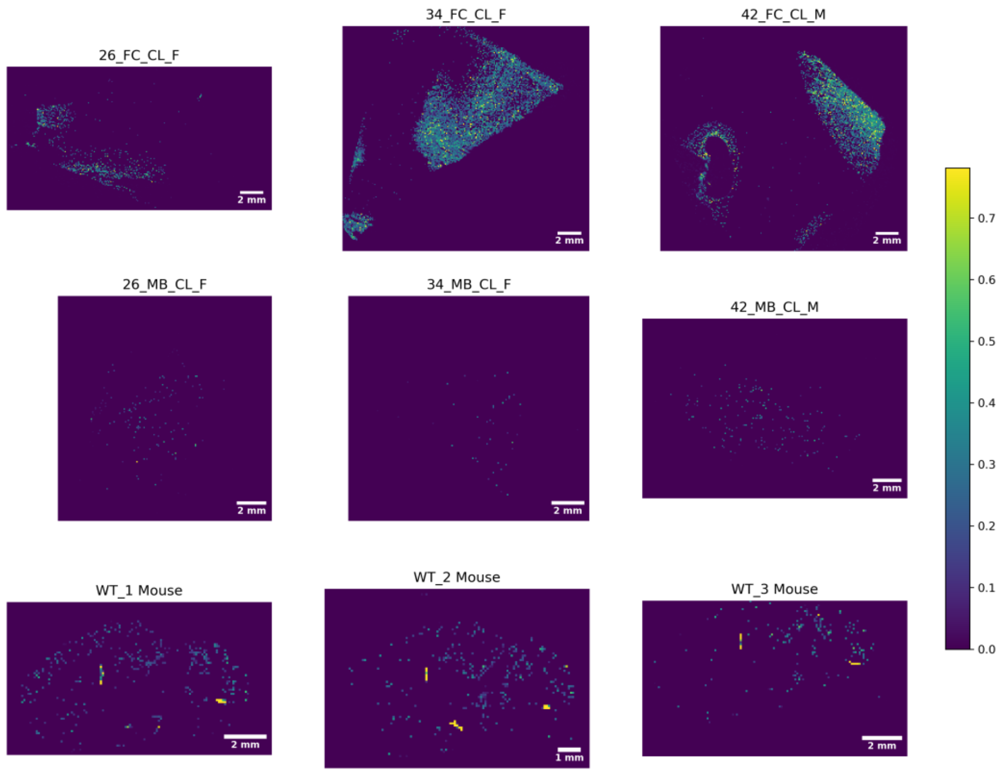


Exact m/z: 1382.8166 | GM2 36:1;O2

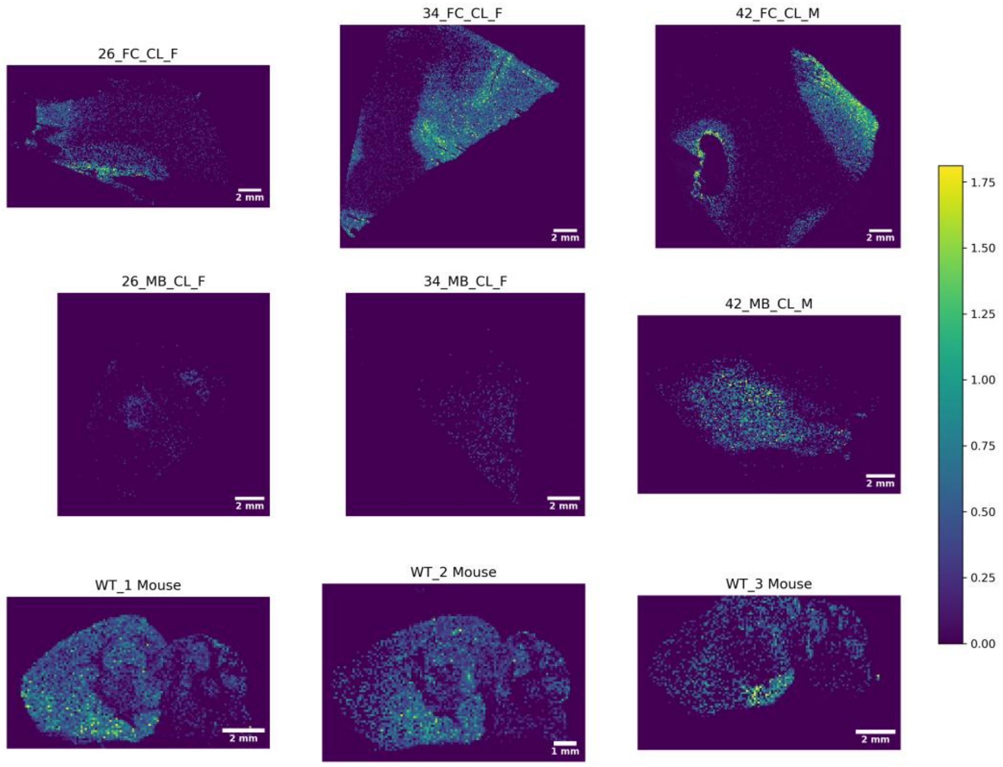

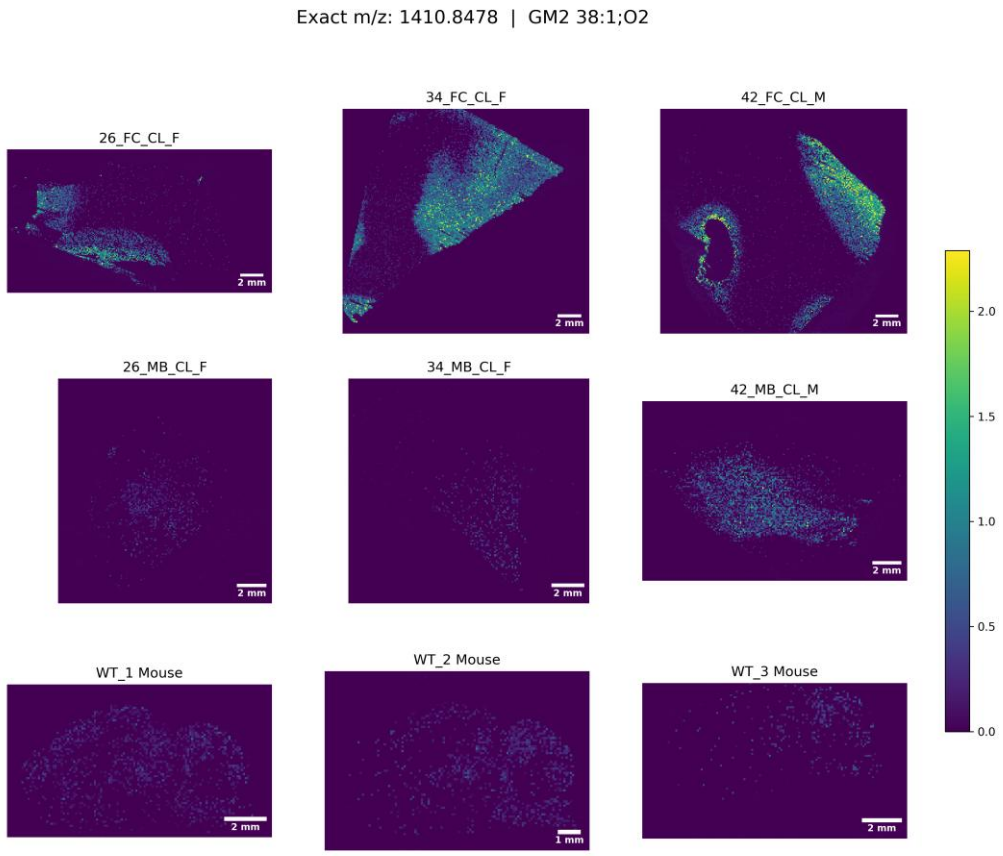
Exact m/z: 1410.8478 | GM2 38:1;O2
26_FC_CL_F
34_FC_CL_F
42_FC_CL_M
26_MB_CL_F
34_MB_CL_F
42_MB_CL_M
WT_1 Mouse
WT_2 Mouse
WT_3 Mouse
2 mm
1 mm
2.0
1.5
1.0
0.5
0.0

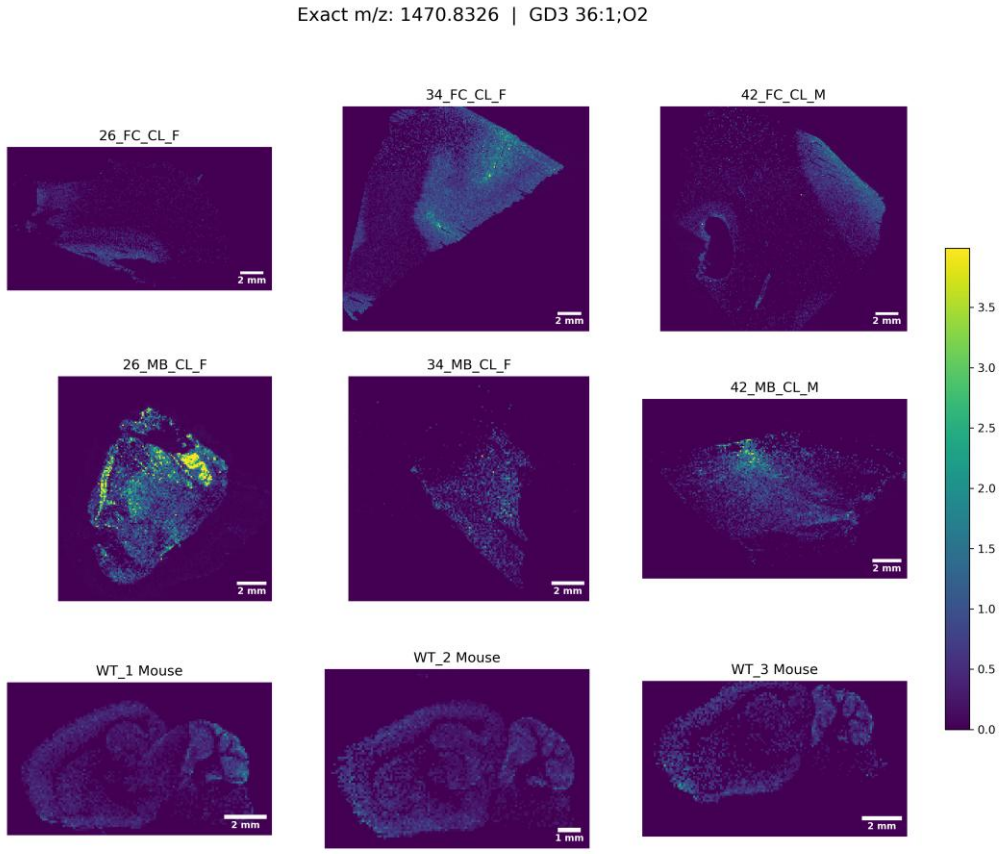
Exact m/z: 1470.8326 | GD3 36:1;O2
26_FC_CL_F
34_FC_CL_F
42_FC_CL_M
26_MB_CL_F
34_MB_CL_F
42_MB_CL_M
WT_1 Mouse
WT_2 Mouse
WT_3 Mouse
2 mm
1 mm
3.5
3.0
2.5
2.0
1.5
1.0
0.5
0.0

Exact m/z: 1498.8639 | GD3 38:1;O2

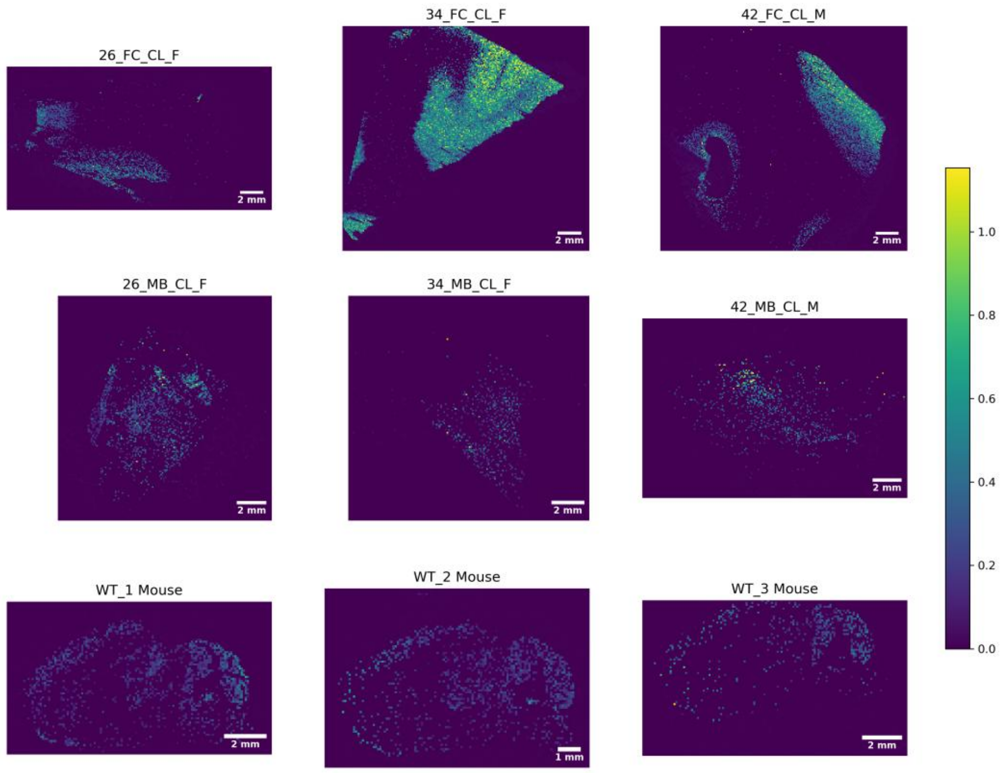


Exact m/z: 1516.8381 | GM1 34:1;O2

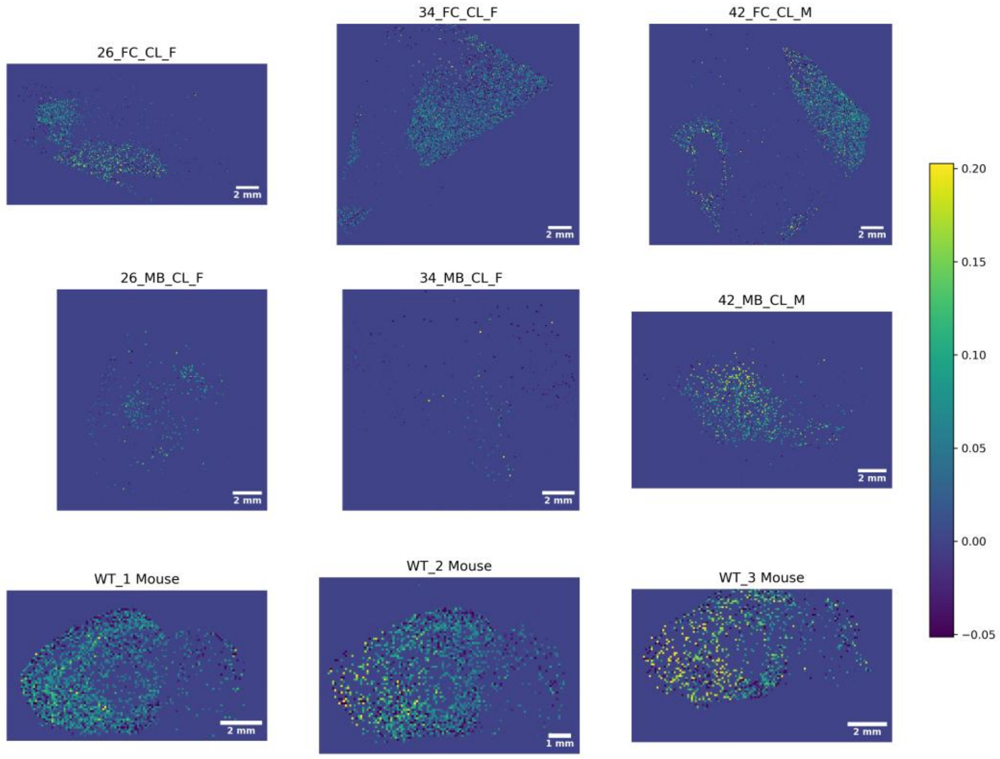

Exact m/z: 1544.8694 | GM1 36:1;O2

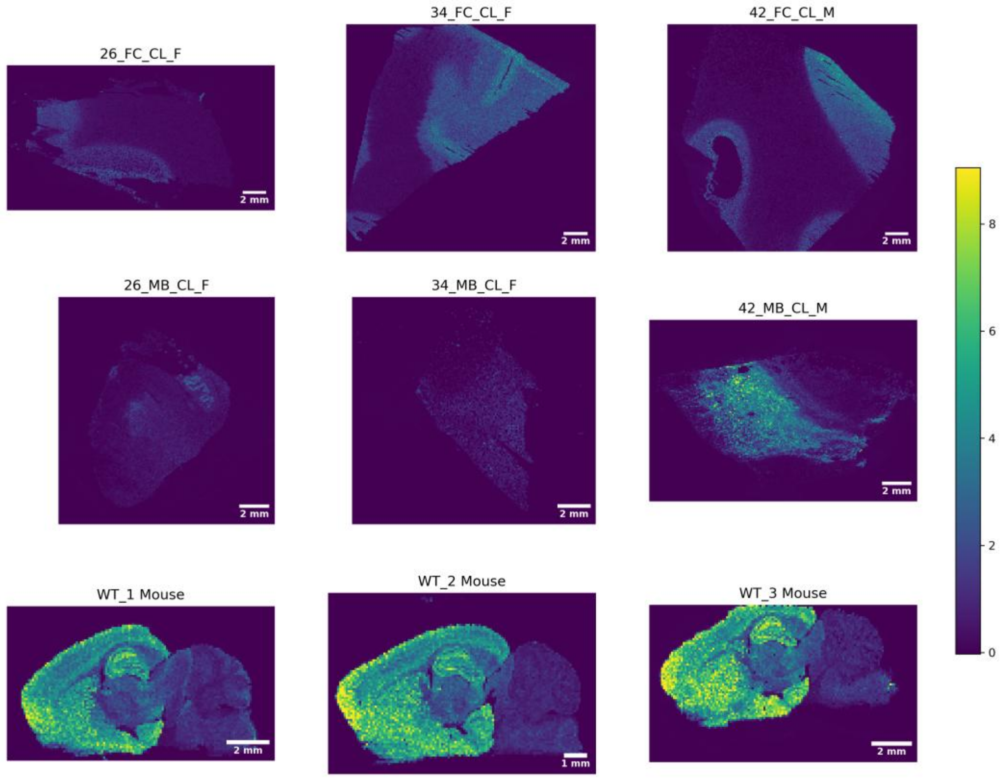


Exact m/z: 1572.9007 | GM1 38:1;O2

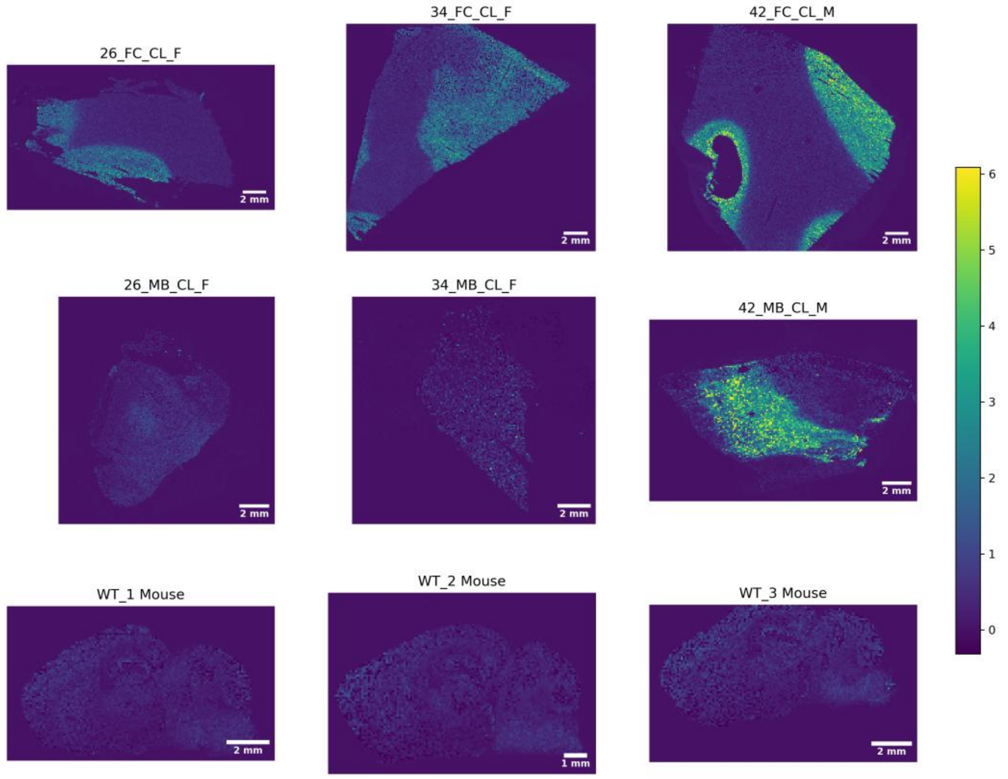

Exact m/z: 1673.912 | GD2 36:1;O2

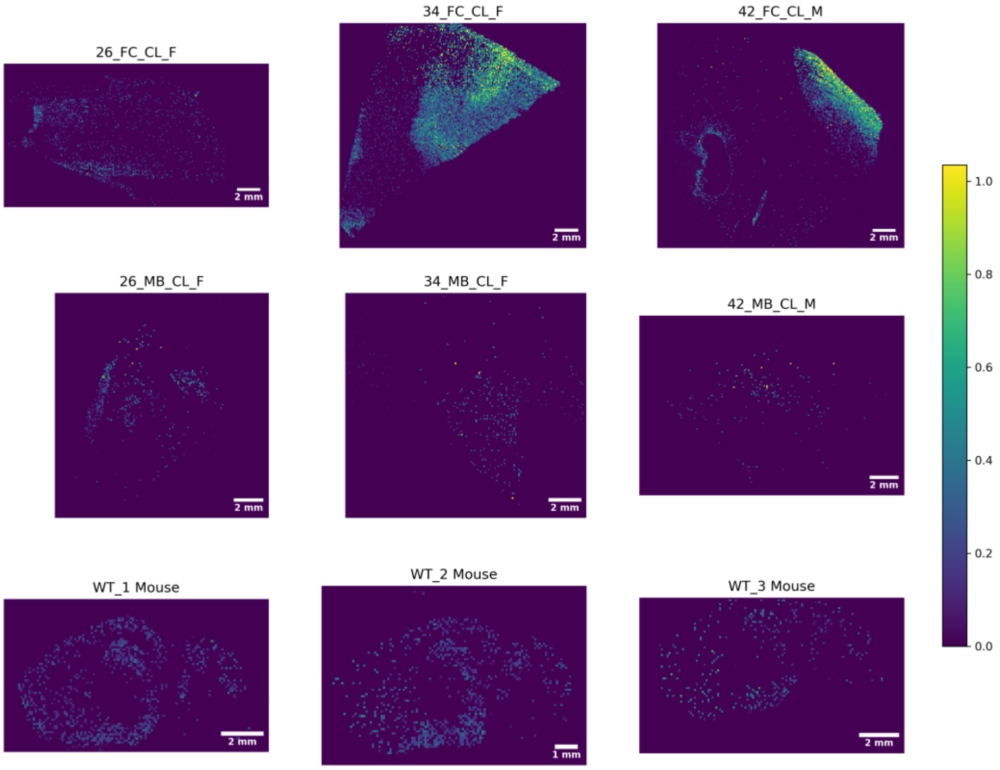


Exact m/z: 1701.9433 | GD2 38:1;O2

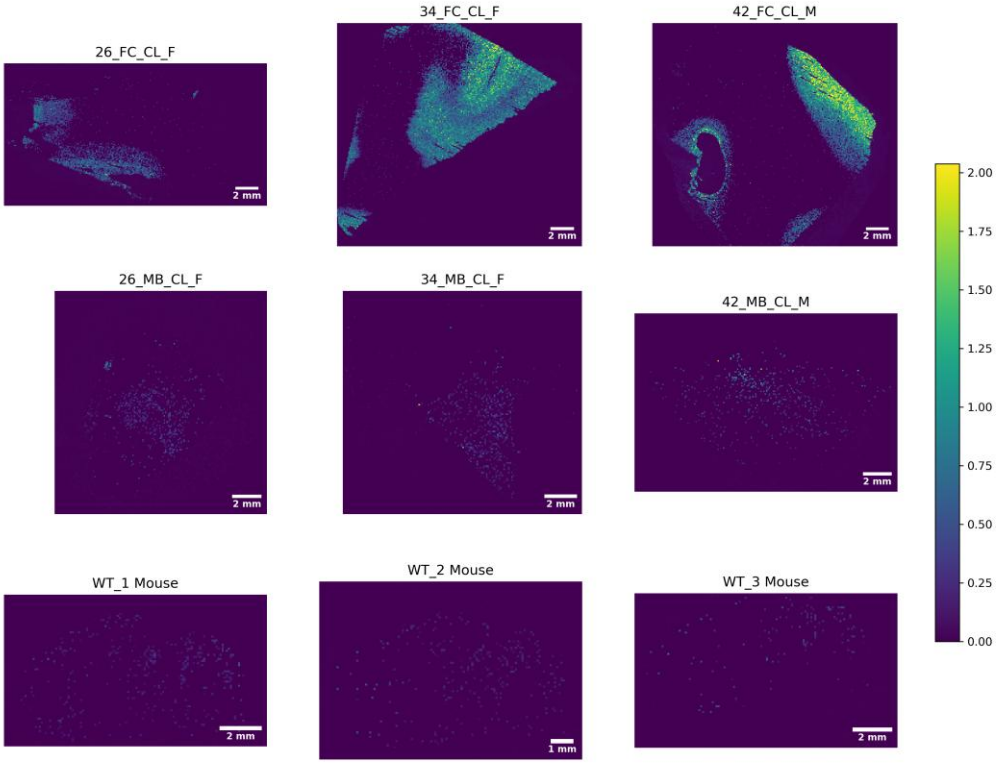

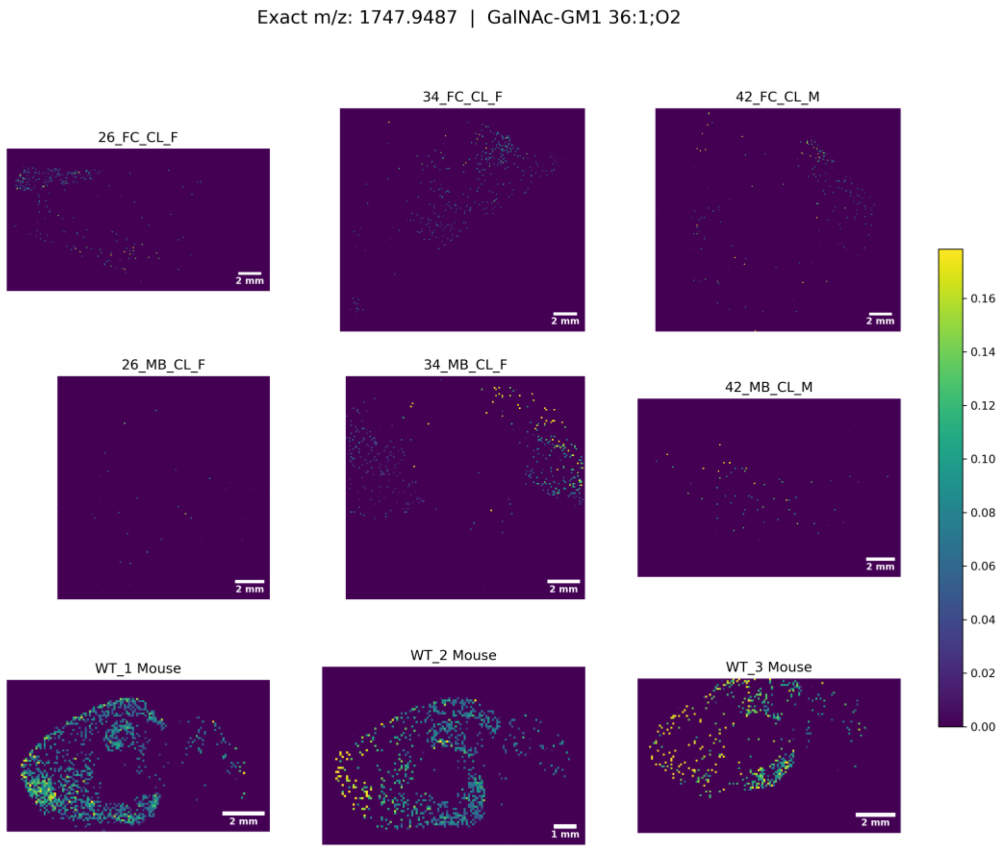
Exact m/z: 1747.9487 | GalNAc-GM1 36:1;O2
26_FC_CL_F
34_FC_CL_F
42_FC_CL_M
26_MB_CL_F
34_MB_CL_F
42_MB_CL_M
WT_1 Mouse
WT_2 Mouse
WT_3 Mouse
2 mm
1 mm
0.16
0.14
0.12
0.10
0.08
0.06
0.04
0.02
0.00

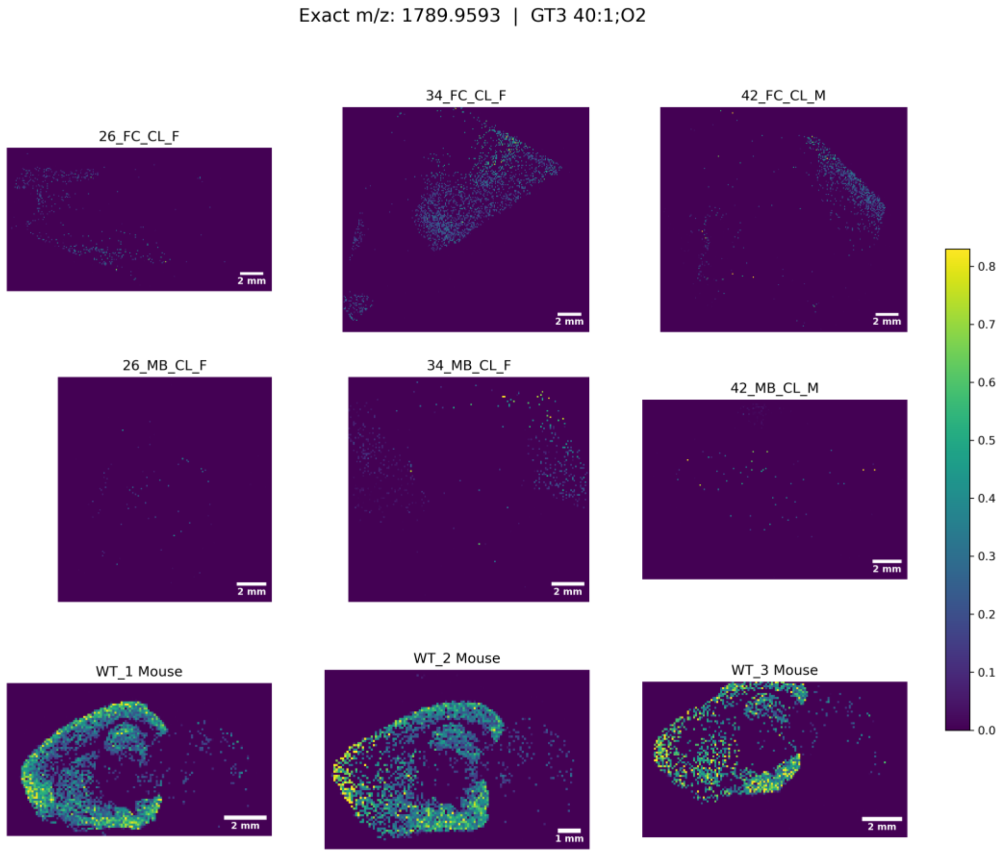
Exact m/z: 1789.9593 | GT3 40:1;O2
26_FC_CL_F
34_FC_CL_F
42_FC_CL_M
26_MB_CL_F
34_MB_CL_F
42_MB_CL_M
WT_1 Mouse
WT_2 Mouse
WT_3 Mouse
2 mm
1 mm
0.8
0.7
0.6
0.5
0.4
0.3
0.2
0.1
0.0

Exact m/z: 1791.975 | GalNAcBeta Cer 38:1;O2

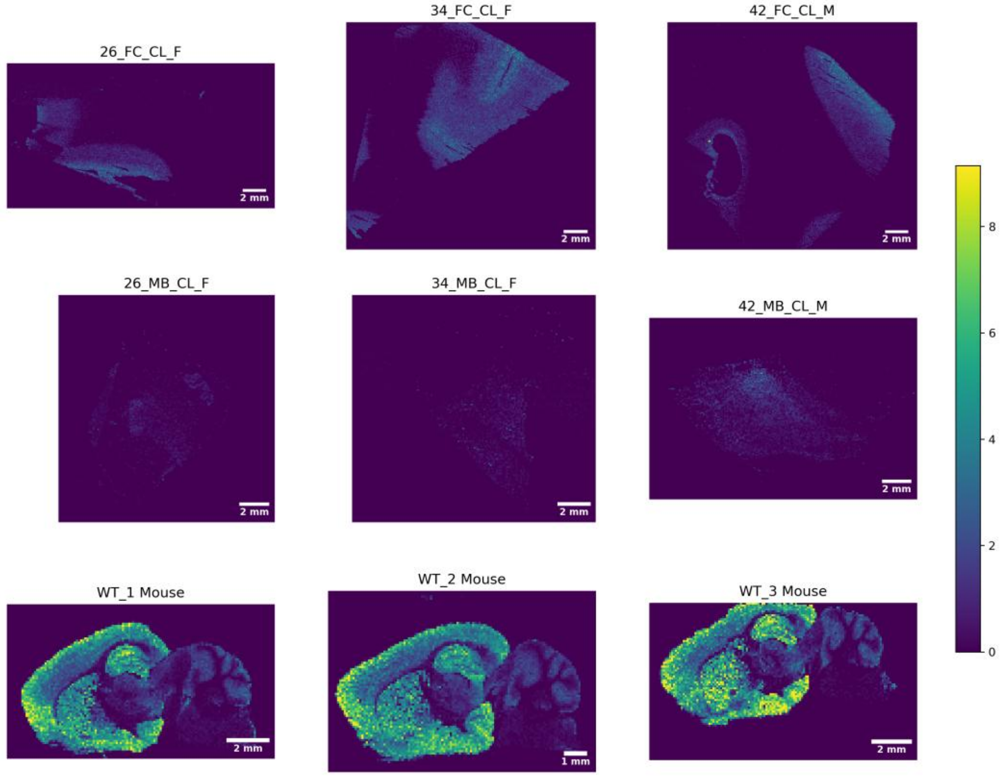


Exact m/z: 1820.0063 | GalNAcBeta Cer 40:1;O2

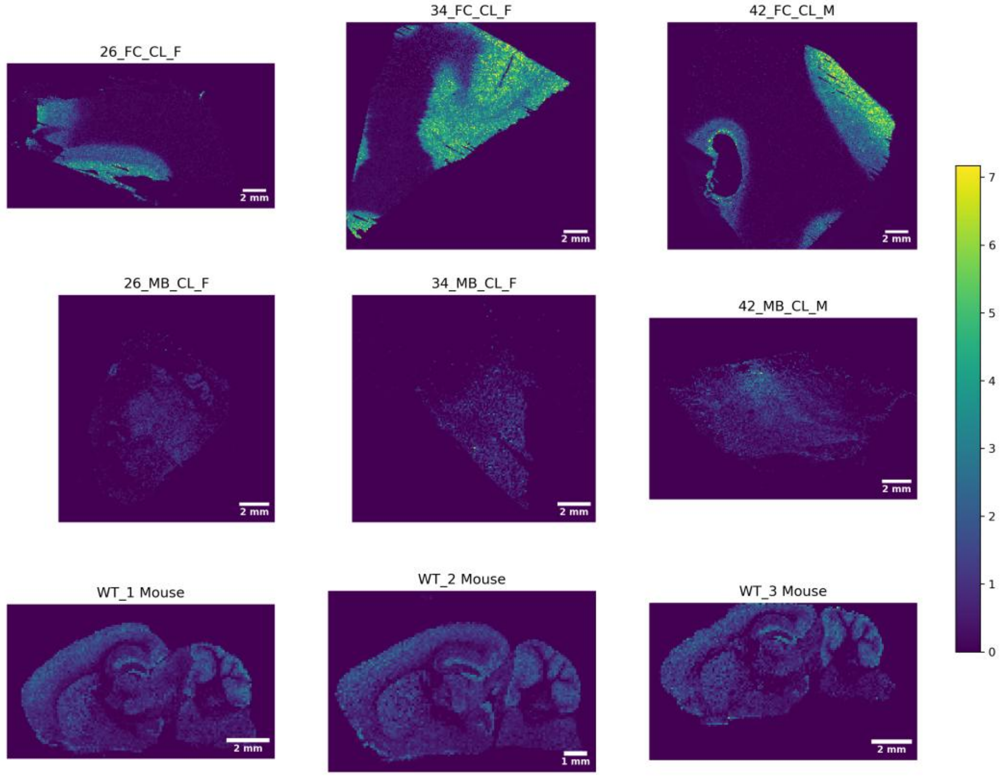

Exact m/z: 1831.9699 | O-Ac-GT3 38:1;O2

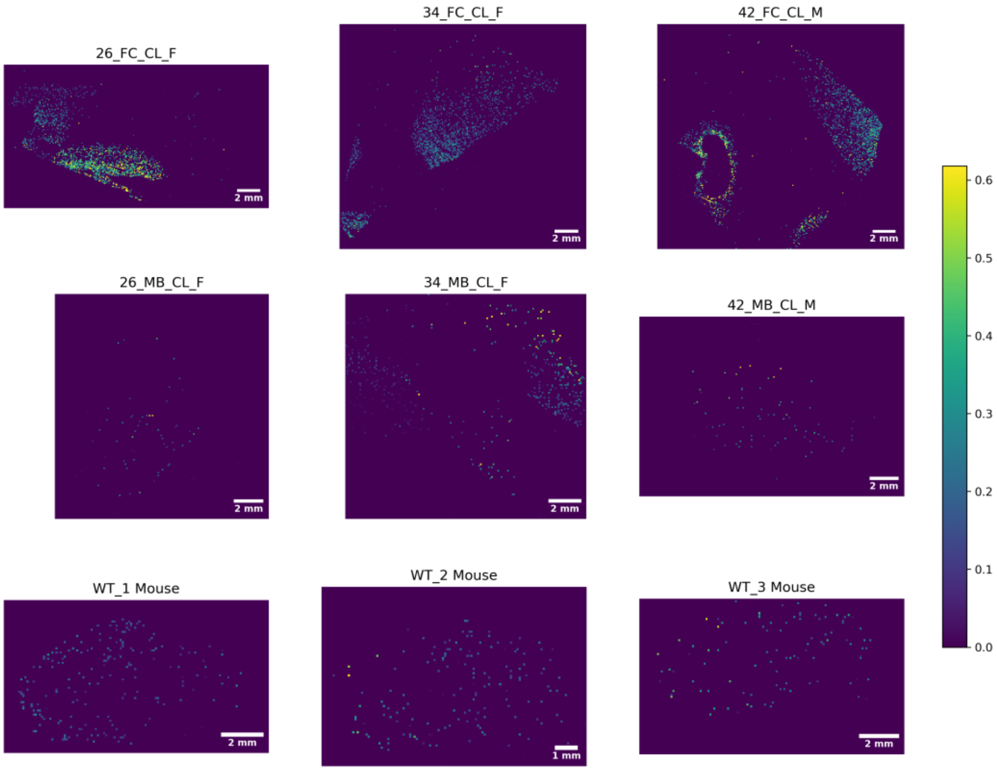


Exact m/z: 1833.9491 | GD1 36:2;O2

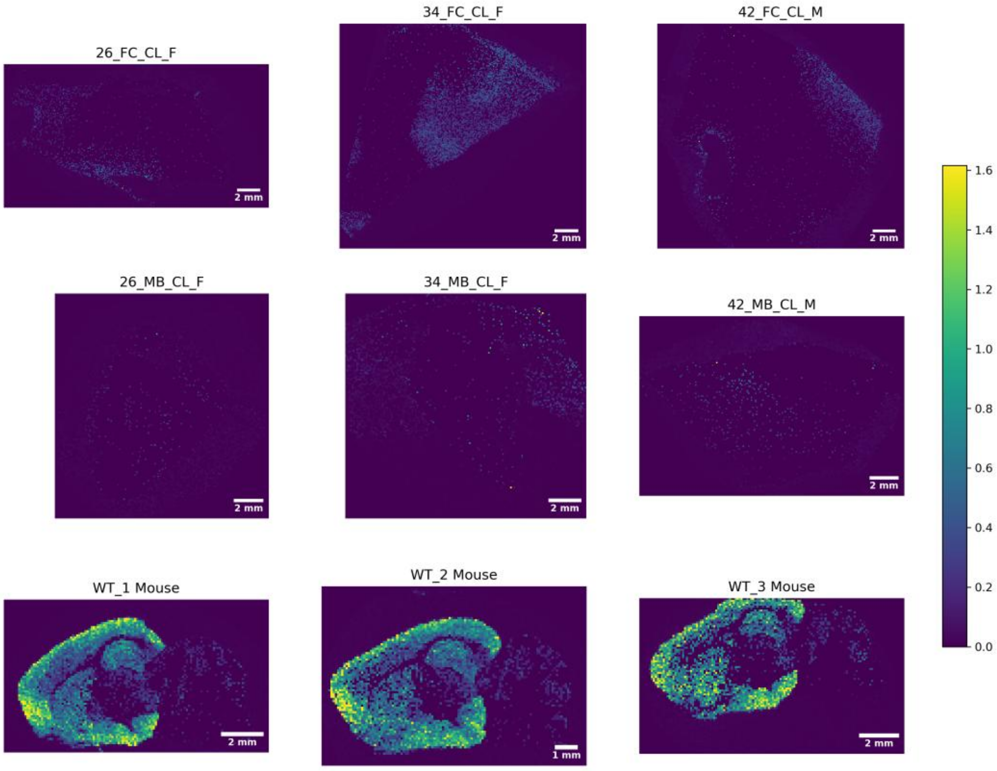

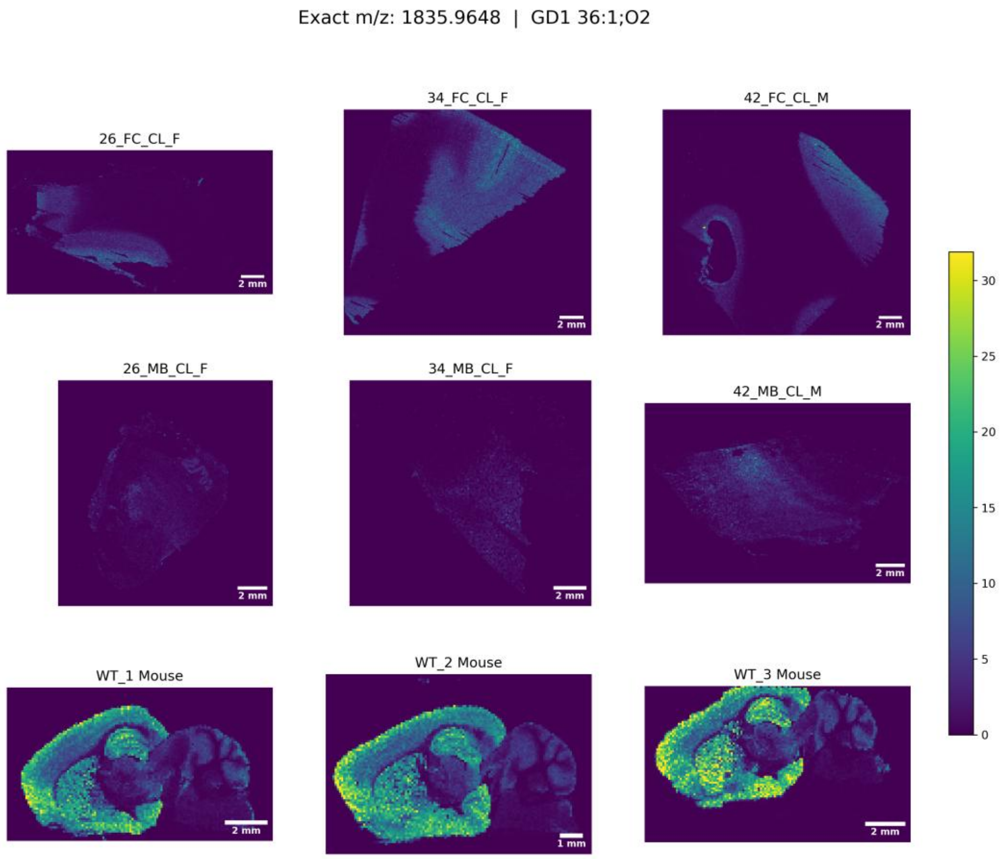
Exact m/z: 1835.9648 | GD1 36:1;O2
26_FC_CL_F
34_FC_CL_F
42_FC_CL_M
26_MB_CL_F
34_MB_CL_F
42_MB_CL_M
WT_1 Mouse
WT_2 Mouse
WT_3 Mouse
2 mm
1 mm
0
5
10
15
20
25
30

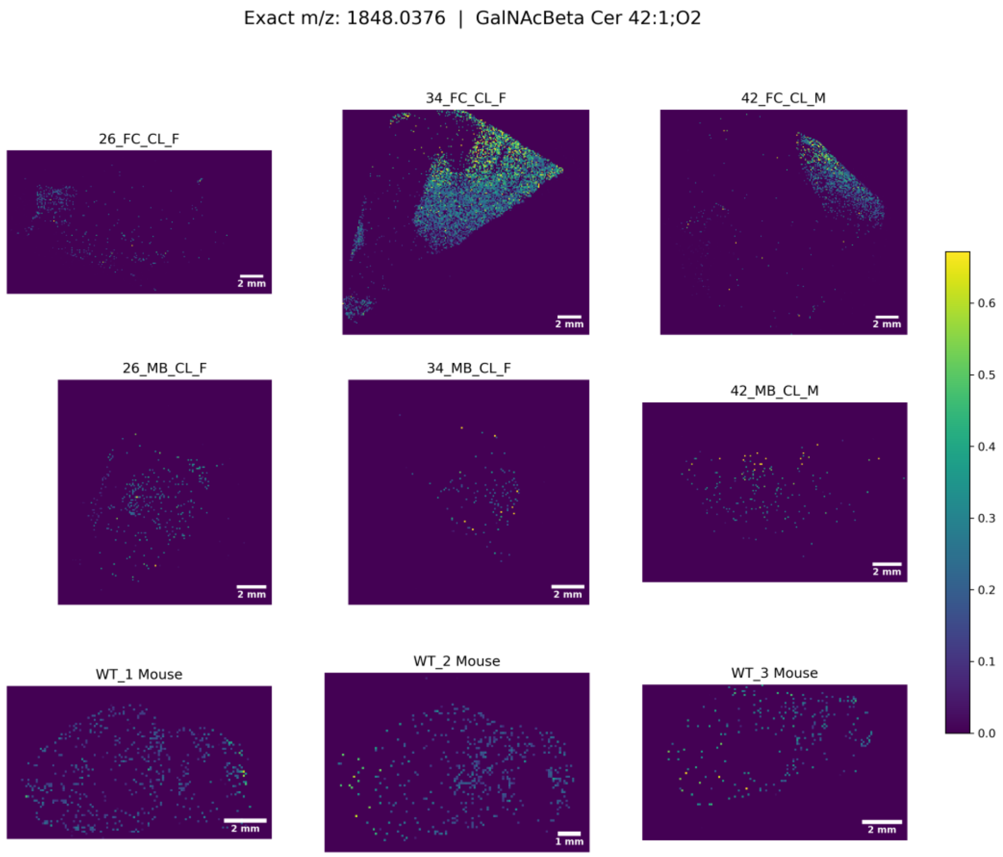
Exact m/z: 1848.0376 | GalNAcBeta Cer 42:1;O2
26_FC_CL_F
34_FC_CL_F
42_FC_CL_M
26_MB_CL_F
34_MB_CL_F
42_MB_CL_M
WT_1 Mouse
WT_2 Mouse
WT_3 Mouse
2 mm
1 mm
0.0
0.1
0.2
0.3
0.4
0.5
0.6

Exact m/z: 1849.9804 | GD1 37:1;O2

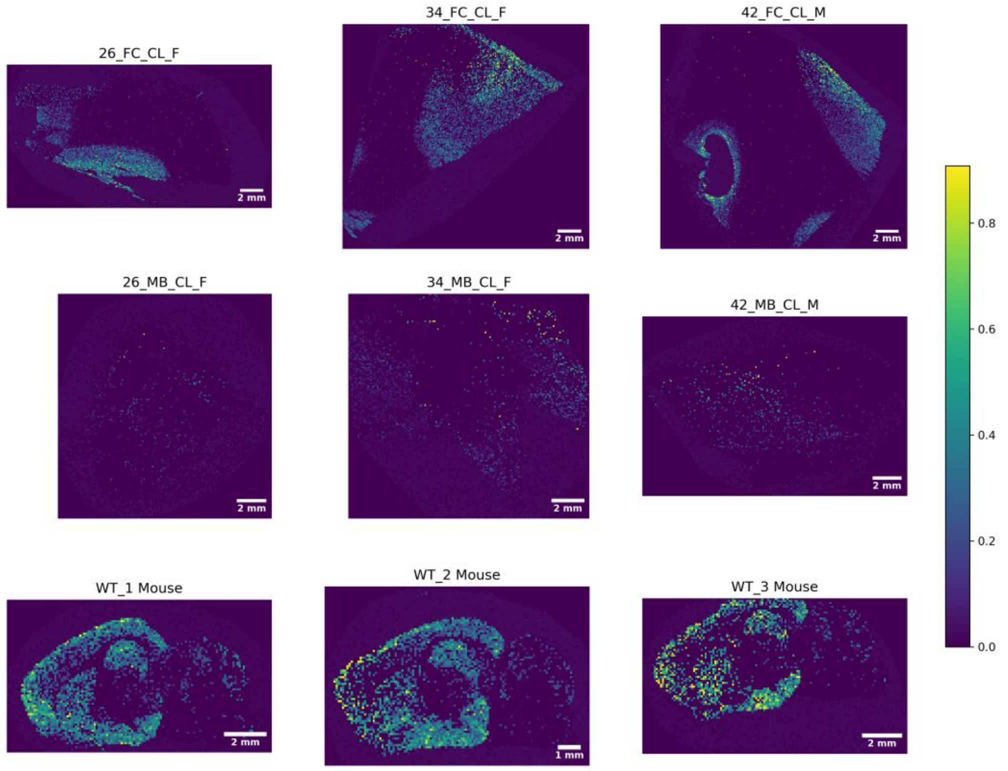


Exact m/z: 1861.9804 | GD1 38:2;O2

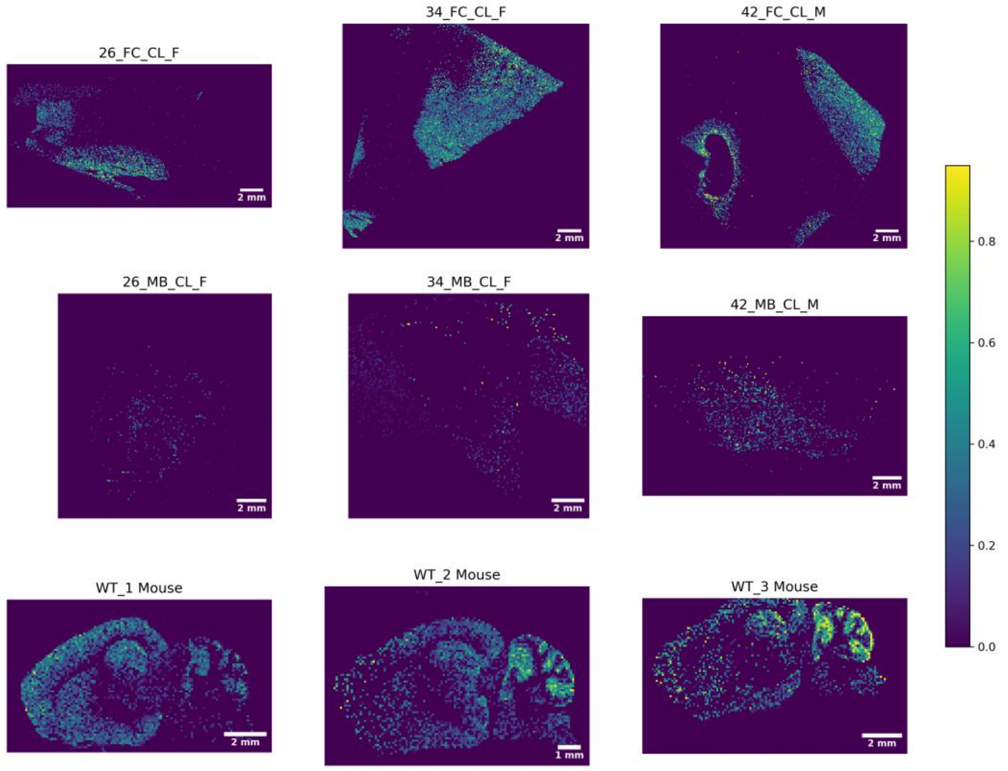

Exact m/z: 1863.9961 | GD1 38:1;O2

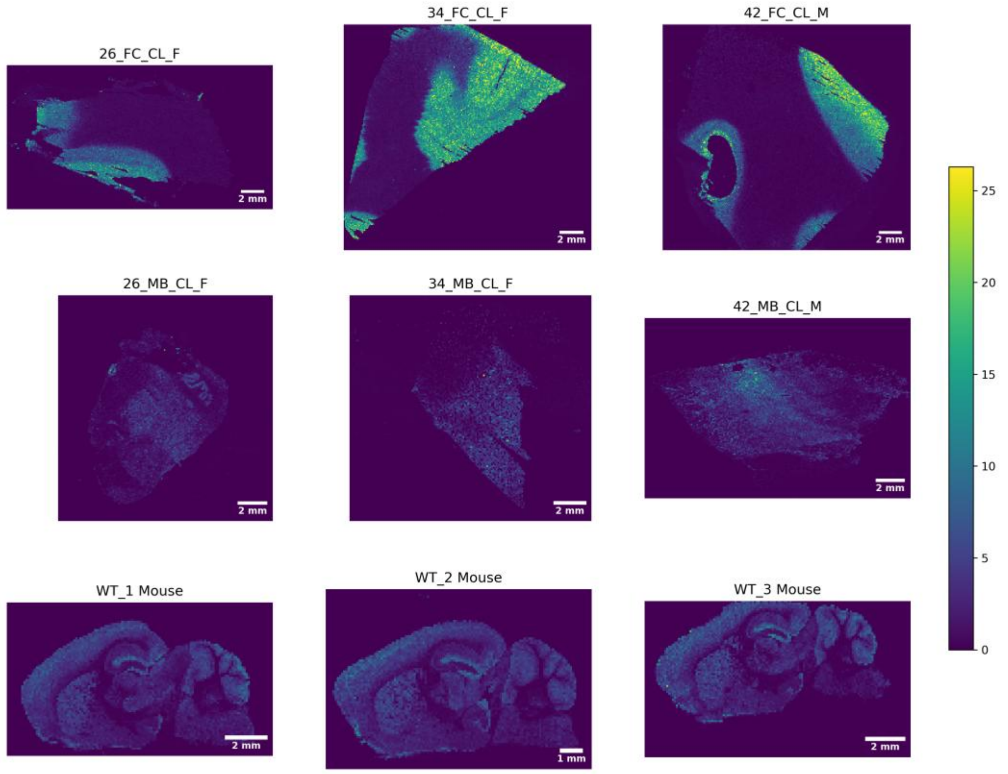


Exact m/z: 1875.9961 | GD1 39:2;O2

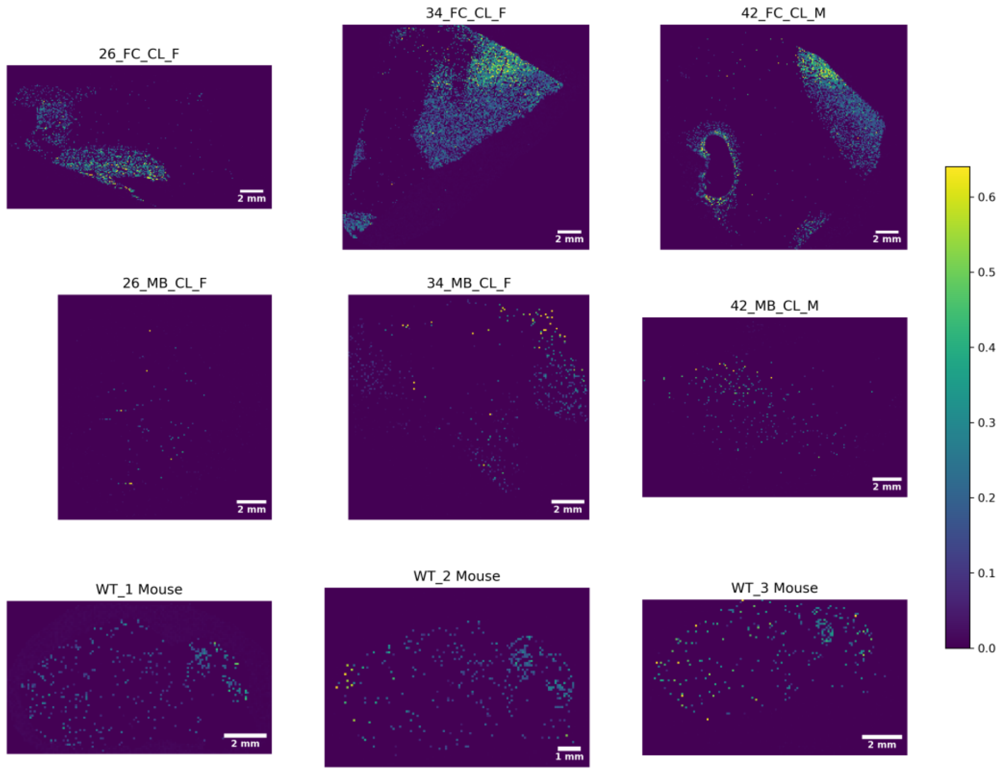

Exact m/z: 1877.9753 | O-Ac-GD1 36:1;O2

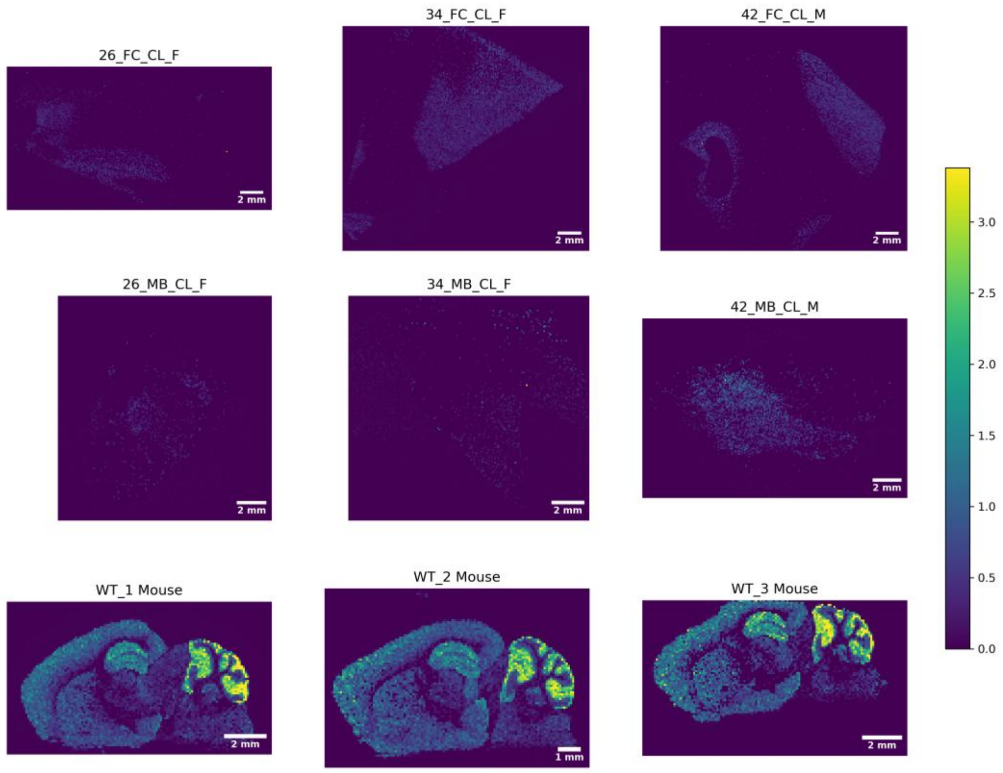


Exact m/z: 1878.0117 | GD1 39:1;O2

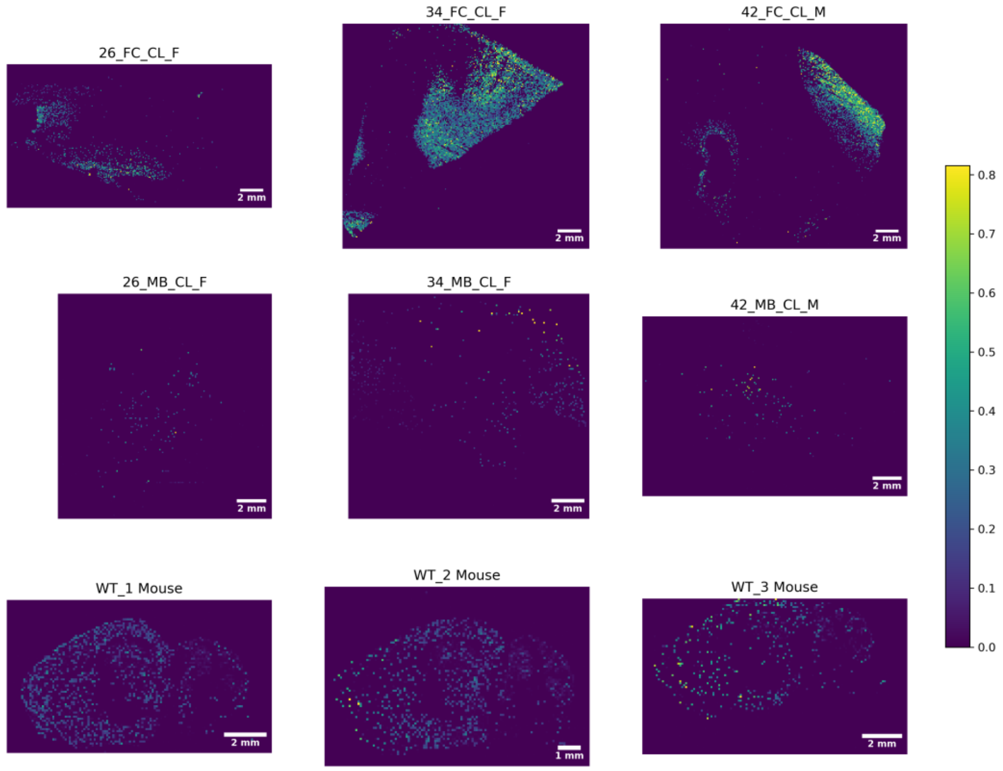

Exact m/z: 1892.0274 | GD1 40:1;O2

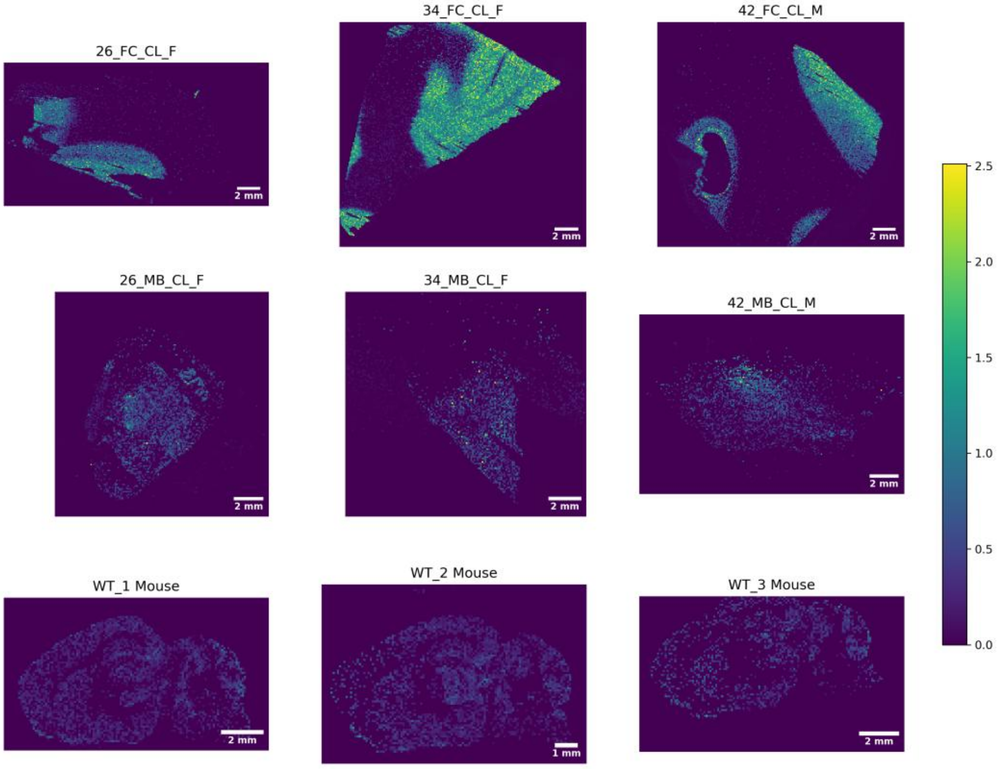


Exact m/z: 1906.0067 | O-Ac-GD1 38:1;O2

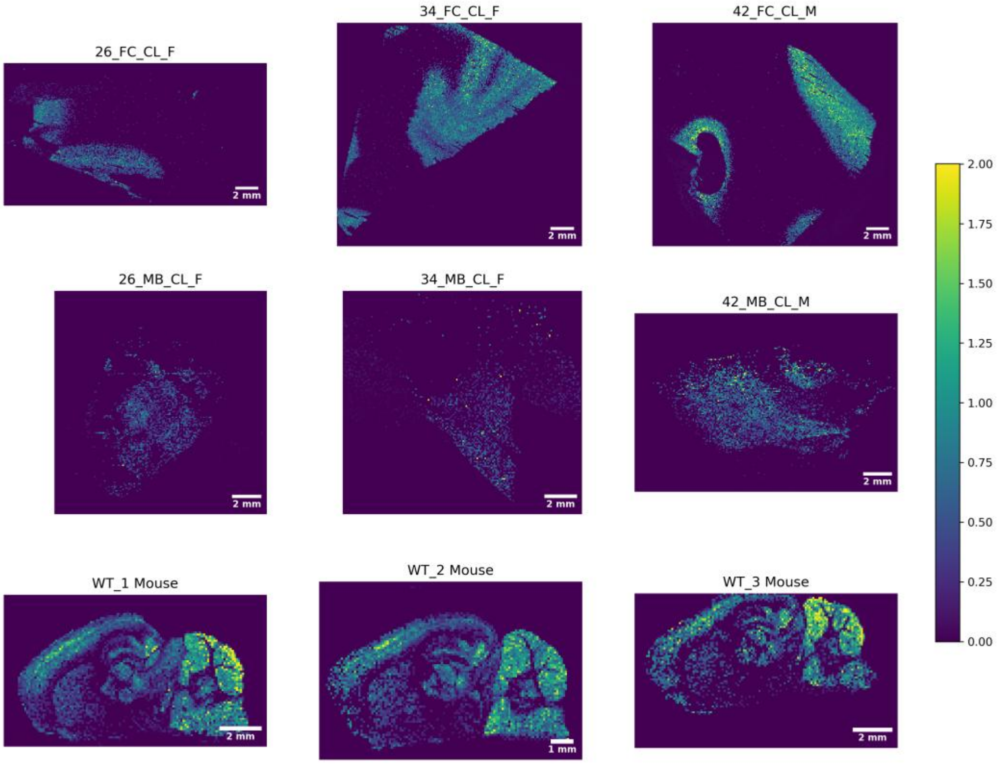

Exact m/z: 2067.0755 | GalNAc-GD1 38:1;O2

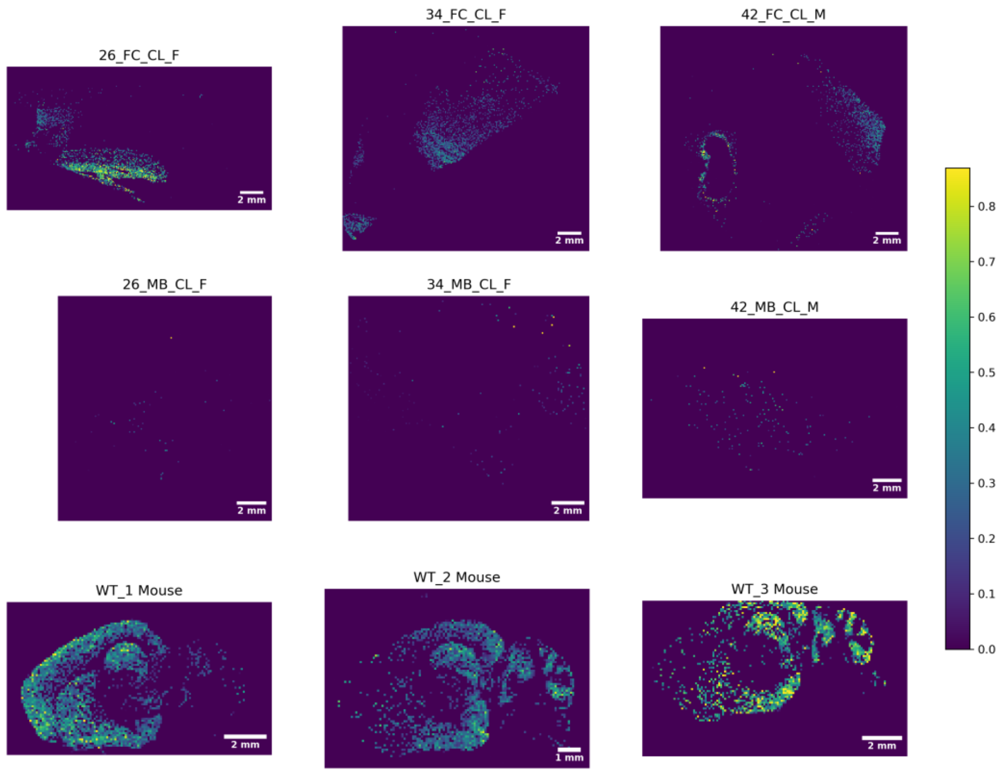


Exact m/z: 2083.0704 | GalNAc-GD1 Neu5Ac/Neu5Gc 38:1;O2

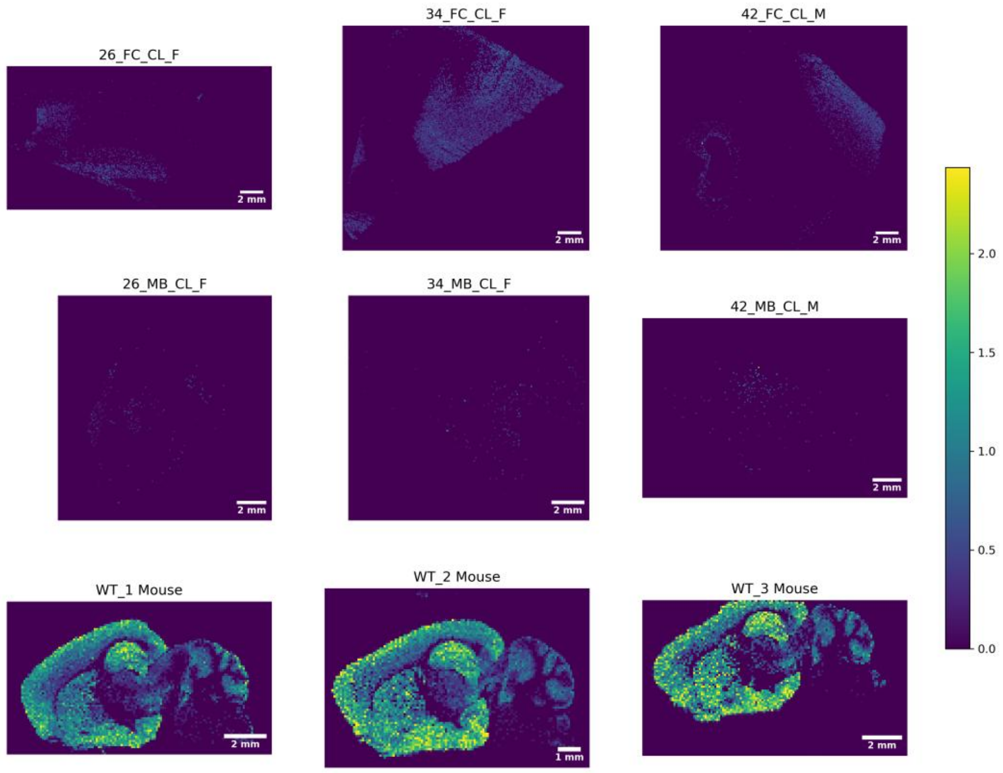

Exact m/z: 2095.1068 | GalNAc-GD1-OH 40:1;O2

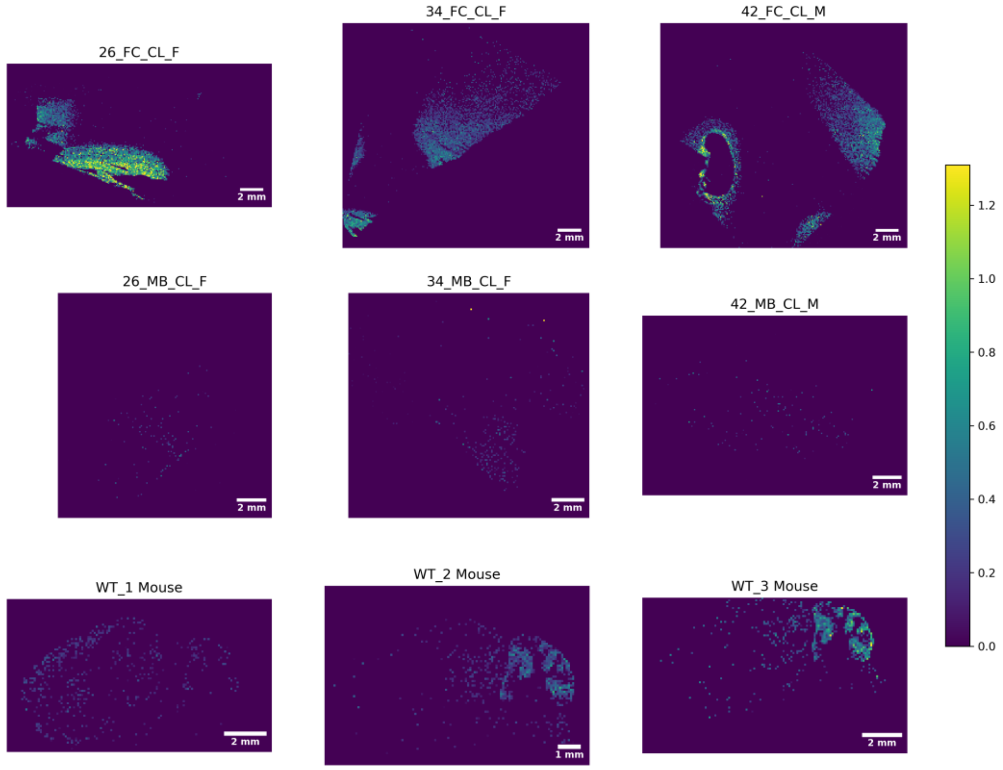


Exact m/z: 2111.1017 | GalNAc-GD1 Neu5Ac/Neu5Gc 40:1;O2

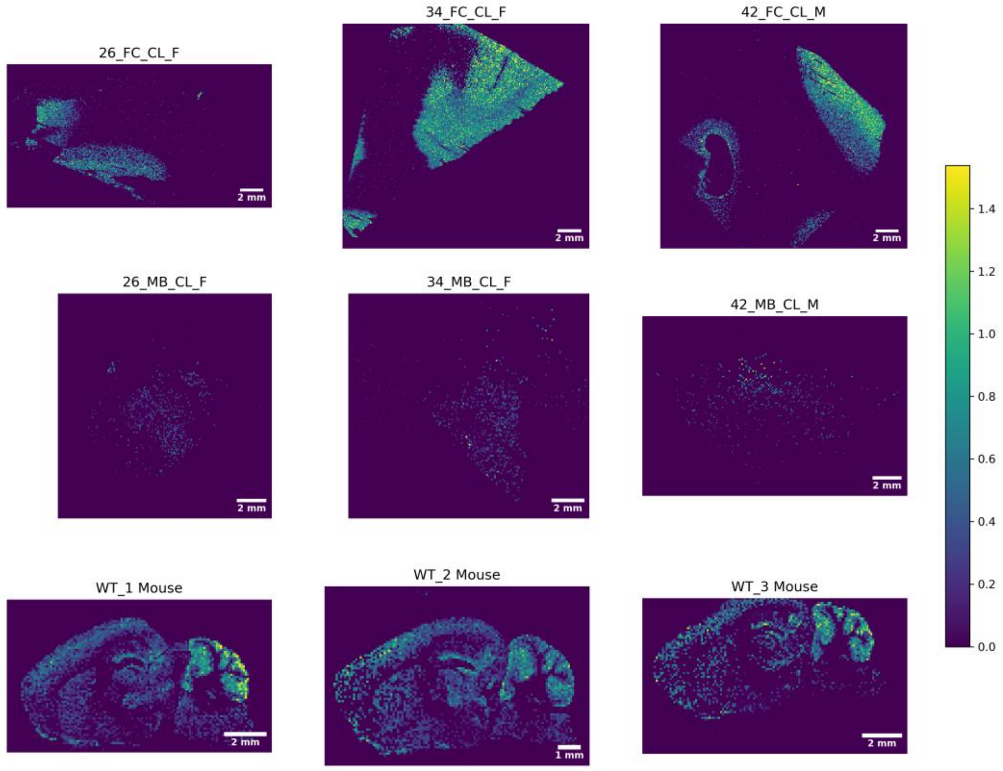

Exact m/z: 2127.0602 | GT1 36:1;O2

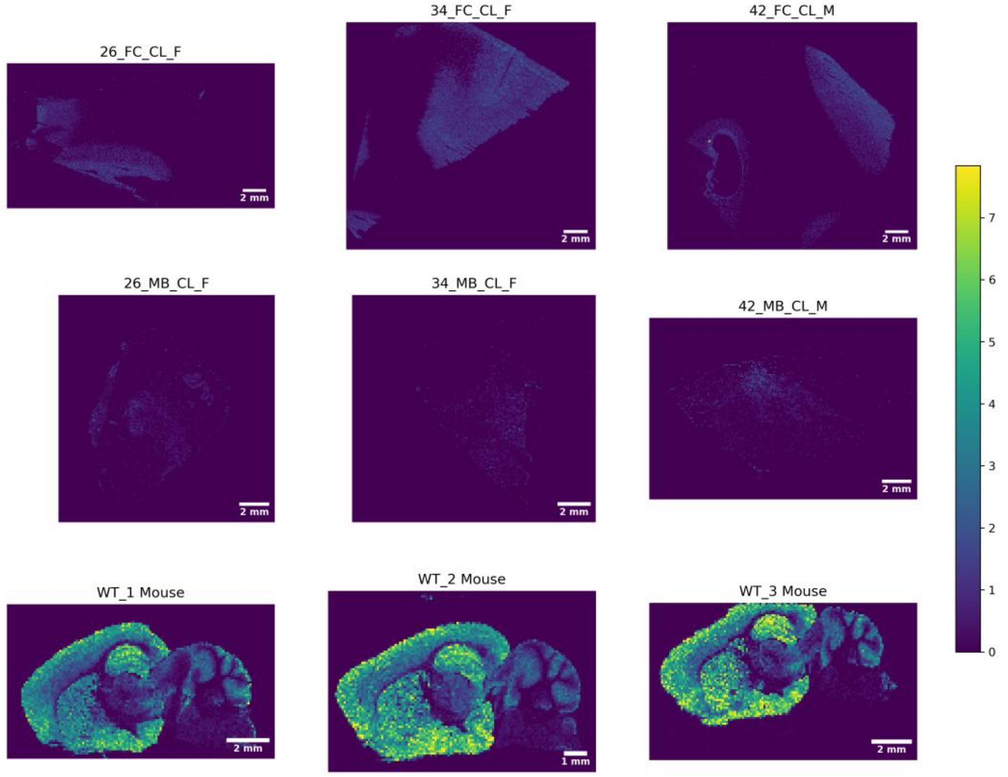


Exact m/z: 2129.0759 | NeuGc Cer 36:1;O2

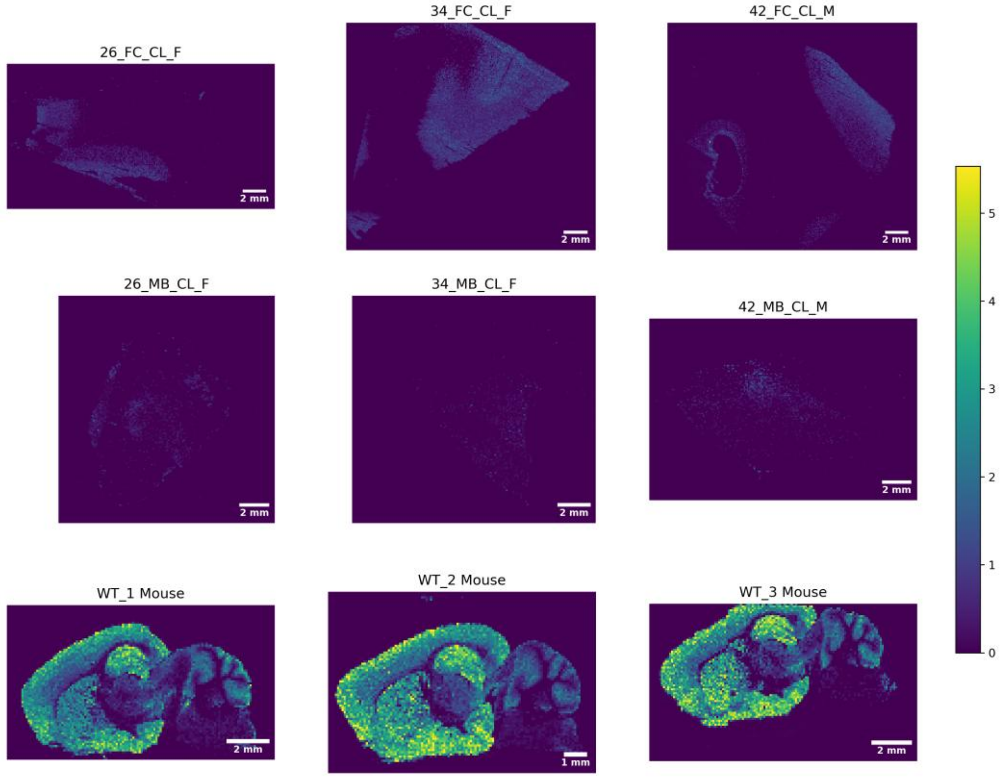

Exact m/z: 2132.0392 | GalAlpha Cer 34:1;O2

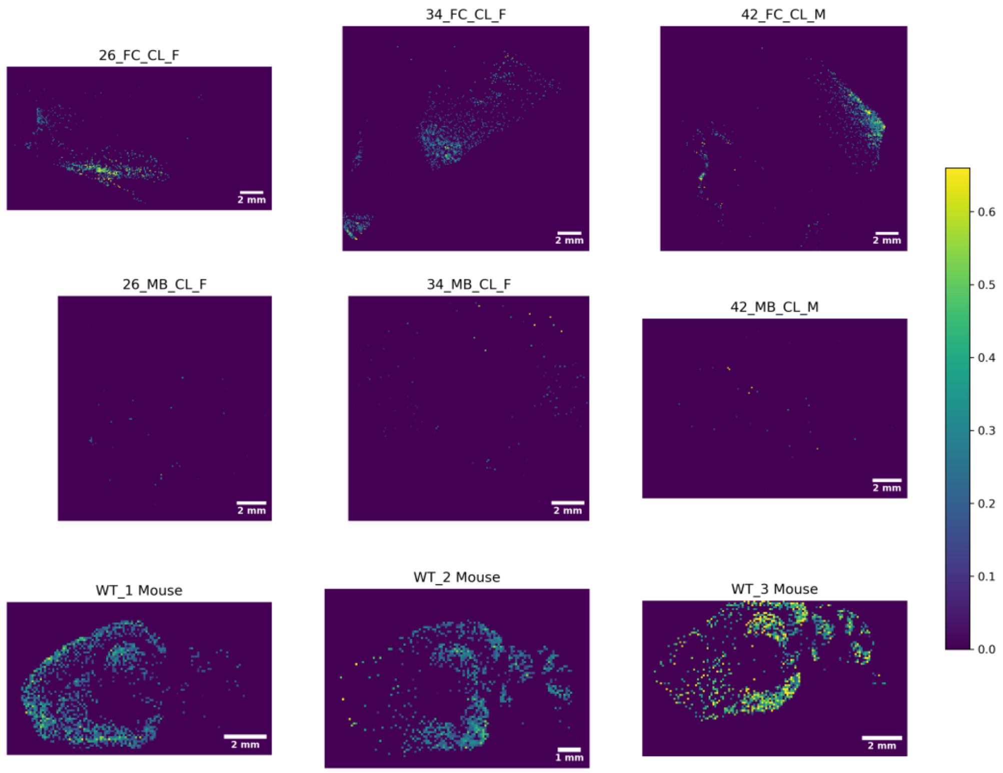


Exact m/z: 2141.0759 | GT1 37:1;O2

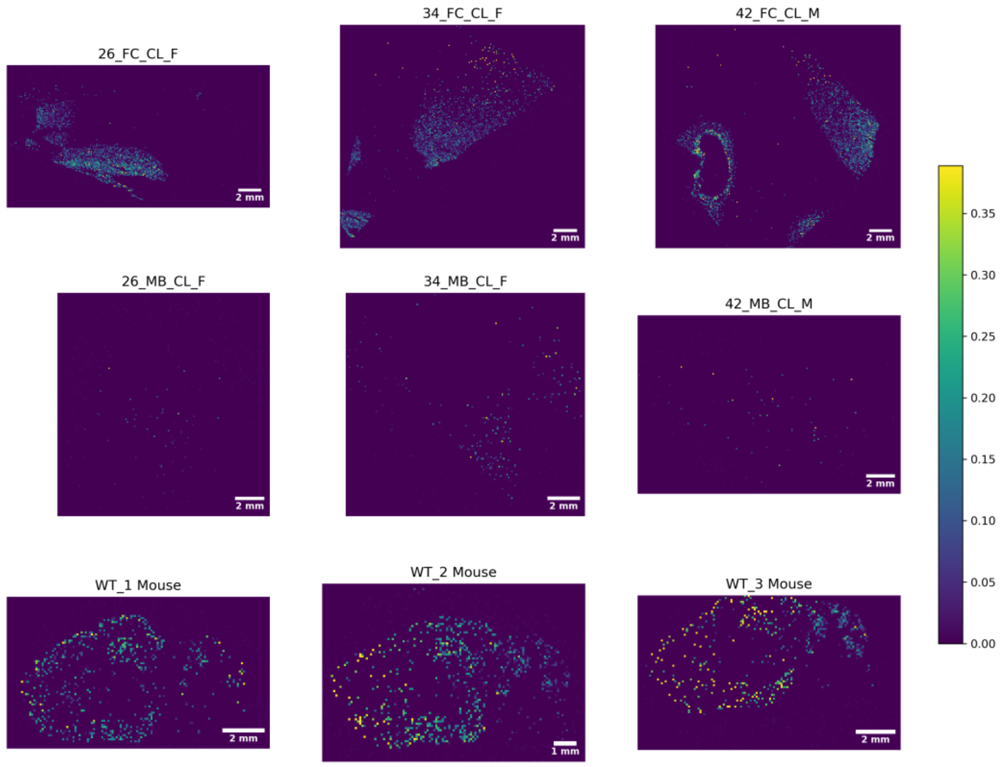

Exact m/z: 2149.0446 | GT1 38:4;O2

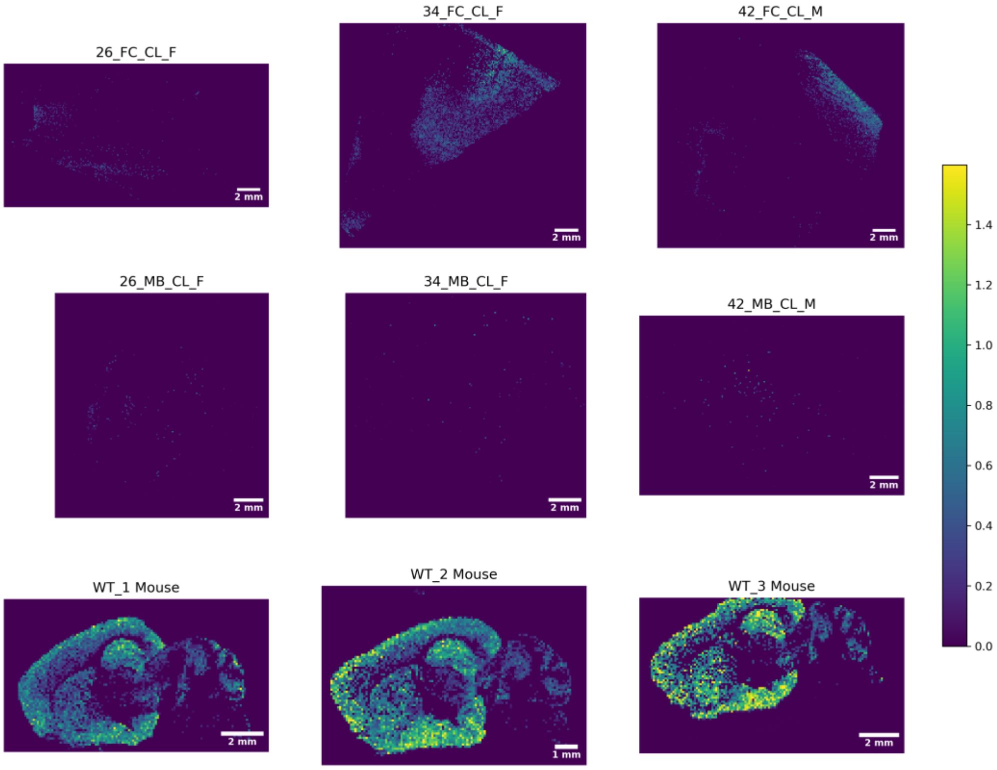


Exact m/z: 2151.0602 | GT1 38:3;O2

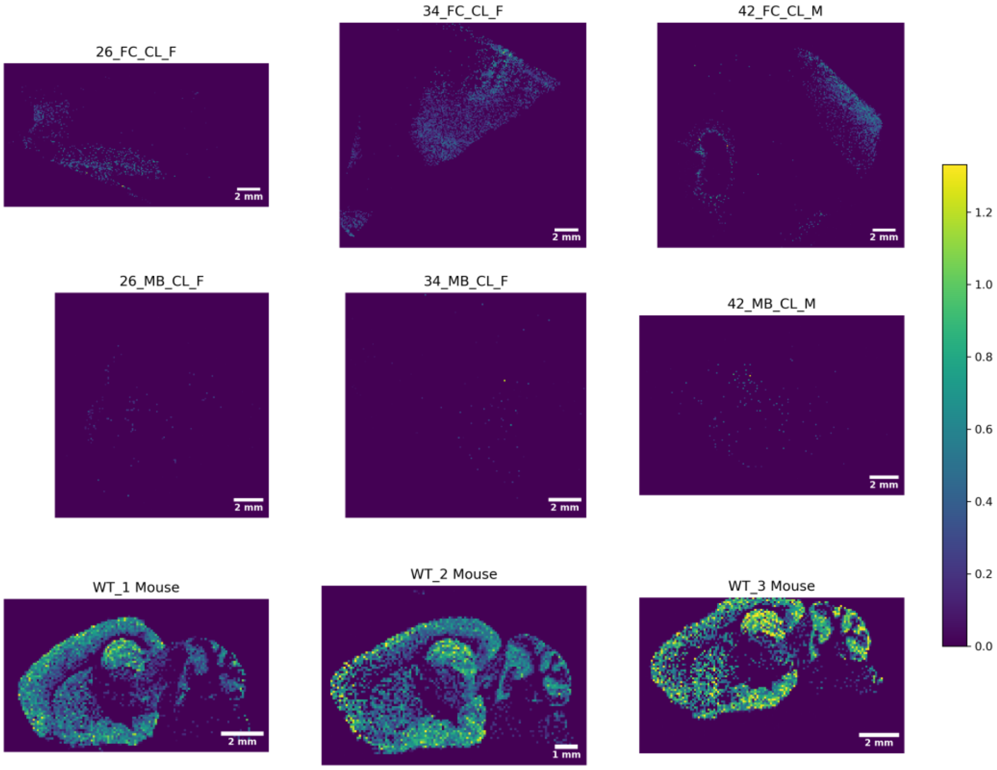

Exact m/z: 2155.0915 | GT1 38:1;O2

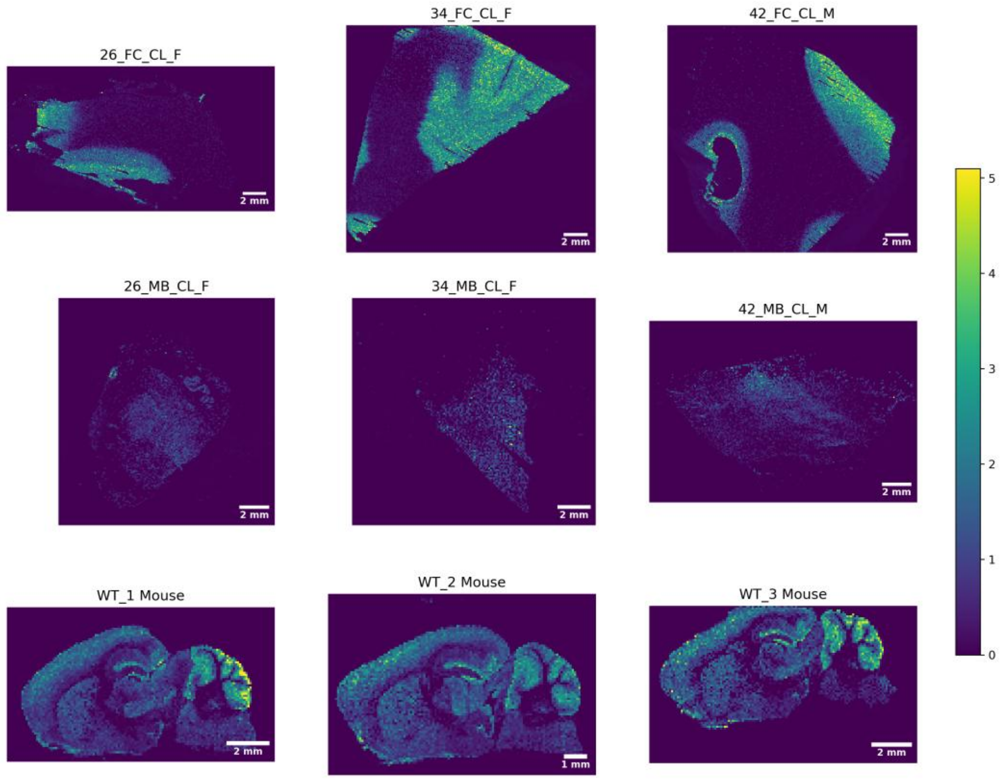


Exact m/z: 2157.1072 | NeuGc Cer 38:1;O2

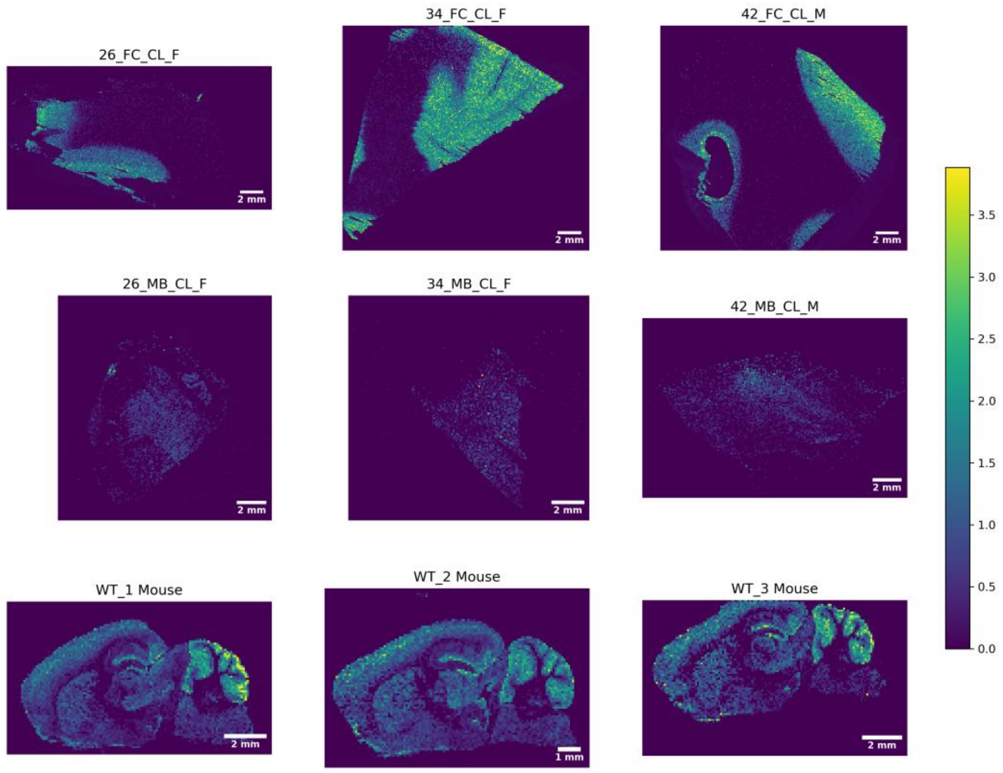

Exact m/z: 2160.0705 | GalAlpha Cer 36:1;O2

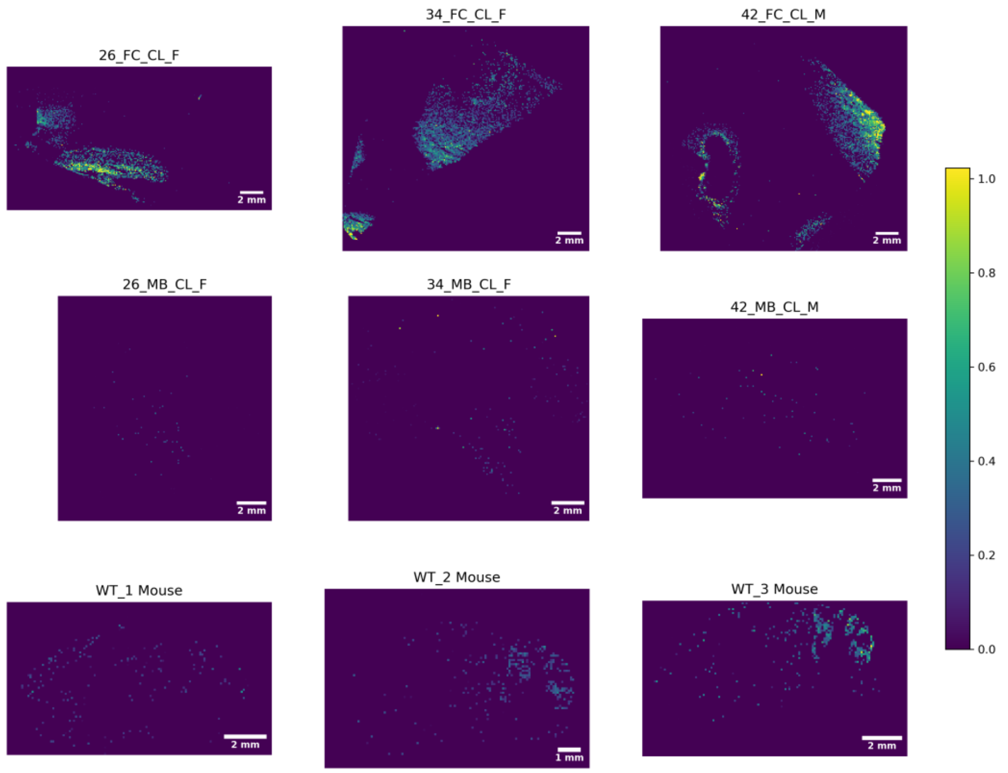


Exact m/z: 2169.0708 | O-Ac-GT1 36:1;O2

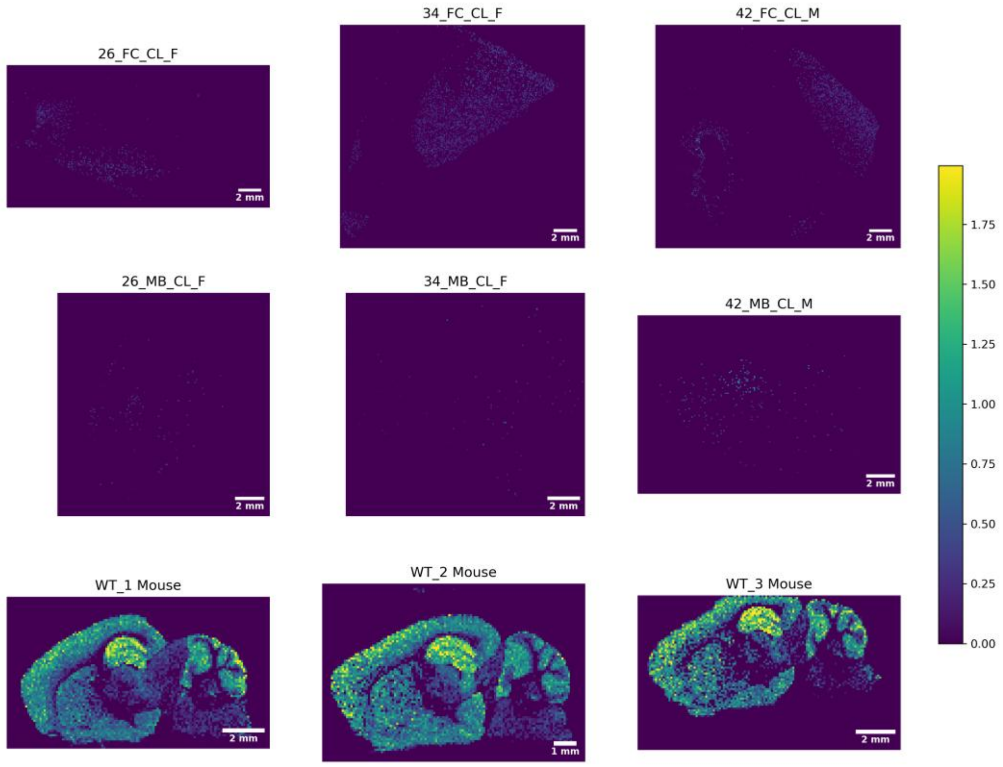

Exact m/z: 2171.0864 | GT1-OH 38:1;O2

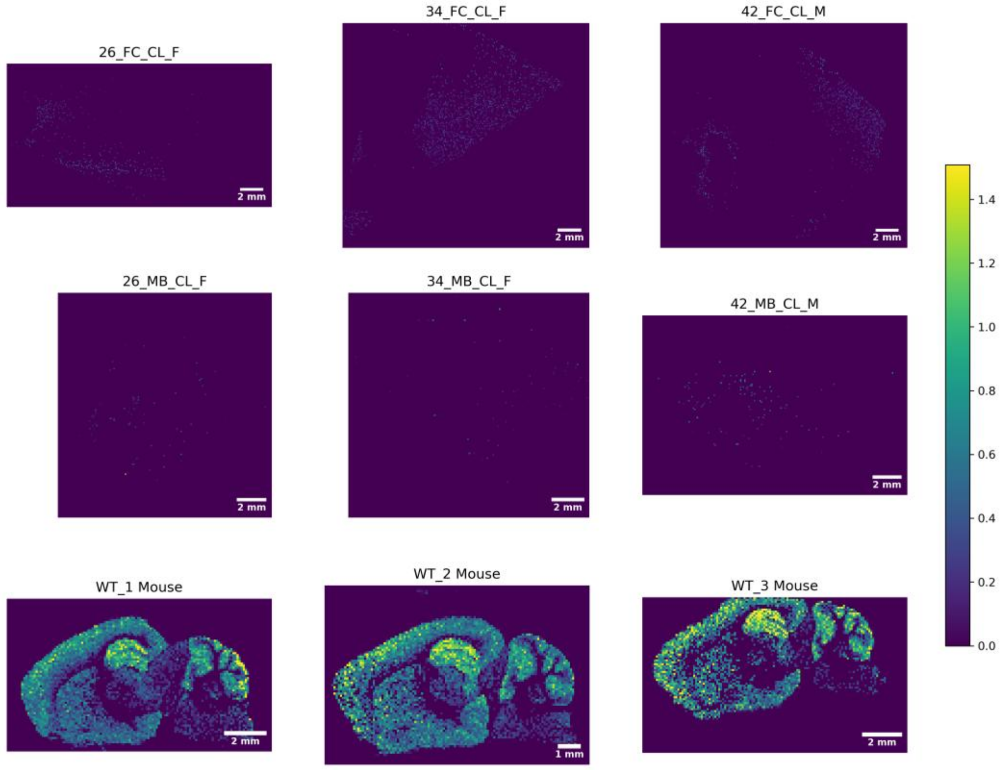


Exact m/z: 2177.0759 | GT1 40:4;O2

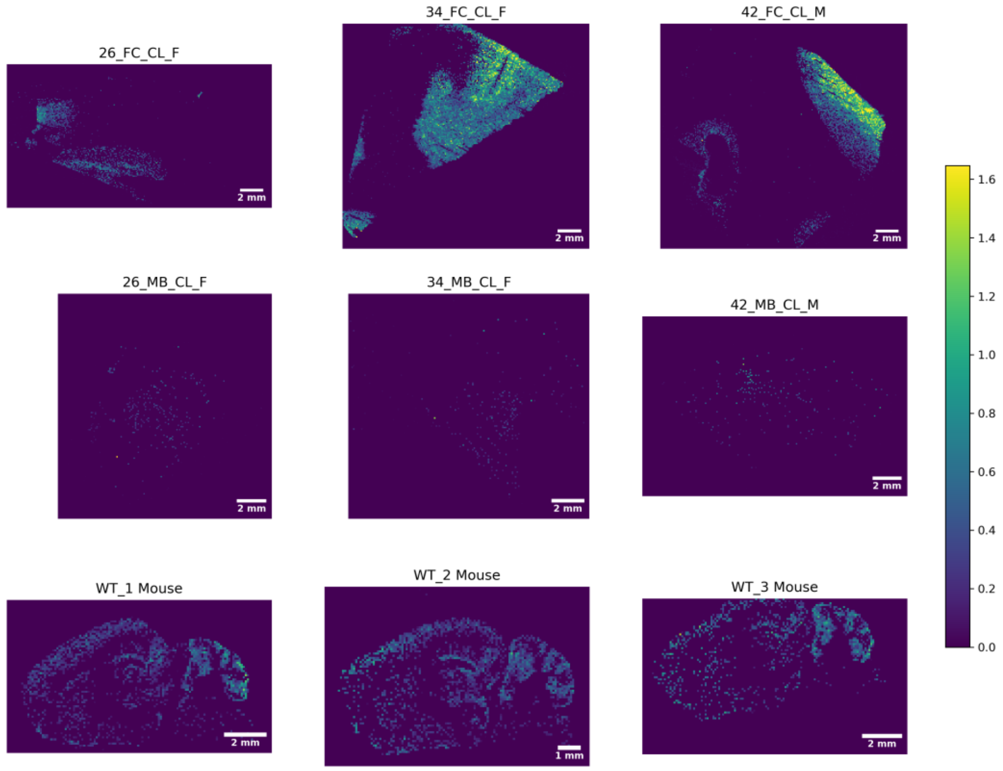

Exact m/z: 2183.1228 | GT1 40:1;O2

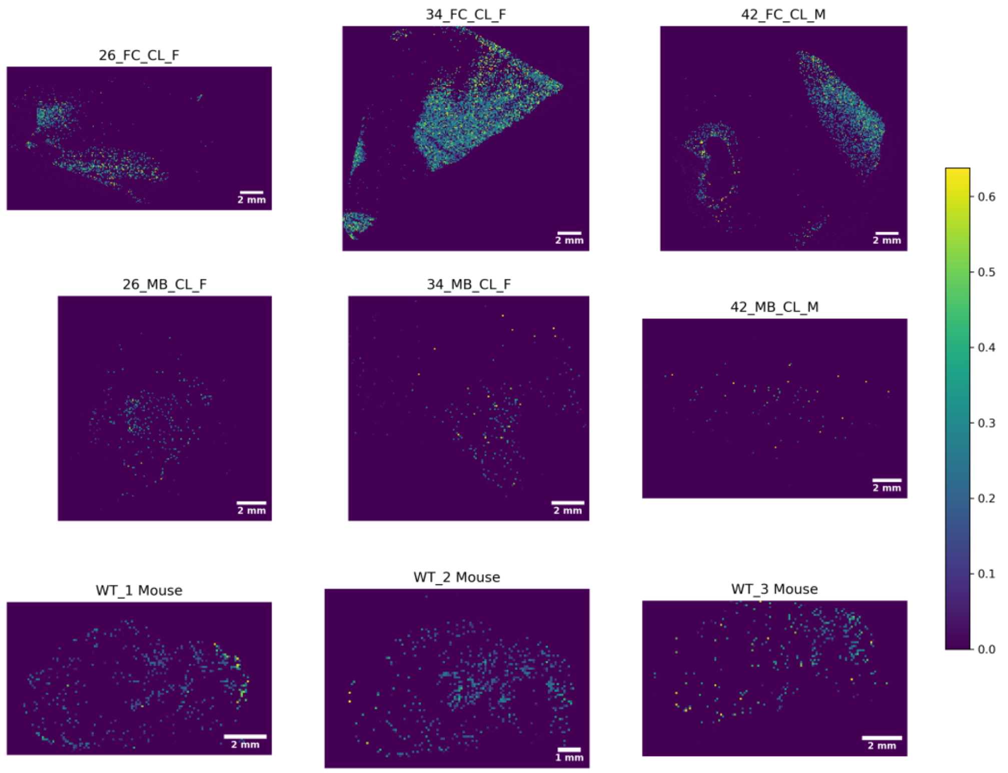


Exact m/z: 2197.1021 | O-Ac-GT1 38:1;O2

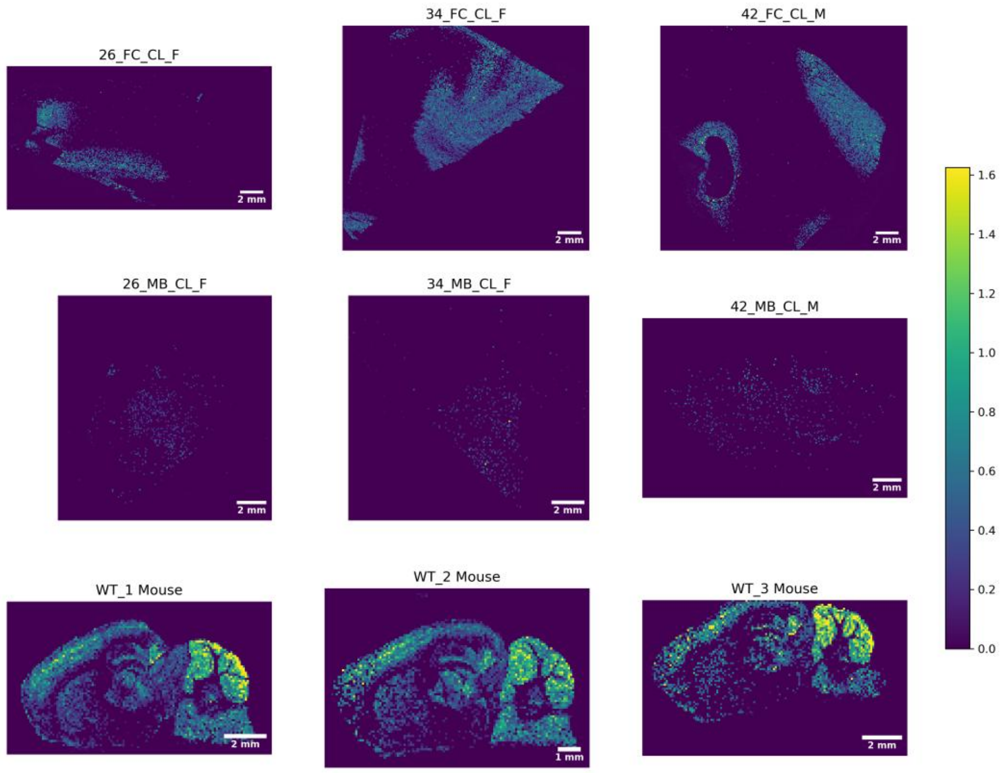

Exact m/z: 2418.1556 | GQ1 36:1;O2

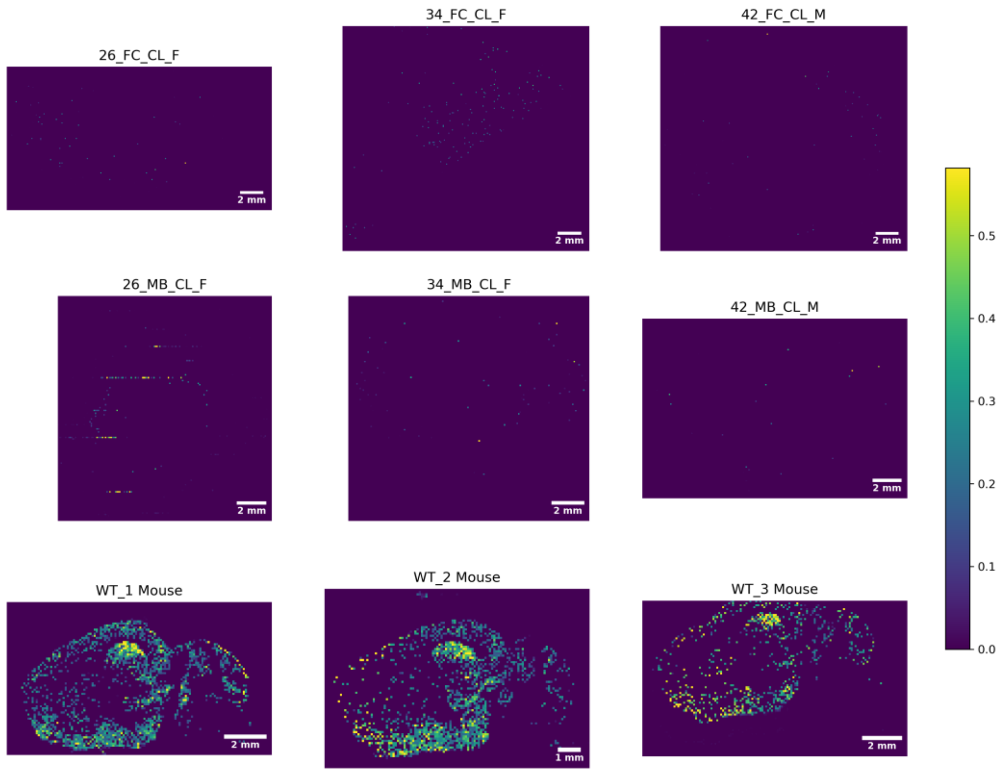


Exact m/z: 2446.1869 | GQ1 38:1;O2

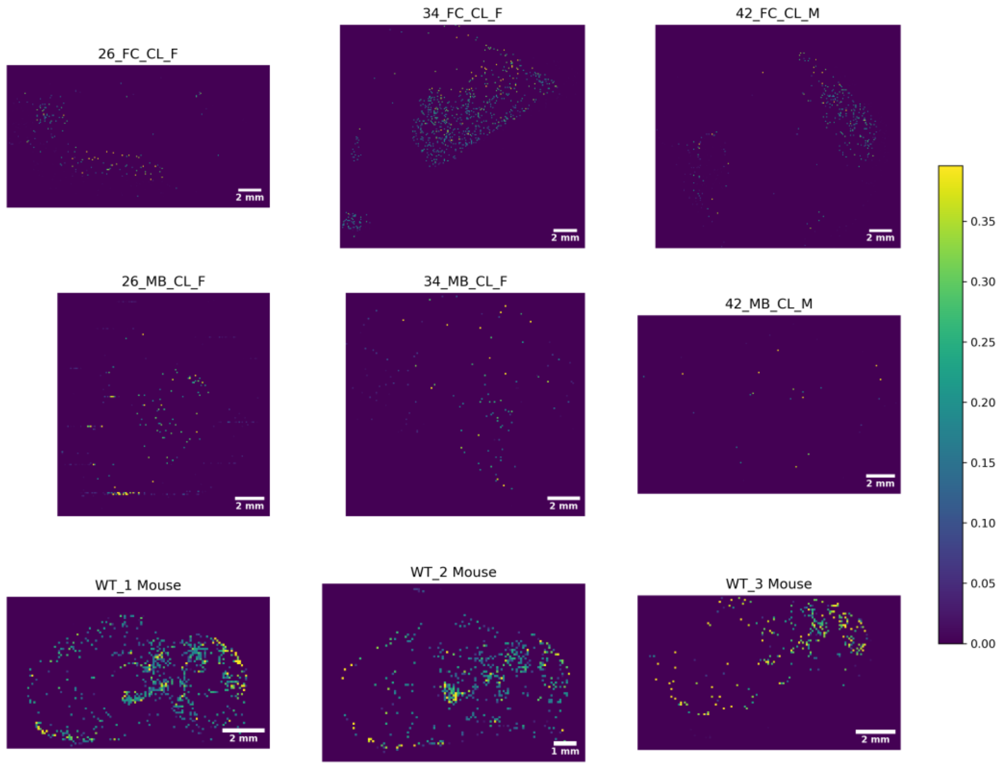

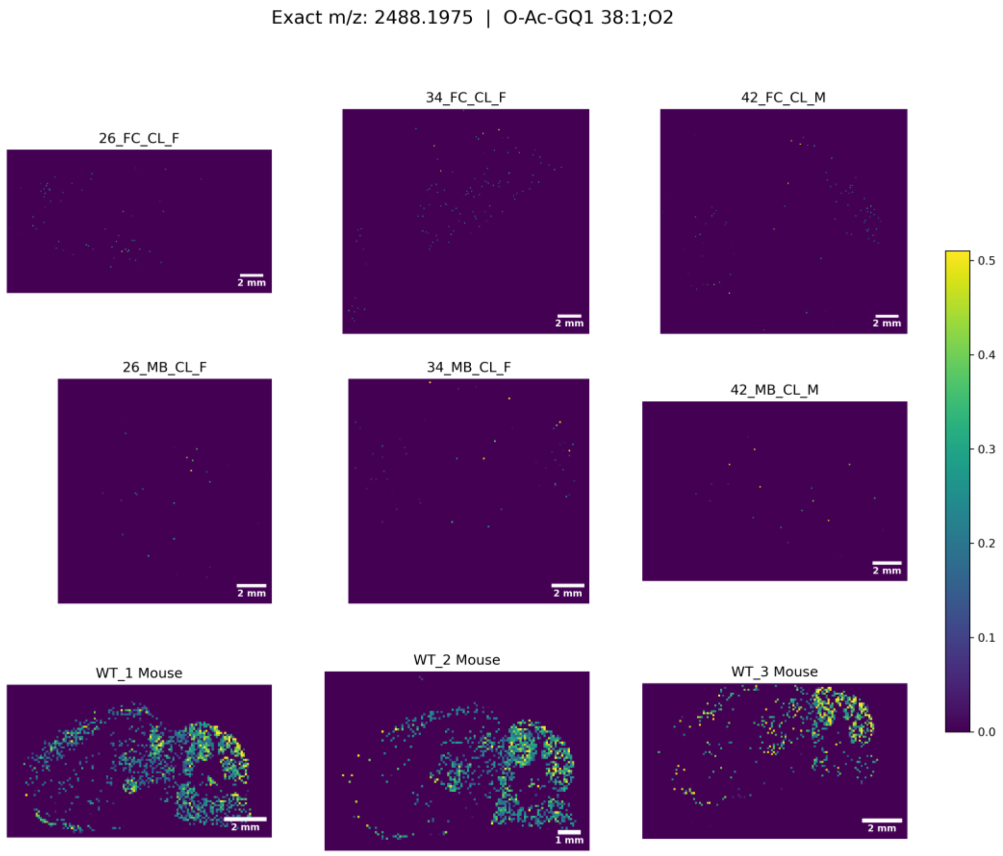
Exact m/z: 2488.1975 | O-Ac-GQ1 38:1;O2
26_FC_CL_F
34_FC_CL_F
42_FC_CL_M
26_MB_CL_F
34_MB_CL_F
42_MB_CL_M
WT_1 Mouse
WT_2 Mouse
WT_3 Mouse
2 mm
1 mm
0.5
0.4
0.3
0.2
0.1
0.0
